# Studying stochastic systems biology of the cell with single-cell genomics data

**DOI:** 10.1101/2023.05.17.541250

**Authors:** Gennady Gorin, John J. Vastola, Lior Pachter

## Abstract

Recent experimental developments in genome-wide RNA quantification hold considerable promise for systems biology. However, rigorously probing the biology of living cells requires a unified mathematical framework that accounts for single-molecule biological stochasticity in the context of technical variation associated with genomics assays. We review models for a variety of RNA transcription processes, as well as the encapsulation and library construction steps of microfluidics-based single-cell RNA sequencing, and present a framework to integrate these phenomena by the manipulation of generating functions. Finally, we use simulated scenarios and biological data to illustrate the implications and applications of the approach.

## 1 Introduction

In his classic systems biology textbook^1^, D. J. Wilkinson notes that “Improvements in experimental technology are enabling quantitative real-time imaging of expression at the single-cell level, and improvement in computing technology is allowing modelling and stochastic simulation of such systems at levels of detail previously impossible. The message that keeps being repeated is that the kinetics of biological processes at the intra-cellular level are stochastic, and that cellular function cannot be properly understood without building that stochasticity into *in silico* models”. From this perspective, systems biology studies control over randomness, and the ways in which living cells exploit variability to grow and function. Counterintuitively, this stochastic *weltanschauung* relies on mental models that are inherently deterministic: differentiation landscapes^2–6^, gene expression manifolds^7^, cellular state graphs^8,9^, gene regulatory networks^10,11^, and kinetic parameters^12^. Analysis of experimental data therefore requires reconciling underlying deterministic structure with biological stochasticity and experimental technical variability, or noise. In particular, distinguishing technical noise from biological stochasticity involves the statistical modeling of experimental readouts, expected noise sources, and the signal-to-noise ratio, and requires consideration of the theoretical and computational tractability of the model.

How can we model these features—latent deterministic structure, biological stochasticity, and technical noise—in a way that balances our models’ ability to adequately describe available data with our own ability to adequately understand the mathematical behavior and biological interpretation of our models? Answering this question is particularly challenging in the context of single-cell genomics, where data sets are large and sparse, the signal-to-noise ratio is low, and stochasticity is one of the defining features of the underlying biophysics^13–15^. Here, we explain why many naïve approaches to understanding the stochastic systems biology of single cells fall short, and describe a theoretical framework that can serve as an alternative. Our framework extends recent work on the mechanistic modeling of single-cell RNA count distributions^16–21^, and addresses both how models can be efficiently fit to single-cell data, and what features of the underlying biology we can hope to learn.

After introducing the general framework, we illustrate its consequences through a series of vignettes. In each case, we consider modeling particular aspects of biological and technical noise, and ask: (1) What do our models help us learn about the underlying biology? and (2) What could go wrong if we ignored these features of our data? We find that certain kinds of noise must be carefully modeled, others are poorly identifiable, while others still cannot be identified at all and can be safely ignored.

## 2 SYSTEMS BIOLOGY AND SINGLE-CELL GENOMICS

### 2.1 Standard approaches to systems biology

If an experiment has ample controls and provides a readout with a high signal-to-noise ratio in the relevant variables, coarse-grained, moment-based models can be ideal. For example, investigations of cell growth have effectively used least-squares regression to fit scaling relationships between cell volume and molecular abundance that hold on average^22,23^. Analogously, experiments leveraging the integration of multiple fluorescent reporters have successfully decomposed molecular noise sources into intrinsic and extrinsic components^24^, leading to numerous analytical^25–28^ and experimental^29–31^ extensions that leverage the lower moments of poorly-characterized biological drivers to describe or delimit the system variability. These approaches, which have found application to new experimental techniques, have origins in the Onsager and Langevin theories of the early twentieth century^32^, which specify the moment behaviors of near-equilibrium statistical thermodynamic systems using Gaussian terms.

Alongside studying biology on a gene-by-gene basis, considerable effort has been dedicated to the discovery of regulatory networks. This problem is considerably more challenging: the number of candidate network modules rapidly grows with the number and size of motifs of interest, and simple moment-based models risk distorting key qualitative features, such as multistability. From the perspective of statistics, network inference requires specifying or bypassing likelihood functions for joint gene expression, which may combine various noise sources in addition to the “signal” of regulation. Typical ways of addressing this challenge include^33,34^:

1. **The purely descriptive approach**, which interprets an expression correlation matrix as a graph, but does not provide an easily interpretable way to extract its “signal.”
2. **Thresholding**, which bins the unknown observed distribution to obtain a known, but lower-information distribution, as with binarization used to construct Boolean networks^35^ or implement the phixer algorithm^36^.
3. **Distributional assertion**, which fits static observations by assuming statistics or observations are Gaussian, as in a variety of popular Bayesian^34^, information-theoretic^37^, and regression-based^38^ methods; this assumption may^39^ or may not^40^ provide accurate results.
4. **The dynamic approach**, which fits a time-dependent trajectory to data using assuming Gaussian residuals; this assumption may reflect stochastic differential equation dynamics^41^ or isotropic observation noise added to a latent process^42–44^.

This overview is far from exhaustive, but it demonstrates a key theme: relatively robust signal, such as the lower moments or the absence/presence of gene expression, can be treated using fairly simple models that rely on highly optimized, well-understood methods and algorithms developed in the context of signal processing and dynamical systems analysis. Which simple model may perform best is not known *a priori*, and heavily depends on the task^33^. Ideally, methods are benchmarked on simulated^39,45^ or well-characterized “gold standard”^33,46^ datasets to glean partial insight about their performance and limitations. In this framework, improving the signal-to-noise ratio requires either designing more precise readouts or sacrificing a portion of the obtained data.

### 2.2 The challenge of single-cell data

Advances in sequencing technologies, most dramatically the rapid commercialization and adoption of single-cell RNA sequencing (scRNA-seq), which can profile millions of cells on a genome-wide scale^47,48^, have been heralded as a promising frontier for systems biology^49–51^. This potential is more striking yet due to simultaneous advances in multiomics, or the measurement of multiple modalities (transient and non-coding RNA species, DNA methylation, chromatin accessibility, surface protein abundance) in individual cells^52,53^, facilitating “integrated” analysis^54–56^. The “big data” from single-cell sequencing have thus served as substrate for a plethora of investigations which are, at first glance, analogous to the research program of systems biology at large: the identification of cell types; their aggregation into trajectories; the discovery of gene modules that consistently differ between cell types or throughout a differentiation trajectory; and the visualization of low-dimensional summaries reflecting some component of the data structure.

To identify these coarse-grained motifs in the structure of single-cell datasets, it is common practice to analyze cell–cell graphs, constructed from measures of expression similarity, to attempt to construct cliques (cell types), shortest paths (trajectories), and neighborhood-preserving low-dimensional embeddings (visualizations). In addition, relatively simple parametric distributions are widely used, with the Gaussian assumption popular for the lower moments (e.g., to compute measures of differential expression), and the lognormal or negative binomial used to describe count distributions^57,58^. Standard single-cell RNA sequencing data provide snapshots of processes, rendering dynamical analysis fairly complex, but it is common to fit a “pseudotemporal” curve through the dataset by minimizing a Gaussian error term between this curve and some transformation of the cells’ expression levels^59,60^.

Here, however, the underlying assumptions break down. Single-cell data are intrinsically and qualitatively different from readouts of typical systems biology experiments, with drastic implications for analysis. Single-cell data are large and sparse, with a preponderance of technical noise effects, poorly characterized batch- and gene-level biases, and low per-cell copy numbers^13–15^. Improving the signal-to-noise ratio by designing more targeted experiments is challenging, as commercial technology is designed to quantify molecules on a genome-wide scale. More problematically, typical distributional assumptions and data transformations risk losing a considerable amount of signal in the low-copy number regime. This challenge informs part of the broader discussion of the relative roles of data analysis and mechanistic hypotheses in genomics^19,20,61^, as analyses are not constrained by mechanism or theory and may contradict existing knowledge.

More specific critiques have considered whether various analyses are appropriate or excessively heavy-handed. For example, sparsity has led to *ad hoc* procedures to “correct” the data, which may in turn lead to incorrect conclusions^62–64^. Normalization and log-transformation, which attempt to remove technical biases and prepare the data for dimensionality reduction, rely on assumptions, such as high copy numbers and homogeneity, that are routinely violated in single-cell datasets^65,66^. Dimensionality reduction risks distorting both local and global relationships between data points^19,67,68^. Finally, the use of cell–cell graphs constructed from noisy data reifies relationships which may not reflect those in the original tissue, and risks introducing hard-to-diagnose errors into downstream analysis^19,69^. Although these issues span the entire process of analysis, all, at least partially, trace back to uncomfortable compromises in the treatment of uncertainty and variation in a regime unforgiving of approximations.

### 2.3 Stochastic modeling of intracellular network dynamics

Stochasticity is, then, mandatory, and we ignore it at our own risk. Therefore, we advocate for probabilistic alternatives to the “extraction” of signal from scRNA-seq datasets. Since biology is stochastic, the noise *is* the signal. To quantify and characterize the components of deterministic mental models—differentiation landscapes, kinetic parameters, and similar low-dimensional abstractions^70^—in a principled way, we need to combine them with stochastic terms which result from specific hypotheses about the underlying biophysics and chemistry^20^, or risk confirmation bias^19^.

The development of stochastic models offers advantages beyond loss function book-keeping. If multiomic data are available, there is typically a self-consistent way to extend the models accordingly^71^. Although likelihoods induced by stochastic processes are challenging to analyze and implement, they provide appealing statistical properties. When the data are sufficiently informative, full distributions provide better estimates than moments^40^. When they are not, probabilistic approaches are appropriately conservative, as they report, rather than elide, the parameter degeneracies. A thorough mathematical understanding of model behaviors—i.e., precisely which parameters are identifiable and which are degenerate, as well as how much data must be collected—enables the design of informative experiments^20,72^. Finally, the use of mechanistic models, parametrized by rate constants, allows us to draw conclusions about the mechanistic bases and effects of perturbations^73^.

These principles have guided fluorescence-based singlecell transcriptomics for nearly twenty years. To obtain as much information as possible from entire copy-number distributions^40,74^, the field has developed a considerable arsenal of theoretical tools^75,76^ and solution strategies^77–79^. It is, then, particularly natural to build scRNA-seq models that *extend* processes consistent with fluorescence imaging: this approach allows us to leverage existing theory, as well as encode the intuition that technology-dependent effects should be independent from biological ones. A particularly popular class of models involves the bursty production of RNA and its Markovian degradation^73,80^, which can be analyzed in the chemical master equation (CME) framework^81,82^. The key theoretical points have already been applied in the context of single-cell sequencing; for example, the Poisson, Poisson-gamma, and Poisson-beta distributions, which are common in sequencing analyses^58,63,83,84^, are three of the limiting distributions induced by this class of models^20,80,85^. However, this possible mechanistic basis is only rarely^84,86–88^ invoked in the development of analysis methods.

### 2.4 Outlook

Unfortunately, we cannot simply apply existing methods from fluorescence transcriptomics; the scale and chemistry of singlecell technologies create additional desiderata. General CME solutions are computationally prohibitive and challenging to scale to thousands of genes^89^, requiring careful study of narrow model classes with tractable solutions^17,20^. In addition, connecting biological models to observations requires explicitly representing the experimental process. The existing models for fluorescence data are sophisticated^79^, but cannot be directly applied to sequencing data. Although a variety of models have been proposed for technical noise in single-cell technologies^13,14,90,91^, their chemical foundations, as well as implications for biological parameter identifiability, have been understudied^21^.

In light of this lacuna, we seek to produce a mathematical framework that (1) integrates biological and technical variability in a coherent, modular way; (2) scales to large, multimodal data; (3) can be used to simulate datasets and make testable, quantitative predictions; and (4) affords a thorough mathematical analysis of its components, if not the entire model.

## 3 STOCHASTIC MODELING OF SINGLE-CELL BIOLOGY

Constructing a general-purpose framework for the stochastic modeling of single-cell biology necessitates working at a relatively high level of abstraction, since we would in principle like to account for a range of processes with one formalism. In this section, we motivate our abstract formalism using a collection of concrete, biologically relevant examples.

One of the simplest models of transcription is the *constitutive model*, which assumes RNA is produced at a constant rate^20,92^. It is defined by the chemical reactions

### Box 1

**Generating function methods for studying stochastic biological systems**

Generating functions are ubiquitous tools in stochastic modeling. They are central to the analysis of discrete master equations, as they cast difficult-to-solve infinite-dimensional systems to partial differential equations, which can be treated using standard analytical or numerical methods. A (one-variable) probability distribution *P*(*x*) and its generating function *G*(*g*) are related according to the formulas:

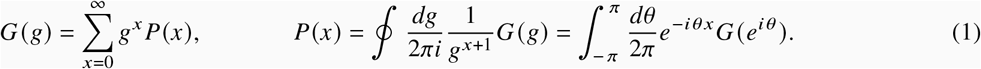

In the stochastic modeling of transcription, certain distributions, such as the Poisson and negative binomial, frequently appear. Because *G* uniquely specifies *P*, we can often invert *G* simply by recognizing its form and matching terms. Below are some generating functions of common distributions (Bernoulli, Poisson, geometric, and negative binomial):

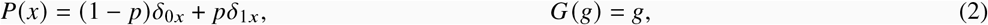

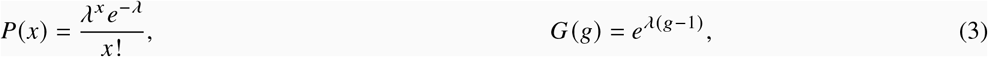

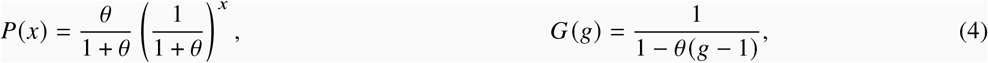

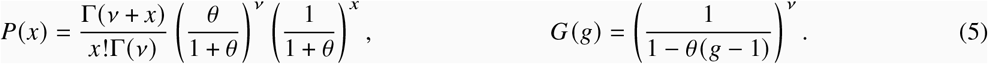

The generating function expressions can often be made more compact by applying the substitution *u* := *g* − 1.

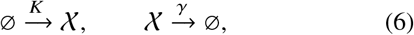

where 𝒳 is a single species of RNA, *K* is the (constant) transcription rate, and *γ* is the degradation rate. The CME that corresponds to this system is

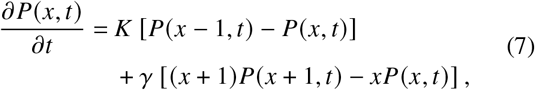

where *P*(*x, t*) is the probability that the system has *x* ∈ ℕ _0_ RNA at time *t*. Solving the above master equation allows us to compare its predictions with experimental scRNAseq data. There are several theoretical approaches for doing this—including using a special ansatz^85^, the Poisson representation^93^, the Doi-Peliti path integral^17,94–96^, and operator techniques^97^—but we would like to highlight a straightforward method that we know works for far more general problems. The idea is to consider a certain transformed version of the probability distribution, which satisfies a partial differential equation (PDE) instead of a differential-difference equation. This PDE, for a large class of biologically relevant systems, can then be solved using the method of characteristics^98^, which converts the problem of solving a PDE into integrating a system of ordinary differential equations (ODEs). This is mathematically equivalent to using certain path integral methods^17,20,99^.

Define the generating functions (GFs)

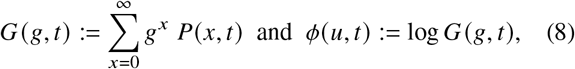

where *g* is on the complex unit circle and *u* := *g* − 1. It is easy to show that *G* and *ϕ* satisfy the PDEs

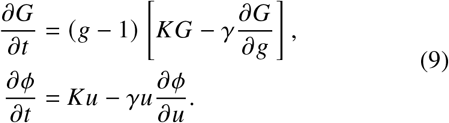

We can use the method of characteristics to find that

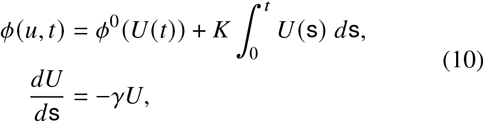

where the *U*(s) ODE has initial condition *U*(s = 0) = *u*, and where *ϕ*^0^ is the initial (log-) generating function of the system. In order to determine *P*(*x, t*) from *ϕ*(*u, t*) = log *G*(*g, t*), we can use an inverse Fourier transform:

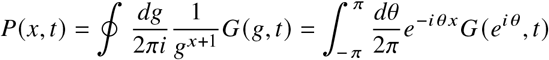

where we integrate over all *g* on the complex unit circle. In practice, this step is done numerically using an inverse fast Fourier transform.

The constitutive model, which produces Poisson distributions at steady state, is too simple for single-cell biology^20^. But fortunately, the technique we have just described can be adapted to predict the behavior of substantially more complex models.

### Multiple types of RNA

One possible generalization of the constitutive model is to so-called *monomolecular systems*^17,85^, which allow phenomena like RNA splicing to be accommodated. An example is the addition of splicing to the constitutive model:

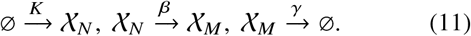

In general, any number of production, conversion, and degradation reactions can be modeled:

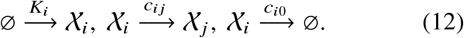

Using the same technique we described earlier, the probability *P*(**x**, *t*) that the system is in state 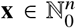 at time *t*, can be shown to be equivalent to the generating function

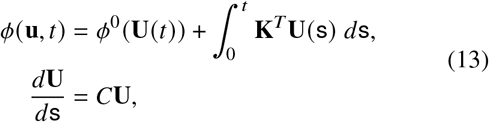

where **U**(s = 0) = **u**, and the *C* matrix is defined via

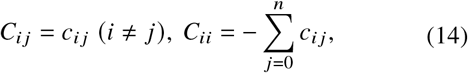

and where *c*_*ii*_ := 0 by convention.

### Multiple gene states

Although the monomolecular model is a step forward, it still does not account for nontrivial transcription rate dynamics. One possibility is that there are multiple gene states, as in the telegraph model^76,97,100^:

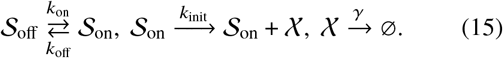

The corresponding three-variable generating function is

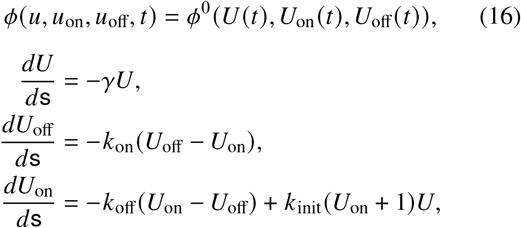

where *U*(0) = *u, U*_off_ (0) = *u*_off_, and *U*_on_ (0) = *u*_on_. If we want to marginalize over gene state, which we usually do since it is not observable, we can set *u*_off_ = *u*_on_ = 0. Notice that the relevant ODEs are now nonlinear (Riccati-type) equations, which make them difficult to solve by hand. In general, considering multiple gene states, or other kinds of added complexity like autocatalytic reactions, yields nonlinear characteristic ODEs. This is no obstacle for numerical integration, however.

### Gene regulation

Another possibility we would like to account for is nontrivial gene regulation. In previous work^20^, we considered two models of transcription rate variation: the gamma Ornstein–Uhlenbeck (Γ-OU) model, which assumes variation is due to changes in the mechanical state of DNA; and the Cox–Ingersoll–Ross (CIR) model, which assumes it is due to fluctuations in the concentration of an abundant regulator molecule. Analyzing them can be mathematically challenging, since the discrete stochastic dynamics of RNA production and degradation are coupled to the continuous stochastic process that controls the transcription rate. Fortunately, both models and many generalizations of them can be solved using the method of characteristics. For example, the CIR model (assuming two RNA species) is defined by a stochastic differential equation (SDE) and three reactions:

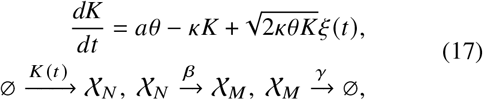

and its solution is^20^

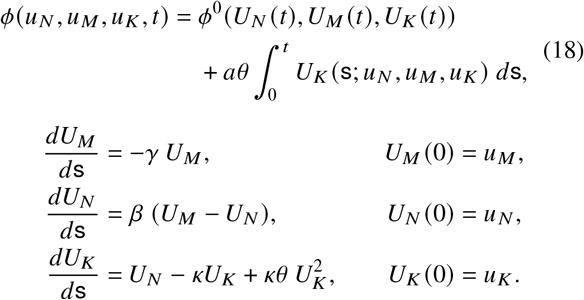

Thus, it is straightforward to couple dynamics defined on different types of state spaces: categorical (e.g., gene states), continuous (e.g., transcription rates), and discrete (e.g., RNA counts), using the generating function approach. In all cases, one obtains a generating function solution in terms of a finite set of (possibly nonlinear) ODEs. The total number of ODEs is equal to the total number of degrees of freedom.

One feature of single-cell biology that is challenging to capture using this approach is feedback. For example, proteins expressed by a gene may affect the transcription rate of that gene. Although exact solutions for systems involving feedback are available in certain simple cases^101–104^, particularly when there is only one chemical species, more general results have proven elusive. From the point of view of our approach, including chemical reactions that involve feedback yields generating function PDEs which are not first order, and cannot be solved in terms of ODEs via the method of characteristics (as explored in more detail in the supplemental information).

### Transient effects

In the context of development or reprogramming, we are especially interested in using single-cell genomics data to study transient processes. In particular, certain cell types or subtypes (like neural progenitor cells) only exist for a certain window of time, and by collecting single-cell data we are taking a snapshot of many cells, each of which may be in a different part of the process. How does this affect observed RNA counts?

Different cells being observed at different times means we are not interested in *P*(**x**, *t*), but *P*(**x**, *t*) averaged over some distribution that indicates how likely we are to sample different times. The shape of the sampling distribution *f*(*t*) depends on when cells tend to exit a given state (e.g., by differentiating into a different cell type). Nontrivial sampling distributions are compatible with our generating function approach, since we can simply modify the distribution that appears. For a model with one discrete species, we can write the full generating function *G*_tot_ as

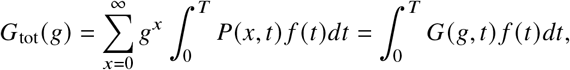

i.e., we can obtain it by integrating the generating function that captures intrinsic noise.

### Technical noise

In single-cell genomics experiments, we do not directly observe a given cell’s RNA counts, but those numbers filtered through a noisy sequencing process^21^. In microfluidics-based sequencing, noise can come from some combination of droplets not capturing all molecules (especially types of RNA with low copy numbers), errors in amplification, and reads not being uniquely identifiable. We would like to account for these kinds of technical noise in a way that is both principled, and compatible with our generating function approach to modeling intrinsic noise.

Consider a simple example, in which the relevant biology is described by the one-species constitutive model (Equation 7), and each RNA molecule is observed independently with probability *p*. The probability of observing *x*_obs_ molecules of RNA, given a biological distribution *P*(*x, t*), is

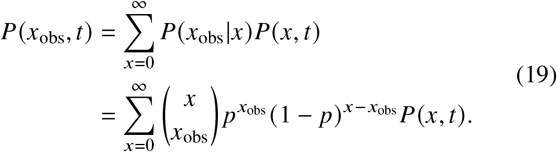

The corresponding generating function *G*_tot_ is

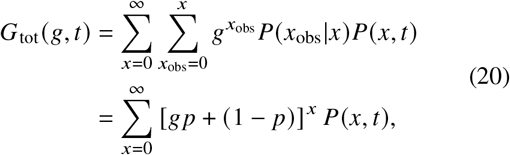

i.e., the result is the same as without technical noise, except that we have *g* → *gp*+(1 − *p*). In general, including technical noise requires us to replace the usual *g*^*x*^ factor with *G*_noise_ (*g, x*), the generating function associated with the observation model:

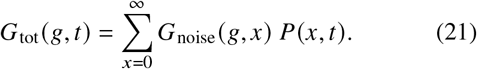

For certain common observation models, like the Bernoulli model just described, or a Poisson noise model, we can say more: since

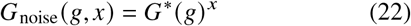

for some *G*\*, including technical noise amounts to replacing *g* with *G*\*, so that *G*_tot_ = *G*(*G*\*) is a composition of generating functions. We typically assume that all technical noise models satisfy Equation 22 for some *G*\*.

## 4 RESULTS

### 4.1 Theoretical framework for stochastic systems biology

We are ready to present our general framework for stochastic systems biology, which accommodates all of the sources of stochasticity described in the preceding section: intrinsic noise, transient effects, and technical noise. In order to balance the amount of biology our models can capture with the mathematical tractability of those models, we restrict our analysis to a fairly general class of systems that can be solved using the method of characteristics. For such systems, we can obtain likelihoods by integrating characteristic ODEs, using the obtained characteristics to construct the generating function, and then doing an inverse (fast) Fourier transform.

This class of systems permits gene state interconversion, as well as the production and processing of RNA and proteins, which could treated as discrete or continuous variables depending on their concentration. We allow zero- and first-order reactions, including state-dependent bursting, interconversion, degradation, and catalysis. However, we disallow higherorder reactions (e.g., binding reactions *A* + *B* → *C*), including feedback-based regulation like protein-promoter binding. Therefore, our analysis focuses on Markovian systems that possess *N* categorical degrees of freedom, corresponding to gene states; *n* discrete ones, corresponding to low-copy number molecular species; and *m* continuous ones, corresponding to transcription rates or high-concentration species. This class of reactions is schematically represented in Figure 1a; crucially, it consists of distinct “upstream” and “downstream” components.

**Figure 1.**
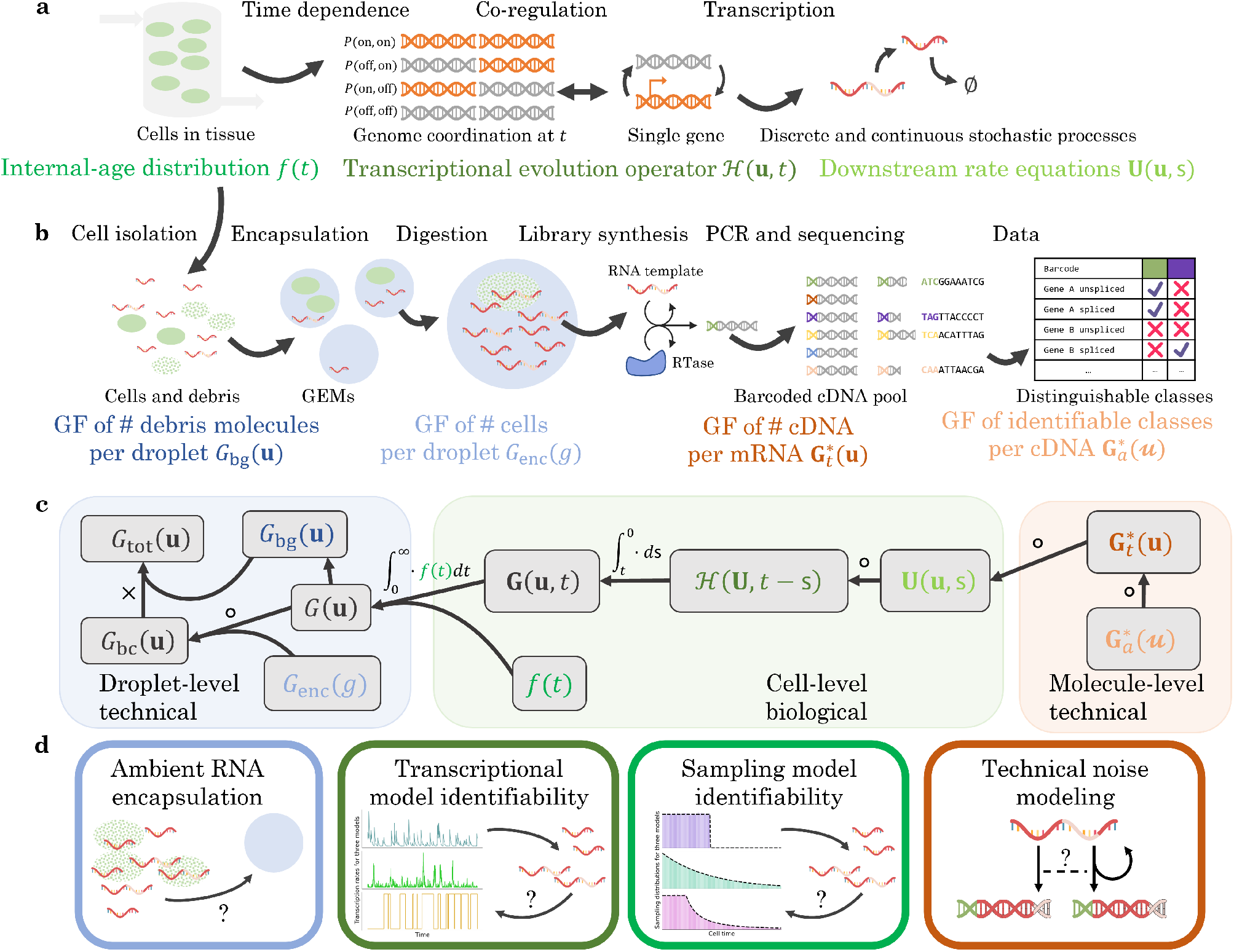
The biophysical and chemical phenomena of interest, as well as the relationships between their generating functions. **a**. The biological phenomena of interest: cell influx and efflux into a tissue observed by sequencing; the time-dependent transcriptional regulation of one or more genes; downstream continuous and discrete processes. **b**. The technical phenomena of interest: the encapsulation of cells and cell debris; cDNA library construction; the loss of information in transcript identification (GF: generating function; RTase: reverse transcriptase). **c**. The structure of the full generating function of the system in **a** and **b**: to obtain the solution, we variously compose, integrate, and multiply the generating functions of the constituent processes. **d**. The stochastic and statistical properties of four components of the full system: the background debris, the transcriptional regulation, the cell/tissue relationship, and the technical noise mechanism.

Given all of a model’s possible reactions, one can write down a corresponding master equation that keeps track of how microstate probabilities change with time:

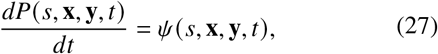

where each microstate consists of *s*, the categorical dimension; **x** ∈ ℕ^*n*^, the *n* discrete dimensions; and **y** ∈ ℝ^m^, the *m* continuous dimensions. The generally complicated function *ψ* aggregates all reaction rates. Master equations like Equation 27 typically consist of an infinite system of coupled ODEs, and hence are difficult to solve in general. This is one reason we chose a particular class of systems: to solve Equation 27 using the method of characteristics, and hence determine a given model’s predictions, all we need to do is solve (a finite number of) ODEs satisfied by the characteristics and GF.

#### Box 2

**An illustration of the solution procedure**

Here, we will illustrate how to solve two simple transcription models using our framework. We assume that RNA is produced with burst event frequency *α* and degrades at a rate *γ*. In the *constitutive* model, each transcription event creates one RNA. In the *bursty* model, each transcription event creates a random number of RNA, distributed according to a geometric random variable with mean *b*. Both models have *N* = 1, *n* = 1, and *m* = 0. Since these models are one-dimensional, the *C* and *D* matrices are 1 × 1. For both of them, *C* = [−*γ*] and *D* = [0]. The ODE for the single characteristic *U* (with initial condition *U*(s = 0) = *u*) is

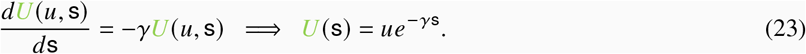

For a general burst distribution *p*(*z*), the transcriptional evolution operator is ℋ(*u*) = −*α*(*F*(1 + *u*) − 1), where *F* is the GF of the number of molecules produced per transcription event. For our two models, we have

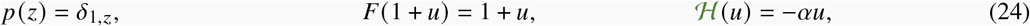

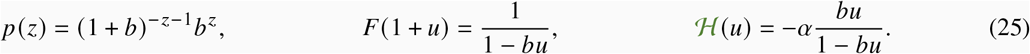

To compute the stationary log-generating functions log *G*, we evaluate the integrals:

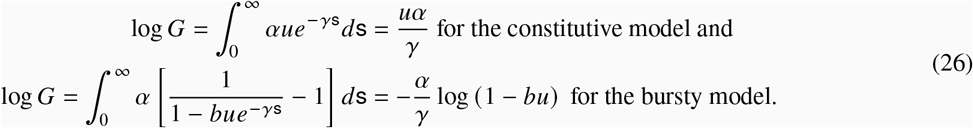

The constitutive model yields a Poisson distribution with mean *α*/*γ* (c.f. Equation 3), whereas the bursty model yields a negative binomial distribution with shape *α*/*γ* and scale *b* (c.f. Equation 5).

The *N*-dimensional GF **G** = (*G*_1_, ⋯, *G*_*N*_)^*T*^ of the system, which is a function of spectral variables **g** and **h**, is defined by

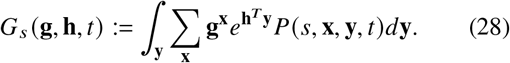

Equation 27 can be converted into a PDE satisfied by **G**:

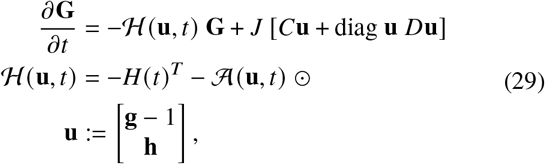

where ⊙ is the Hadamard/elementwise matrix product, *J* is the Jacobian matrix of the generating function, and **u** combines the discrete and continuous degrees of freedom. The time-dependent matrix *H* contains the kinetics of state transitions, whereas the operator 𝒜 describes the drift and bursty production processes, which may depend on state. Therefore, the operator ℋ aggregates the upstream components of the system. The matrix *C* contains interconversion, degradation, and mean reversion-like terms, whereas *D* contains the catalysis and square-root noise terms. ℋ, *C*, and *D* encode a quasi-linear, deterministic, and first-order *N*-component system of partial differential equations in *n* + *m* spectral variables.

Applying the method of characteristics to solve Equation 29 tells us that the downstream part of the system is fully determined by a set of characteristics **U**, which are defined by the ODEs

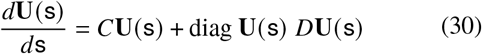

where s is an integration variable, and **U**(s = 0) = **u**. Meanwhile, the generating function **G** can be determined from

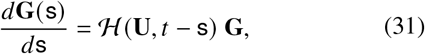

which has initial condition **G**^0^(**U**(*t*)), where **G**^0^ is the generating function of the initial distribution. The upstream components describe how molecule production occurs, and hence depend on ℋ; their influence on the final answer is through the above integral.

The detailed form of Equation 27 is complicated, and the arithmetic exercise of converting it into Equation 29 is tedious. We show how to construct the biological master equation in Section 6.1, write it out in full in Section 6.2, and discuss at a high level how to solve it using our generating function approach in Section 6.3. The terms of the full master equation are annotated in Table S1, and the solution process is described in more detail in supplemental information.

In special cases, the ODEs we obtain can be solved exactly. For example, whenever *D* = 0, the downstream ODE system can be solved analytically by eigendecomposition. If, in addition, only a single gene state is present, *H* vanishes and the upstream component can be evaluated by numerical integration^16^. Finally, in the simplest case of a linear operator 𝒜, we obtain an analytically tractable system equivalent to a deterministic system of reaction rate equations^17,85^.

Although this formulation nominally describes a single gene, it may be exploited to represent multi-gene systems. Conceptually, this strategy entails constructing a model where the transcription of multiple species is controlled by a common regulator. We discuss potential candidate models in Section 6.4; these models instantiate hypotheses to produce ℋ and **U** that represent co-regulation.

To explain the observation of transient processes, such as the simultaneous capture of progenitor and descendant cells from a differentiation process, we take inspiration from previous work in sequencing^86^ as well as chemical reactor modeling^105,106^, and extend the theoretical framework originally proposed in our recent RNA velocity analysis^19^. In brief, the simplest model that accounts for such desynchronization proposes that cells enter a tissue, receive a signal that triggers changes in transcriptional rates ℋ (*t*), and leave at some later point. Sequencing is the observation of cells within the tissue; to find the distribution of RNA counts, we need to condition on the distribution of times since receiving the signal.

As we discuss in Section 6.5, this latter distribution is not arbitrary, and reflects the kinetics of cell entry and exit. In the parlance of chemical reaction engineering, the times are drawn from *f*(*t*), the *internal-age distribution* induced by those kinetics^105,106^. This model affords a particularly simple representation of the generating function:

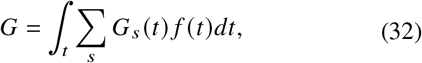

where we marginalize over the gene state, which is typically not observable. Conveniently, this model possesses time symmetry: even though the cells within the tissue are all out of equilibrium, the tissue as a whole is at steady state.

We consider the technical noise phenomena shown in Figure 1b, i.e., the encapsulation of cells and background debris into droplets, as well as the stochasticity in cDNA library construction and sequencing. Under the assumption of independent encapsulation, the generating function of molecule count distributions on a per-droplet level takes the following form:

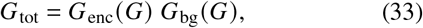

where the *G*_enc_ is the generating function of the cells per droplet, whereas *G*_bg_ is the generating function of background molecules per droplet, which depends on the entire cell population (Section 6.6). Finally, to represent sequencing variability and uncertainty, we evaluate the generating function at a set of transformed coordinates:

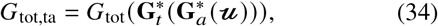

where 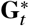 reflects the distribution of cDNA produced per molecule of RNA (e.g., Bernoulli, as in Tang et al.^107,108^), whereas 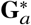 reflects the distribution of ambiguous sequenced fragments, which depends on transformed variables *ᓚ*(Sections 6.7 and supplemental information).

The full generating function of the molecule distribution is given by the composition and integration of the model components, as shown in Figure 1c. To evaluate this generating function, it is necessary to specify all components that make up the model. In the analysis below, we take advantage of the modularity of the system definition to investigate four kinds of modeling choices, their statistical implications, and their compatibility with sequencing data. Specifically, we treat the subsystems illustrated in Figure 1d: background noise in single droplets, stochastic transcription rate models, sampling from a transient process, and variation in technical noise.

### 4.2 Empty droplets

One of the first steps in scRNA-seq data analysis is cell quality control, which excludes cell barcodes that appear to originate from empty droplets from further analysis^57^. For computational tractability, this procedure typically relies on “hard” assignment, such that barcodes associated with a total molecule count above some threshold are treated as cells, whereas barcodes below the threshold are treated as empty droplets. Threshold selection is necessary because even “empty” droplets contain ambient RNA. This ambient RNA, which appears to originate from cells lysed in the preparation process, contaminates empty and cell-containing droplets alike^57^.

The observation of ambient RNA resulting in unwanted molecule counts has led to the development of statistical methods for removing this source of noise, either by estimating and subtracting it^109^ or incorporating it into a stochastic model^110–112^. Conceptually, Equation 33 reflects the latter approach: each droplet contains one or more cells, each with biological generating function *G*, and background, with a generating function *G*_bg_ that depends on *G*. To accurately model the background counts, we need to propose and justify a specific functional form for *G*_bg_. Thus, under the assumption that empty and cell-containing droplets are similarly susceptible to contamination, the former provide a reasonable estimate of ambient distributions in the latter^109^.

The simplest model holds *G*_bg_ to be equivalent to a “pseudobulk” experiment, with molecules randomly sampled from the lysed cell population. If each cell is equally likely to contribute to the pool of free RNA, and diffusion occurs by a simple independent arrival process, we find that the distribution of background should be Poisson, with the mean for each species proportional to its mean in the original cell population, as in, e.g., Fleming et al.^110^ This functional form immediately induces a set of testable predictions: not only are the distributions Poisson, but they are *independent* Poisson, with no meaningful statistical structure remaining between transcripts of a single gene, as well as between different genes, as illustrated in Fig. 2a.

**Figure 2.**
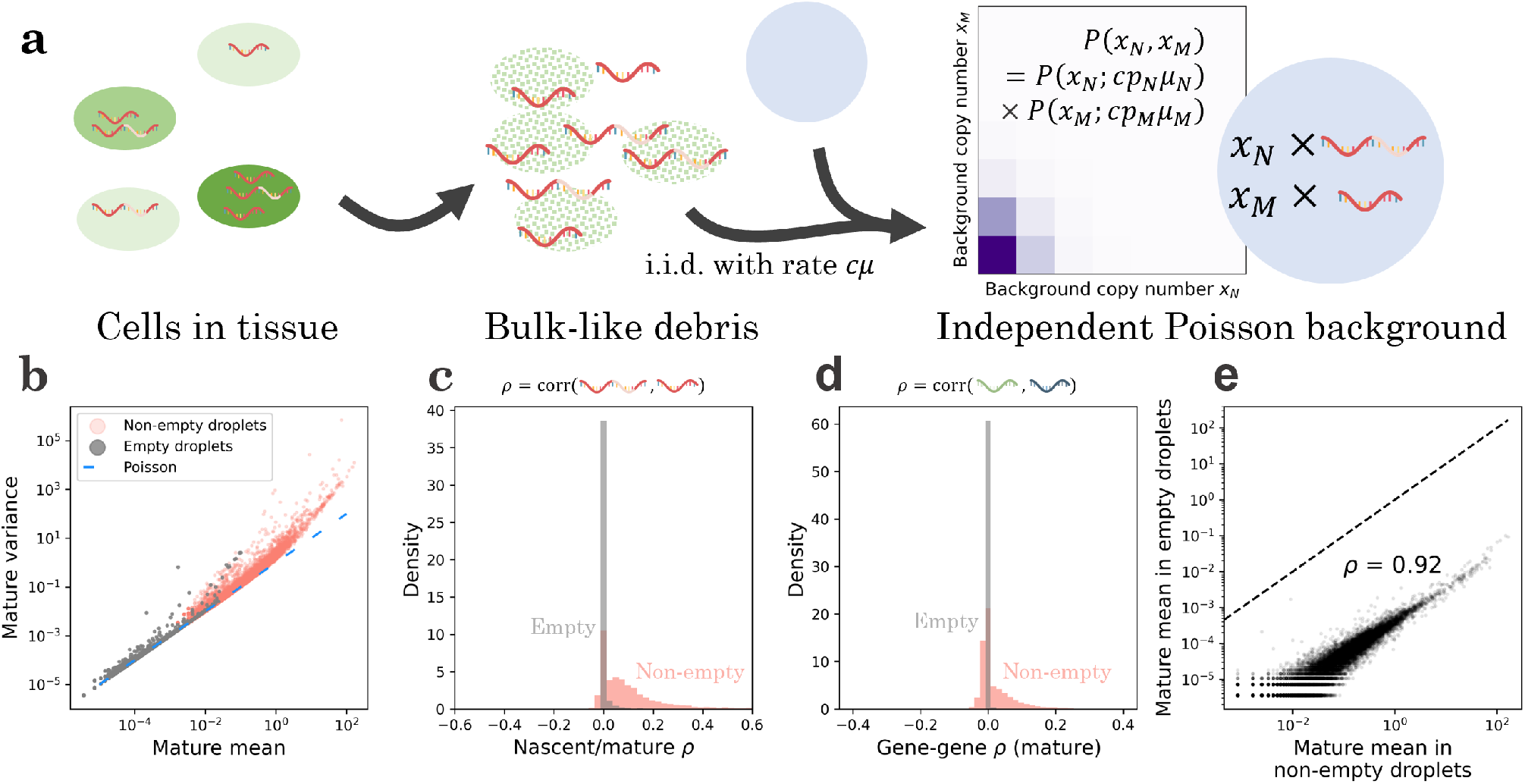
The pseudo-bulk model of background noise is quantitatively consistent with counts from the `pbmc_1k_v3` dataset. **a**. The simplest explanatory model for background noise invokes the lysis of cells (green), which creates a pool of RNA that reflects the overall transcriptome composition but retains none of the cell-level information. If the loose RNA molecules diffuse into droplets (blue) according to a memoryless and independent arrival process, the resulting background distribution (purple: higher probability mass; white: lower probability mass) observed in empty droplets should be a series of mutually independent Poisson distributions, with the mean controlled by the composition in non-empty droplets. **b**.The mature transcriptome in empty droplets has a mean-variance relationship near identity (gray points, *n* = 12, 298), consistent with Poisson statistics (blue line); the non-empty droplets demonstrate considerable overdispersion (red points, *n* = 17, 393). **c**.The mature and nascent transcripts in empty droplets have sample correlation coefficients *ρ* near zero, consistent with distributional independence (gray histogram, *n* = 9, 362); the non-empty droplets demonstrate nontrivial statistical relationships (red histogram, *n* = 14, 365). **d**.The mature transcripts of different genes in empty droplets have sample correlation coefficients *ρ* near zero, consistent with distributional independence (gray histogram, *n* = 75, 614, 253); the non-empty droplets demonstrate nontrivial statistical relationships (red histogram, *n* = 151, 249, 528). **e**.When both are nonzero, the mature count mean in empty droplets is highly correlated with the mean in the non-empty droplets, consistent with the pseudo-bulk interpretation (black points, *n* = 12, 107; dashed line: identity).

To characterize the accuracy of these predictions, we inspected six datasets (Table S2) pseudoaligned with *kallisto* | *bustools*^113^, and compared the data for barcodes passing *bustools* quality control to data for barcodes which were filtered out. As a shorthand, we call the former “non-empty” and the latter “empty” droplets, keeping in mind that this identification is approximate. We fully describe the analysis procedure in Section 6.8.2, illustrate the results for the human blood dataset `pbmc_1k_v3`, and display the results for all datasets in supplemental information.

As shown in Figure 2b, data from non-empty droplets are substantially overdispersed relative to Poisson, whereas data from empty droplets are largely consistent with the Poisson identity mean–variance relationship. However, a small number of relatively high-expression genes are overdispersed. In addition, intra-gene (Figure 2c) and inter-gene (Figure 2d) correlations are typically nontrivial in non-empty droplets, but consistently near zero for empty droplets, supporting distributional independence of the background counts. Finally, the mean expression in empty droplets is highly correlated with mean expression in non-empty droplets, albeit lowered by approximately four orders of magnitude (Figure 2e), supporting the assumption that the original cells are lysed in a uniform fashion.

To characterize the deviations from the pseudo-bulk model, we identified the genes that demonstrated overdispersion in empty droplets (Table S3). A considerable fraction of these genes were associated with mitochondria or blood cells. For example, of the 21 annotated genes overdispersed in the empty droplets of the mouse neuron dataset `neuron_1k_v3`, nine were mitochondrial (*mt-Nd1, mt-Nd2, mt-Co1, mt-Co2, mtAtp6, mt-Co3, mt-Nd3, mt-Nd4*, and *mt-Cytb*), three coded for hemoglobin subunits (*Hba-a1, Hba-a2*, and *Hbb-bs*), and two coded for blood cell-specific proteins (*Bsg, Vwf*)^114,115^. On the other hand, of the 10 annotated genes overdispersed in the empty droplets of the desai_dmso dataset, generated from cultured mouse embryonic stem cells^116^, six (*mt-Nd1, mt-Co2, mt-Atp6, mt-Co3, mt-Nd4, mt-Cytb*) were mitochondrial and none were blood cell-specific^114^ (Table S4).

Since overdispersion implies that contamination involves non-independent encapsulation of these molecules, the results suggest that the cell-free debris contain, among other structures, entire mitochondria or erythrocytes, when they are present in the source tissue. These membrane-bound structures may diffuse into droplets, then lyse and release all of their contents at once. In other words, empty droplets do not merely have disproportionally high mitochondrial content, as has been noted previously^110,117,118^; they have *nontrivially distributed* mitochondrial content, which can hint at the mechanism of its incorporation, and improve interpretation where simple thresholds may be misleading^118^. We hypothesize that cases where the model fails can be leveraged to discover more complicated forms of contamination, such as molecular aggregates^112^.

In addition, we examined the total UMI counts in empty droplets, which should be Poisson (Fano = 1) if each individual gene’s distribution is Poisson. For the human blood dataset demonstrated in Figure 2, the empty droplets had fairly significant overdispersion (Fano ≈43), which decreased, but did not disappear (Fano ≈7.6), once the 53 significantly overdispersed genes were excluded. This result suggests that, although the pseudo-bulk model is approximately valid, some residual variance, possibly due to variability in per-droplet capture rates, is present and needs to be modeled to fully describe the stochasticity in single-cell datasets.

### 4.3 Noise-corrupted candidate models of transcriptional variation

A considerable fraction of the variability in single-cell datasets arises from cell-to-cell and time-dependent variation in the transcription rates. These sources of variation control distribution shapes. By carefully analyzing candidate models, we can characterize the prospects for model selection: for example, if different models produce nearly identical distributions, selection is impossible and the choice of model is somewhat arbitrary. More interestingly, such analysis can guide the design of experiments: models may be indistinguishable based on some kinds of data, but not others^20^. This perspective has guided the interest in characterizing noise behaviors^74,119^: distributions provide strictly more information than averages, and allow us to distinguish between regulatory mechanisms. Similarly, multivariate distributions provide more information than marginal distributions. Obtaining *different* data (multiple molecular modalities) is qualitatively more useful than obtaining more data (a larger number of cells) or better data (observations less corrupted by noise).

We illustrate this key point using the simple model system depicted in Figure 3a, which features intrinsic, extrinsic, and technical noise. The continuous stochastic process denoted by *K* drives the rate of transcription of nascent RNA. We consider three different possibilities for *K*: the gamma Ornstein–Uhlenbeck process, which models DNA winding and relaxation; the Cox–Ingersoll–Ross process, which models the fluctuations in a high-copy number activator^20^; and the telegraph process, which models variation due to random exposure of the locus to transcriptional initiation^76,97,100^. All three transcription rate models are described by three parameters^20,100^. After a Markovian delay, nascent RNA are converted to mature RNA; after another Markovian delay, the mature RNA are degraded. When the system reaches steady state, it is sequenced; each biological molecule has a probability *p* of being observed in the final dataset. We seek to use imperfect count data to fit parameters and distinguish models. We fully describe the procedures in Section 6.8.3.

**Figure 3.**
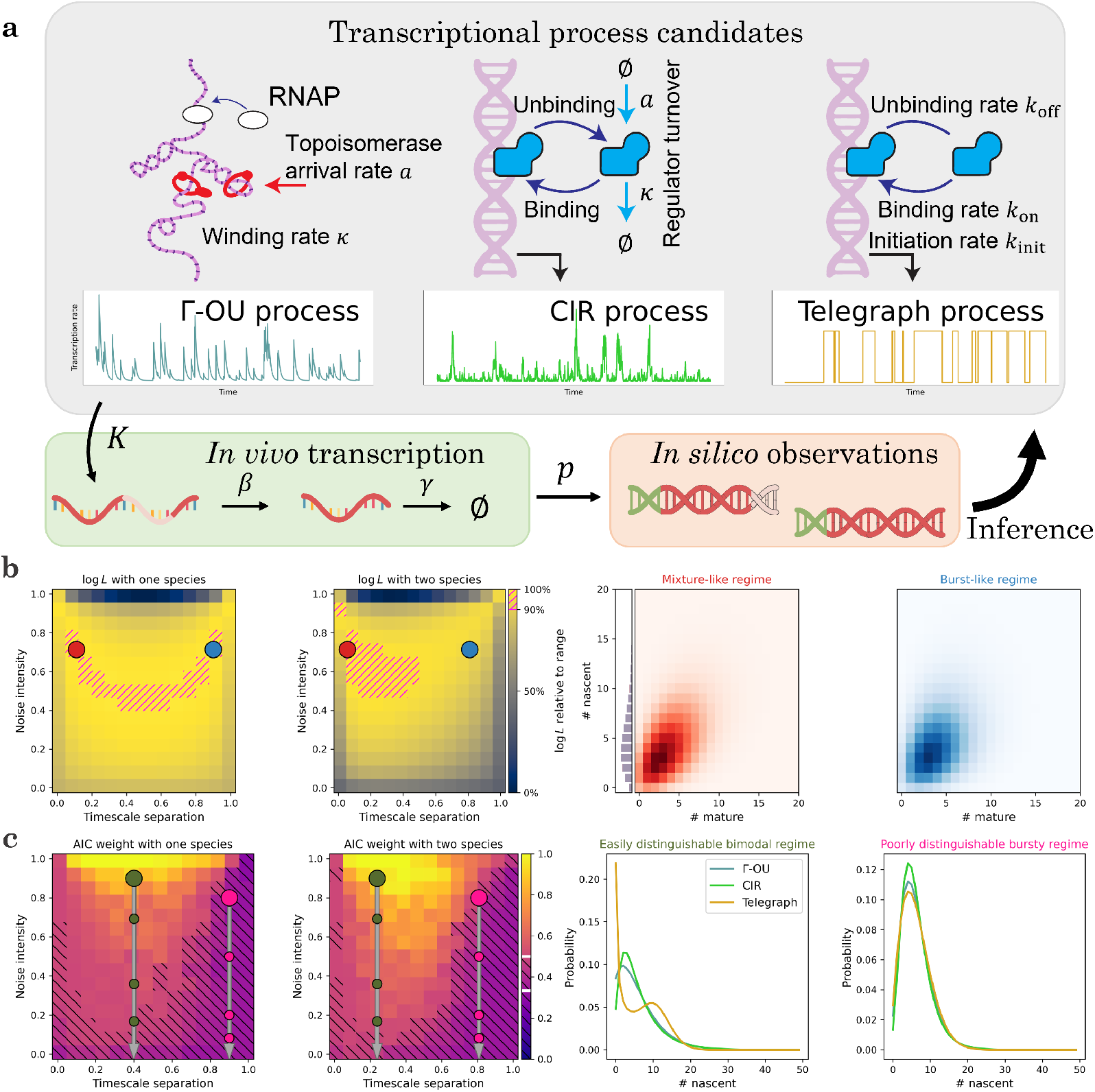
The stochastic analysis of biological and technical phenomena facilitates the identification and inference of transcriptional models. **a**. A minimal model that accounts for intrinsic (single-molecule), extrinsic (cell-to-cell), and technical (experimental) variability: one of three time-varying transcriptional processes *K* generates molecules, which are spliced with rate *β*, degraded with rate *γ*, and observed with probability *p*. Given a set of observations, we can use statistics to narrow down the range of consistent models. **b**. Given a particular model, parameter regimes indistinguishable using a single modality become distinguishable with two. The mixture-like and burst-like regimes both produce negative binomial marginal distributions, but have different correlation structures (Left: data likelihoods over the parameter space, computed from 200 simulated cells; Γ-OU ground truth; red point: true parameter set in the mixture-like regime; color: log-likelihood of data, yellow is higher, 90th percentile marked with magenta hatching; blue: an illustrative parameter set in a burst-like parameter regime with a similar nascent marginal but drastically different joint structure. Right: nascent marginal and joint distributions at the points indicated on the left. Nascent distributions nearly overlap). **c**. Given a location in parameter space, models are easier to distinguish using multiple modalities. However, the performance varies widely based on the location in parameter space and the specific candidate models: for example, the telegraph model has a well-distinguishable bimodal limit when the process autocorrelation is slower than RNA dynamics. In addition, all else held equal, drop-out noise effectively decreases the noise intensity, lowering identifiability (Left: Γ-OU Akaike weights under Γ-OU ground truth, average of *n* = 50 replicates using 200 simulated cells; color: Akaike weight of correct model, yellow is higher, regions with weight < 0.5 marked with black hatching; large circles: illustrative parameter sets; smaller circles: distributions obtained by applying *p* = 50%, 75%, and 85% dropout to illustrative parameter sets while keeping the averages constant. Right: the three candidate models’ nascent marginal distributions at the large points indicated on the left).

Even if we have perfect information about the true averages of the transcriptional strength and the molecular species, the systems can exhibit a wide variety of distribution shapes and statistical behaviors. This variety can be summarized by a two-dimensional parameter space, which was introduced in Fig. 2 of Gorin and Vastola et al.^20^ The “timescale separation” governs the relative timescales of the transcriptional and molecular processes; if it is high, the transcriptional process is faster than RNA turnover. The “noise intensity” governs the variability in the transcriptional process: if it is high, the process exhibits substantial variability that translates to overdispersion in the RNA distributions. The bottom edge of this parameter space produces Poisson distributions of RNA, the top left corner produces Poisson mixtures of the law of *K*, and the top right corner yields bursty dynamics that do not typically have simple analytical solutions^20^.

Although these regimes reflect very different transcriptional kinetics, they can produce indistinguishable distributions. The first panel of Figure 3b demonstrates the likelihood landscape of a dataset generated from the gamma Ornstein– Uhlenbeck (Γ-OU) transcriptional model, evaluated using the nascent marginal and *p* = 1. The mixture-like true parameters are indicated by a red point and the top decile of likelihoods is indicated by hatching. The Γ-OU model’s transcription rate has a gamma stationary distribution, which produces approximately Poisson-gamma, or negative binomial, RNA marginals in this regime. However, the bursty regime, indicated by a blue point, also yields a negative binomial-like marginal^20^, preventing us from identifying the kinetics.

On the other hand, if we evaluate likelihoods using the entire two-species dataset, we obtain the landscape in the second panel of Figure 3b: the symmetry is broken, and the parameters can be localized to the mixture-like regime. The source of this improved performance is evident from examining the distributions, shown in the third and fourth panels of Figure 3b. The nascent marginals are essentially identical; no amount of purely nascent count data can distinguish between them. However, the bivariate distributions show subtle differences, such as higher nascent/mature correlations in the true regime, which can be used for inference. This approach is analogous to Fig. 4b of Gorin et al.^21^, where bivariate data are used to disambiguate differences which would otherwise be indistinguishable due to the degeneracies of steady-state distributions.

**Figure 4.**
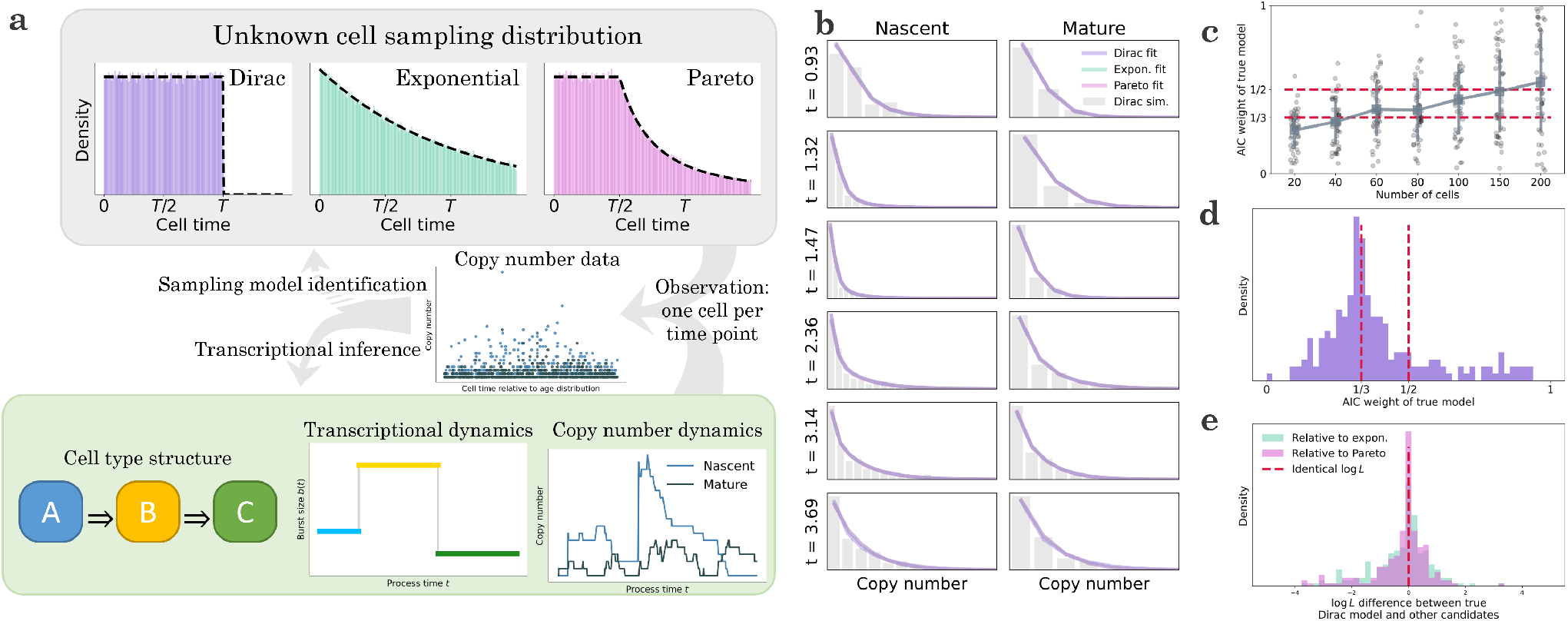
Given ordered and labeled snapshot data obtained from a transient differentiation process, we can typically fit the copy number data, but identifying the mechanism of the snapshot is more challenging. **a**. A minimal model that accounts for the observation of transient differentiation processes in scRNA-seq: cells enter a “reactor” and receive a signal to begin transitioning from cell type A through B and to C. The change in cell type is accompanied by a step change in the burst size, which leads to variation in the nascent and mature RNA copy numbers over time. Given information about the cell type abundances and the cells’ time along the process, we may fit a dynamic process to snapshot data and attempt to identify the underlying reactor type, which determines the probability of observing a cell at a particular time since the beginning of the process. **b**. In spite of the considerable differences between the reactor architectures, they produce nearly identical molecular count marginals (histogram: data simulated from the Dirac model, 200 cells; colored lines: analytical distributions at the maximum likelihood transcriptional parameter fits for each of the three reactor models. Analytical distributions nearly overlap). **c**. The true reactor model may be identified from molecule count data, but statistical performance is typically poor (points: Akaike weight values for *n* = 50 independent rounds of simulation and inference under a single set of parameters; blue markers and vertical lines: mean and standard deviation at each number of cells; blue line connects markers to summarize the trends; red lines: the Akaike weight values 1/3, which contains no information for model selection, and 1/2, which gives even odds for the correct model; two-species data generated from the Dirac model; uniform horizontal jitter added). **d**. The reactor models are poorly identifiable across a range of parameters, and rarely produce Akaike weights above 1/2 (histogram: Akaike weight values for *n* = 200 independent rounds of parameter generation, simulation, and inference under the true Dirac model; red line: the Akaike weight values 1/3 and 1/2; two-species data for 200 cells generated from the Dirac model; parameters were restricted to the low-expression regime *μ* + 4*σ* ≤ 25 for both species). **e**. The challenges in reactor identification arise because all three models produce similar likelihoods (histograms: likelihood differences between candidate models and the true Dirac model for *n* = 200 independent rounds of parameter generation, simulation, and inference; red line: no likelihood difference; two-species data for 200 cells generated from the Dirac model; parameters were restricted to the low-expression regime *μ* + 4*σ* ≤ 25 for both species).

In addition, the timescale separation and noise intensity determine the model distinguishability. To quantify this, we use the Akaike weight *w*_*ϖ*_, which transforms log-likelihood differences into model probabilities^120^. For example, if the Akaike weight is near 1/3, the models are indistinguishable; if the correct model’s weight is near 1, we can confidently identify the model from the data. The first panel of Figure 3c demonstrates the average Akaike weight landscape of datasets generated from the Γ-OU model, computed using the nascent distribution at the same coordinate. We indicate the region *w*_*ϖ*_ < 1/2 by hatching. As the Akaike weight may be interpreted as a posterior model probability^120^, this somewhat arbitrary threshold gives even odds for choosing the correct model, on average.

The intermediate regime, indicated by a large olive green point, tends to yield fairly high Akaike weights, consistent with the two-model case explored in Fig. 3a of Gorin and Vastola et al.^20^ On the other hand, the burst-like regime, indicated by a large pink point, provides considerably less ability to distinguish the models. As expected, the situation improves somewhat when using bivariate data (second panel of Figure 3c): the Akaike weights increase throughout the parameter space, and the bursty regime data move closer to even odds for model selection.

To illustrate the source of the identifiability challenges, we plot the nascent marginals of the models at the two points. In the intermediate regime, the Γ-OU and CIR models yield moderately different distributions, whereas the telegraph model is immediately distinguishable by its bimodality (third panel of Figure 3c). In contrast, in the bursty regime, the distributions are all unimodal and less identifiable (fourth panel of Figure 3c); the Γ-OU and telegraph marginals are particularly similar, as they converge to the same negative binomial limit^20^.

Interestingly, this formulation fully characterizes the effect of certain forms of technical noise. If the transcriptional and observed molecular averages are fixed, but the experiment fails to capture some molecules, the distributions are identical to those obtained by deflating the transcriptional noise intensity. In other words, even though technical noise affects the molecules, its theoretical effects are indistinguishable from decreasing the variability of the transcriptional process. As the noise levels increase, the RNA distributions are pushed toward the indistinguishable Poisson limit at the bottom edge of the reduced parameter space. We quantify how rapidly the information degrades by plotting smaller circles on the first and second panels of Figure 3c to indicate the effect of 50%, 75%, and 85% dropout, in that order from top to bottom.

### 4.4 Distributions obtained from a transient process

Due to the interest in understanding developmental processes, the characterization of transient process dynamics is a key problem in single-cell analyses. The use of mechanistic models with multimodal data, which we emphasize here, was originally pioneered in the context of the RNA velocity framework, which attempts to exploit the causal relationship between nascent and mature RNA to fit transient processes^86^. However, the implementations proposed so far use relatively simple noise behaviors^59,86,121^, which do not recapitulate the bursty transcription observed in living cells. As discussed in our recent analysis of RNA velocity methods^19^, this leads to us to hold some reservations about the robustness and appropriate interpretation of results obtained by this class of methods.

The inference of transient dynamics from snapshot data is a formidable problem due to a combination of theoretical and practical factors. Most fundamentally, it is not precisely clear what a snapshot *is*: how does a single measurement simultaneously capture the early and late states in a differentiation process? To develop an explanatory model, we take inspiration from the existing work on cyclostationary processes^122,123^, cell cycle ensemble measurement modeling^124–126^, Markov chain occupation measure theory^127–129^, and chemical reactor engineering^105,106^. In the typical stochastic modeling context, we fit count data using stationary distributions *P*(**x**), obtained as the limit lim_*t*→∞_ *P*(**x**, *t*) of a transient distribution. By the ergodic theorem^130–132^, this distribution, when it exists, coincides with the occupation measure 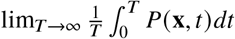, i.e., observations drawn from a single trajectory over a sufficiently long time horizon, rather than from multiple trajectories at once. Conveniently, the ergodic limit has time symmetry with respect to measurement: the distribution does not depend on the timing of the experiment. In the transient case, we cannot take these limits. However, we *can* retain time symmetry by proposing that the experiment samples cells at almost surely finite times *t* since the beginning of the process. Therefore, we conceptualize data as coming from a set of cells indexed by *c*, such that each cell’s time *t*_c_ is sampled from *f*(*t*), and counts are drawn from some distribution *P*(**x**, *t*_c_), which is not typically available in closed form. This formulation yields Equation 32, which requires specifying the distribution *f*.

We illustrate some of the challenges and implications using the model system shown at the bottom of Figure 4a. The underlying transient structure involves transitions through three cell types, each characterized by a particular transcriptional burst size. The transient transcription process produces nascent and mature RNA trajectories for each cell; however, we only obtain a single data point per trajectory. Even if we have perfect information about the cell times, it is far from clear that we can accurately reconstruct the transcriptional dynamics from snapshot data (center of Figure 4a).

In addition, we wish to know whether we can identify the *mechanism* of the snapshot collection. We can imagine cells entering and exiting the observed tissue in multiple ways, which correspond to different choices of *f*(*t*). Some natural choices are uniform, which implies the cells stay in the tissue for a deterministic time^86^; decreasing over time, so cells can exit immediately; or uniform, then decreasing, so cells must stay in the tissue for some duration but are free to leave afterward. These choices can be modeled by Dirac, exponential, and Pareto residence distributions. In the parlance of chemical reactor engineering, these configurations are known as the plug flow reactor, the continuously-stirred tank reactor, and the laminar flow reactor, respectively. Their *f*(*t*), which are the reactor internal-age distributions, are well-known in the chemical engineering literature^105,106^, and shown at the top of Figure 4a. It is not *a priori* obvious the configurations are mutually distinguishable from count data. If they are not, the choice of *f*(*t*) is immaterial for inference.

We generated snapshot data from the Dirac model and fit it under all three models. To efficiently evaluate snapshot distributions, we designed an algorithm which essentially “recycles” *t*_c_ for trapezoidal quadrature. The method is fully described in Section 6.8.4. As shown in Figure 4b, despite only having access to a single observation per time point, all models yield results visually close to the true marginals. However, despite these superficial similarities, quantitative model identification is possible: for the simulated dataset shown, the true Dirac model achieves an Akaike weight of *w*_*ϖ*_ ≈ 79%, whereas the exponential and Pareto both achieve *≈10%. Dec*reasing the dataset size substantially degrades the identifiability (Figure 4c). Even at higher sizes, spread is considerable; for example, a 150-cell dataset gives approximately even odds (*w*_*ϖ*_ > 1 2) on average, but individual realizations vary from confidently correct (*w*_*ϖ*_ ≈ 1) to confidently wrong (*w*_*ϖ*_ ≈ 0).

To understand the robustness of model identifiability, we generated 200 synthetic datasets at random parameter values, constrained to have fairly low expression. We observed poor identifiability, with even or better odds for the correct model in only 20% of the cases (Figure 4d). This performance appears to be attributable to quantitative similarities between all three models’ likelihoods. As shown in Figure 4e, given data of this quality, we cannot even narrow the scope down to two models, as neither of the candidate models performs conspicuously worse than the true Dirac configuration. Therefore, it is possible to fit snapshot data approximately equally well using a variety of models; candidates for *f*(*t*) are identifiable *in principle*, but challenging to distinguish from any particular dataset. This simulated analysis implies that the details of the reactor configuration may not matter much, providing a basis for omitting this model identification problem for real data.

### 4.5 Variability in library construction

To properly interpret single-cell data, we need to exhibit caution regarding the technical noise behaviors and consider multiple possible candidate models. However, before fitting distributions, we must fully characterize the models and understand which of their parameters are actually identifiable with the data at hand. For example, the two-species models explored in Section 4.3 produce distributional forms that are closed under the assumption *p*_*N*_ = *p*_*M*_ = *p*, i.e., the magnitude of the observation probability *p* is impossible to identify from count data alone. Interestingly, when *p*_*N*_ ≠ *p*_*M*_ (that is, when nascent and mature RNA may have different observation probabilities), what we can learn about technical noise heavily depends on the form of the biological noise. For example, under slow transcriptional variation (as in the mixture and Poisson limits of the models explored in Section 4.3), the RNA distributions contain no identifiable information whatsoever about the technical noise, regardless of the amount of data. On the other hand, if transcription is bursty, the distributions depend on the ratio of *p*_*N*_ and *p*_*M*_, but not their absolute values (Section 6.8.5). This theoretical result calls for further investigation: how much information can we obtain *in practice*, given finite data?

To understand the prospects for distinguishing parameters, we consider the simple model system shown in Figure 5a, which involves bursty transcription with average burst size *b*, splicing, degradation, and molecular capture with species-specific probabilities. To characterize how much information about *p*_*M*_/*p*_*N*_ we can identify from count data, we simulated 200 datasets at the ratio values 1/4, 1, and 4, and calculated their likelihoods over (10^−2^, 10^2^). We repeated this analysis using synthetic datasets with 20, 50, 100, and 200 cells, and plotted the average of the posterior distributions for each condition. As shown in Figure 5b, color-coded by the ground truth *p*_*M*_/*p*_*N*_ and intensity-coded by the number of cells, the posteriors are, on average, consistent with the true value. However, even with perfect information about the averages and the nascent RNA distribution, the uncertainty is considerable; at larger dataset sizes, we can typically localize the ratio to an order of magnitude, but not much further.

**Figure 5.**
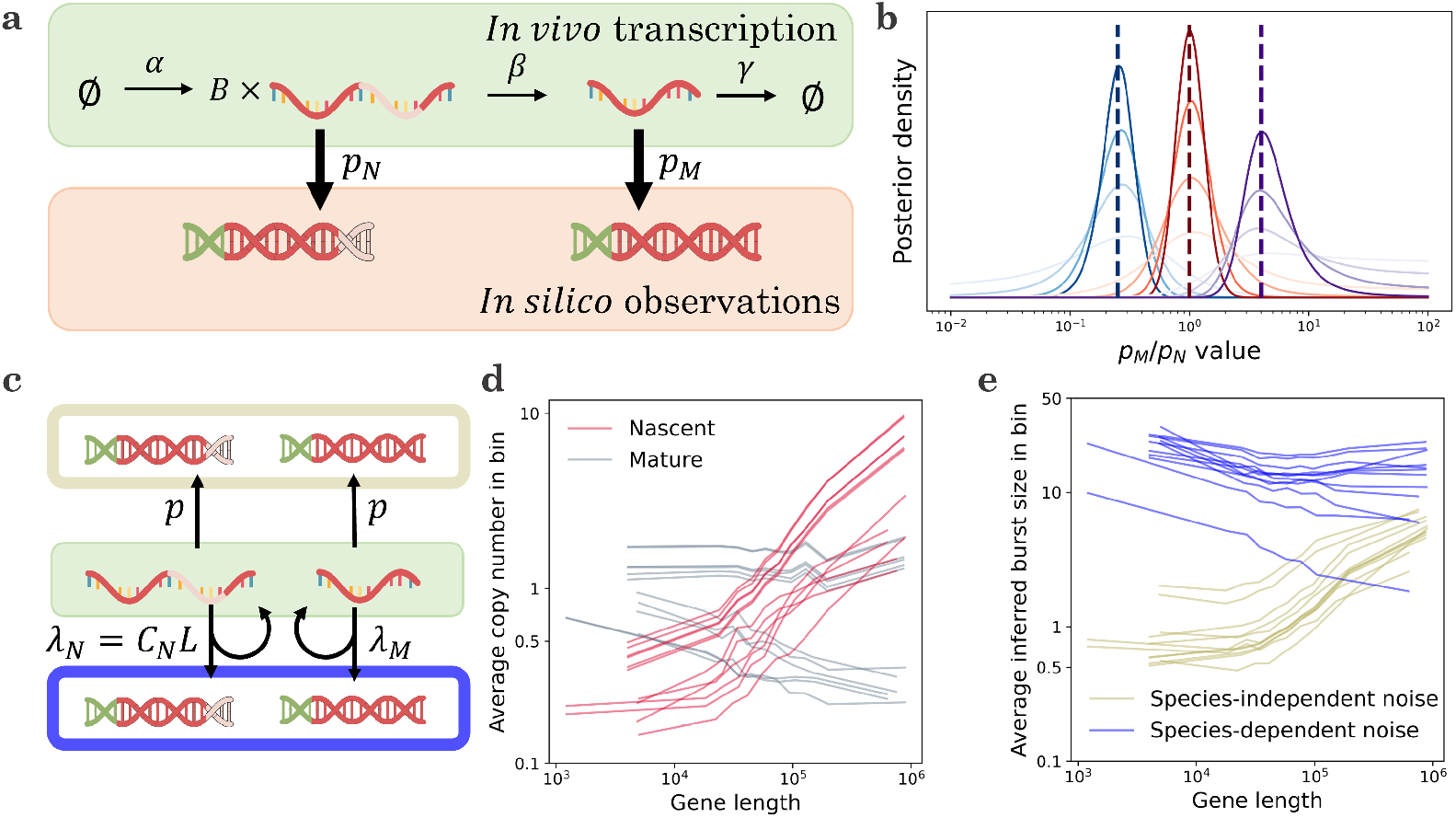
Technical noise models may be identified from count data, either by direct application of statistics or by imposing informal priors about the biological variability. **a**. A minimal model that accounts for non-homogeneous noise: transcriptional events occur with frequency *α*, generating geometrically-distributed bursts *B* with mean size *b*; the molecules are spliced with rate *β* and degraded with rate *γ*. Nascent molecules are observed with probability *p*_*N*_ and mature molecules are observed with probability *p*_*M*_. **b**. Given information about the nascent distribution and the mature mean, it is possible to use joint distributions to obtain information about the ratio of observation probabilities (curves: average posterior likelihoods, computed from 200 independent synthetic datasets; color: true value of *p*_*M*_/*p*_*N*_, blue: 1/4, red: 1, purple: 4; dashed lines: location of each true value; color intensity: from lightest to darkest, synthetic datasets with 20, 50, 100, and 200 cells). **c**. Two models considered in Gorin et al.^21^: the species-independent bias model for length dependence in averages, which proposes that nascent and mature RNA are sampled with equal probabilities, and the species-dependent bias model, which proposes that the nascent RNA sampling rate scales with length (top, gold: kinetics of species-independent model; bottom, blue: kinetics of species-dependent model; center, green: the source RNA molecules used to template cDNA). **d**. A variety of single-cell datasets produce consistent and counterintuitive length-dependent trends in nascent RNA observations (lines: average per-species gene expression, binned by gene length; red: nascent RNA observations; gray: mature RNA statistics; data for 2,500 genes analyzed in Gorin et al.^21^). **e**. Fits to the species-independent model show a strong positive gene length dependence for inferred burst sizes, whereas fits to the species-dependent model show a modest negative gene length dependence, which is more coherent with orthogonal data (lines: average per-gene burst size inferred by *Monod*^133^, binned by gene length; gold: results for species-independent model; blue: results for species-dependent model; data for genes analyzed in Gorin et al.^21^ after goodness-of-fit).

Given the statistical challenges illustrated by simulations, we speculate that it may be more fruitful to use prior information about biology and physical intuition about sequencing to construct technical noise models. For example, in a recent paper^21^, we fit models that represent two competing hypotheses (Figure 5c). The first has identical, gene-specific observation probabilities *p* for the nascent and mature species. In this model, the inferred burst size is *b p*, as these two parameters are not mutually identifiable. The second has a gene length-dependent technical noise term for the nascent species, which coarsely represents a higher rate of priming for long molecules with abundant intronic poly(A) tracts, and a shared genome-wide term for the mature species, which represents priming at the poly(A) tail. In this model, the inferred burst size is *b*.

These models attempt to explain the trend summarized in Figure 5d: across a wide range of datasets, nascent RNA averages exhibit a pronounced length dependence^134^ not evident in mature RNA^135^. The first model explains the trend by a species-independent bias, as *b* and *p* control nascent as well as mature RNA levels. Conversely, the second model explains it by a species-dependent bias. Both models produce fair fits to the data (as demonstrated, e.g., by the low rate of rejection by goodness-of-fit in Sections S7.4 and S7.5.2 of Gorin et al.^21^).

However, the trends in the resulting inferred parameters are strikingly different: the species-independent bias model predicts that longer genes have higher *b p*. Ascribing this trend to the *b* term—longer genes have higher burst sizes— contradicts burst size trends from fluorescence microscopy^136^. Ascribing it to the *p* term—longer genes have higher sampling probabilities—is physically unrealistic, because mature RNA molecules are depleted of the internal poly(A) tracts necessary for priming^137^. On the other hand, the species-dependent model predicts a modest negative relationship between length and burst size, which is more coherent with orthogonal data.

This technical noise model is a relatively simplistic low-order approximation, since all genes have the same mature molecule capture rate *λ*_*M*_ and length scaling *C*_*N*_. Nevertheless, it foregrounds a key modeling principle of the investigation: in the absence of prior information, biological parameters need to be fit on a gene-by-gene basis, but technical noise should be constructed using a common genome-wide model that varies in a mechanistic rather than arbitrary way. In sum, the mathematics enable us to define and fit systems, but to understand whether the fits are sensible, we need to contextualize and compare them with previous results and physical intuition.

## 5 DISCUSSION

The results we have derived provide a blueprint for the holistic modeling of single-cell biology and sequencing experiments. First, we have outlined a generic mathematical framework for treating stochasticity in living cells. By exploiting the generating function representation, we reduce discrete, continuous, and mixed reactions to operators in a system of differential equations. These ODEs can be straightforwardly solved via numerical integration to compute model properties, including likelihoods. This approach recapitulates and subsumes a wide range of previous results^16,17,20,21,75,76,85,100,138,139^.

By treating the discrete and continuous degrees of freedom on equal footing, our approach makes certain otherwise challenging problems straightforward to solve, as illustrated in Section 6.8.1. By making simplifying assumptions—chiefly, the assumption of independent and identically distributed sampling—we reduce the modeling of technical variation to the composition of generating functions. Our framework may be used in its current form, or as a substrate for developing more sophisticated models of transcriptional regulation and sequencing that subsume it in turn. This process simply involves instantiating hypotheses, converting them into probabilistic models, and constructing model solutions using a procedure analogous to the one presented in Figure 1c.

We believe this framework comprises a productive vision for the interpretation of large datasets, but many technological and mathematical challenges remain. For example, the library construction biases are dependent on molecule-specific factors that we do not yet fully understand, because their effect is heavily convolved with biological variability. In Figure 5, we considered two extreme cases, where the noise strength/length scaling is either unconstrained or forced to be identical for all genes. We anticipate that careful investigation of technical biases will be necessary to construct models that constrain the technical biases based on RNA chemistry, while allowing for gene-to-gene and droplet-to-droplet variability.

In Section 6.7 and supplemental information, we discuss the challenges associated with modeling ambiguous species, motivated by the limitations of short-read sequencing for distinguishing between spliced and unspliced forms of the same RNA gene product^140^. It is worth noting that even the spliced/unspliced binary is a convenient simplification primarily adopted because of data availability^86,113^; we stress that a truly comprehensive treatment requires defining intermediate states^19^, their relationships, and their mutually indistinguishable classes. These computational foundations do not yet exist, although we have attempted a partial solution in recent work^16^ and outlined some promising directions in supplemental information. Therefore, despite our immediate interest in bivariate RNA distributions, our framework is designed to generalize to other modalities as they become practical to quantify.

The full generating function solutions we have outlined here are typically not computable directly. By construction, the generating function needs to be evaluated on a grid; Fourier inversion produces a grid of microstate probabilities, which needs to be quite large to avoid artifacts^139^. If the grid dimension is 𝔰_*i*_ for each discrete species *i*, the overall state space size is 𝔰 = *N* Π^*i*^ 𝔰_*i*_. Even in the simplest case, where we only quantify and fit discrete counts, evaluating the probability mass function requires storing and inverting an *n*-dimensional array, which usually has size 𝔰 far too large to be practical (e.g., Fig. S5b of our prior work on bursty models^16^).

When applicable, the generating function approach has numerical advantages over the stochastic simulation algorithm (SSA)^141–143^, which approximates distributions by the empirical distributions of trajectories, and finite state projection (FSP)^78^, which directly integrates a version of the master equation confined to a finite 𝔰. Specifically, if we only care about a particular species *i*, we can evaluate its marginal using a grid of size *N*𝔰_*i*_ with 𝔰 *log* 𝔰 time complexity. In the worst-case scenario, FSP requires a grid of size 𝔰 with 𝔰^3^ time complexity, as evaluating a particular marginal requires explicitly evaluating the probabilities for the entire grid, then marginalizing. Similarly, SSA requires explicitly simulating the entire system to obtain the marginals, and has the drawback of the usual inverse square root Monte Carlo convergence^144,145^. In addition, FSP is not compatible with the generating function manipulations used to represent technical noise, SSA is relatively challenging to adapt to time-dependent rates^146^, and neither FSP not SSA is readily compatible with continuous stochastic processes (although exact^20^ and approximate^20,147,148^ hybrid schema can be constructed with some work). In the future, the “curse of dimensionality”— the reliance on grid evaluation—may be possible to bypass altogether by training neural networks to predict probability distributions, but this approach is as of yet in its nascence^149–152^ and will require considerable further development to apply to general systems.

Nevertheless, SSA and FSP are substantially more general than the approach we outline here. The simulation- and matrix-based methods only require a list of reactions, whereas the generating function methods also require those reactions to produce readily solvable partial differential equations. We have omitted phenomena which would be trivial to treat using FSP and SSA, such as regulation involving feedback. To our knowledge, these phenomena, which are mathematically analogous to adding multi-molecular interaction terms, cannot be directly treated with the method of characteristics. Instead, they require perturbative methods^77^ or fairly complicated special function manipulations^101–104^, which do not easily generalize. We illustrate the challenges in supplemental information, using the example of downstream species catalyzing gene state transitions.

We have, until now, stressed applications to “snapshot” single-cell data from dissociated tissues; however, our framework may be extended to spatial single-cell data; for instance, we can define transcriptional parameters that depend on the cell’s coordinates in the tissue. In this case, the typical systems biology goals translate to fitting a time- and space-dependent function that governs these parameters. However, the generating function formulation relies on the assumption of cells being stochastically independent; it is far from clear that this should hold for densely sampled spatial data, and more sophisticated alternatives, such as agent-based models, may be needed^153,154^.

Despite these challenges, the framework is already quantitatively useful. To fully “explain” a dataset, we need to fit gene-specific transcriptional mechanisms, genome-wide technical noise and co-expression parameters, and cell type structure, while controlling for potential misspecification. However, at this time, it may be more fruitful to focus on narrower questions, using assumptions, orthogonal data, or simulated benchmarking to justify omitting some parts of the problem^19^. We have applied this “bottom-up” approach to single-cell data, considering, in turn, the estimation of transcriptional kinetics and technical noise^21,133^, the identification of transcriptional models^20^, the analysis of co-regulation patterns^16^, and the determination of nuclear transport kinetics^155^. Conversely, it may be valuable to apply a “top-down” approach, augmenting an existing method with biophysically meaningful noise, as we have proposed in the context of transient processes^19^ and neural network dimensionality reduction^71^.

We anticipate that making meaningful progress on the stochastic modeling project championed by Wilkinson will require extended “real contact”^156^ between systems biology, genomics, and mathematics. The general framework we propose, which unifies a variety of previous work, represents one step towards this synthesis. The role of mathematics here is key; as Wilkinson noted, the stochastic systems biology of single cells cannot be “properly understood” without stochastic mathematical models.

## DATA AVAILABILITY

Notebooks that reproduce all of the results in the figures are hosted at https://github.com/pachterlab/GVP_2023. The raw data used to generate Figure 2b-c, as well as related supplementary figures, are hosted as the Zenodo package 7694182. The data and *Monod* fits reported in Figure 5d-e, originating from Gorin et al.^21^, are hosted as the Zenodo package 7388133, and were originally generated using the notebooks and scripts at https://github.com/pachterlab/GP_2021_3/.

## ACKNOWLEDGMENTS

G.G. and L.P. were partially funded by NIH 5UM1HG012077-02 and NIH U19MH114830. J.V. was partially funded by NIH 1U19NS118246-01. The RNA, DNA, and cDNA illustrations were derived from the DNA Twemoji by Twitter, Inc., used under the CC-BY 4.0 license. The authors thank Dr. A. Sina Booeshaghi, Maria Carilli, Tara Chari, Taleen Dilanyan, Dr. Kristján Eldjárn Hjörleifsson, Meichen Fang, Catherine Felce, and Delaney Sullivan for fruitful discussions of co-regulation, contamination, transient behaviors, catalysis, fragmentation, genomic alignment, and a variety of other phenomena and processes. Part of this work was performed during G.G.’s Data Sciences Co-op with Celsius Therapeutics, Inc.

## 6 METHODS

### 6.1 Master equation models of transcription

We are interested in continuous-time stochastic processes that combine categorical, nonnegative discrete, and (usually nonnegative) continuous degrees of freedom. To solve these systems, we begin by separately defining their allowed transitions and converting them to master equation forms.

The categorical variable, denoted by *s* ∈ {*1*, ⋯, *N*}, represents the instantaneous state of a multi-state gene. By assuming that the state interconversions are Markovian and independent of all other components of the system, we can define *H*_*ij*_, the rates of transitioning from state *i* to state *j* :

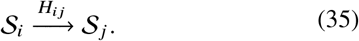

These rates can be summarized in the state transition matrix 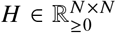, such that *H*_*ii*_ = − Σ_*j≠*] *i*_ *H*_*i j*_ and Σ_*j*_ *H*_*i j*_ = 0 to enforce the conservation of probability. This set of transitions can be represented by a master equation involving finitely many ODEs, which tracks the probabilities of each state *s* at a time *t*:

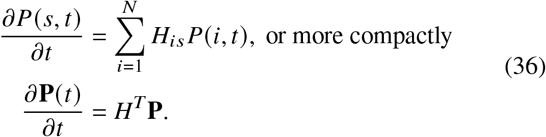

As this system is expressed in terms of a differential equation for an arbitrary time *t*, the relation holds for time-dependent *H*. For simplicity, we assume that *H* is deterministic.

The nonnegative discrete variables, denoted by 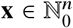, represent molecular copy numbers. We assume that *n* molecular species participate in four classes of transitions, and can summarize their effect by considering their reaction schema and effect on *x*_*i*_, the number of molecules of species *i*:

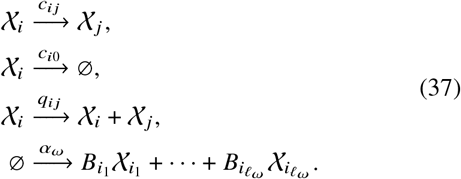

First, species *i* can be converted to species *j* with rate *c*_*ij*_ *x*_*i*_. Second, species *i* can spontaneously degrade with rate *c*_*i*0_*x*_*i*_. These classes of monomolecular transitions, which either maintain or reduce the total number of molecules in the system, can be summarized in the matrix *C*^*dd*^ ∈ ℝ^*n*×*n*^, such that 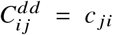 and 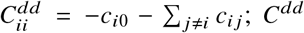; is the matrix governing the associated reaction rate equations^17,85^. Third, species *i* participate in autocatalysis at the rate *q*_*ii*_, or catalysis of species *j* at the rate *q*_*ij*_. These reactions can be summarized by the matrix 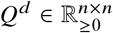, such that 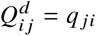. Finally, molecules can be produced. In the general case, a burst of production simultaneously creates molecules of *ℓ*_*ω*_ discrete species 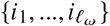. We assume bursts are described by a Poisson arrival process, with burst frequency 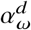 and the nontrivial *ℓ*_*ω*_-variate joint distribution 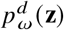 of non-negative burst sizes 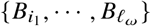. This formulation includes the trivial case of Poisson point process production of species *i*, for which *ℓ*_*ω*_ = 1 and 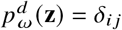, the degenerate distribution located at unity for species *i* and zero for all other species.

This mass action model, which tracks molecule counts, can be represented by an equivalent discrete chemical master equation, which tracks the probability of each microstate **x**:

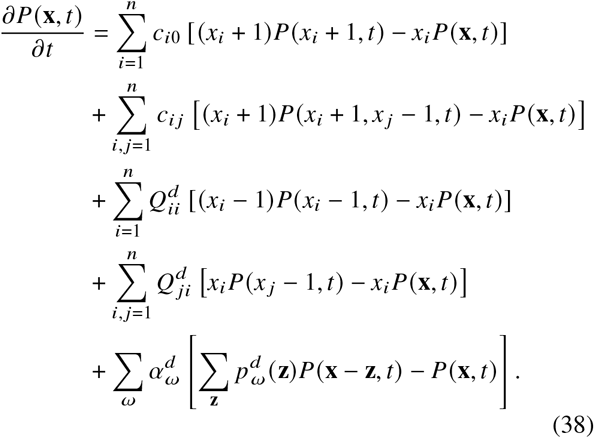

For simplicity of notation, species that do not occur in a reaction are elided from the master equation, as in previous work on modeling bursty transcription^16^. As above, this equation holds even if the rates are time-dependent. For the purposes of this report, we assume only *α*_*ω*_ and *p*_*ω*_ can vary over time.

The nonnegative continuous variables, denoted by 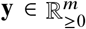, represent concentrations or coarsely-modeled noise sources. We assume that these variables are governed by Ornstein–Uhlenbeck-type stochastic differential equations:

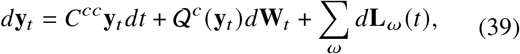

where **y**_*t*_ is a realization of the process, **W**_*t*_ is an w-dimensional Brownian motion, and **L**_*ω*_ is a subordinator. The matrix *C*^cc^ ∈ ℝ^m×m^ sets the mean-reversion terms, whereas the operator 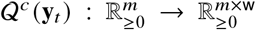 sets the level of noise. We assume that each **L**_*ω*_ only includes drift or compound Poisson terms. The drift terms have the form 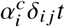. To slightly lighten the notation, we can aggregate all drift terms under *ω* = 1, ⋯, *m* as 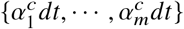; some of these entries may be zero. The compound Poisson terms have the form 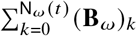, such that N_*ω*_(*t*) is a Poisson random variable with mean 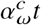 and (**B**_*ω*_)_*K*_ is a set of independent and identically distributed realizations of the random variable **B**_*ω*_. This random variable has a nontrivial *ℓ*_*ω*_-variate joint density 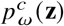on 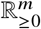, with the remaining *m* −*ℓ*_*ω*_ dimensions concentrated at zero. We note that this formulation entails a slight abuse of notation, as *ω* is used to index over discrete burst processes as well as continuous drift and jump components.

For simplicity, we assume the noise term takes the form of an uncoupled square-root diffusion, such that w = **m** and 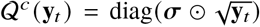. The symbol ⊙ *denotes* the elementwise/Hadamard product of two vectors, the square root should be interpreted as elementwise, and all elements of the constant volatility vector *σ* are non-negative. Although this choice of 𝒬^*c*^ is somewhat restrictive, it produces a particularly simple diffusion tensor Σ:

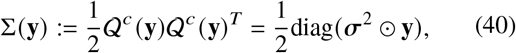

where the square *σ*^2^ should be interpreted as elementwise. This formulation can be reframed as a Fokker-Planck equation^158^, which tracks the probability density of each microstate **y**:

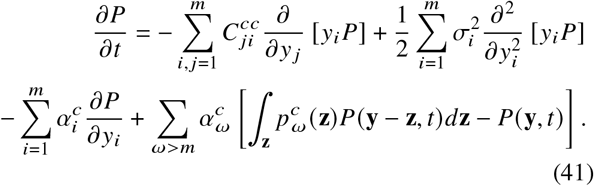

As above, we assume that only the components of **L**_*ω*_ can vary in time.

In addition to these discrete- and continuous-only terms, we need to account for these components’ interactions. For example, we may want to represent the production of a discrete species controlled by a continuous variable, e.g., a time-varying transcription rate^20^:

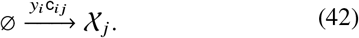

This reaction has the rate *y*_*i*_c_*ij*_. This class of reactions can be summarized in the matrix 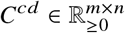, such that 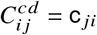. In other words, this class of reactions contributes the following terms to the overall master equation:

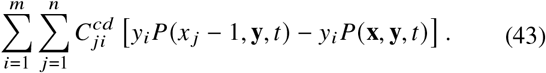

Finally, we may want to represent the production of a continuous species from a discrete one, e.g., the rapid translation of high-abundance protein from low-abundance RNA^139^. This class of reactions simply adds a term proportional to 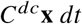 to the expression for **y**_*t*_. The matrix 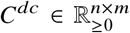contains the relevant rates, such that 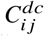 is the rate of producing the continuous species *i* from discrete species *j*. Therefore, we append a set of drift-like terms to the Fokker-Planck equation:

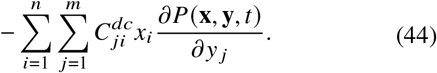

To construct the full master equation, we need to define a system of *N* coupled equations. To do so, we essentially add Equations 36, 38, 41, 43, and 44, replacing all instances of *P* with **P**(*s*, **x, y**, *t*). However, to account for differences in transcription between gene states, we allow the *ω*-associated terms to vary with *s*. The full master equation is reported below in Equation 45.

### 6.2 The full master equation

The full master equation for *P*(*s*, **x, y**, *t*) is:

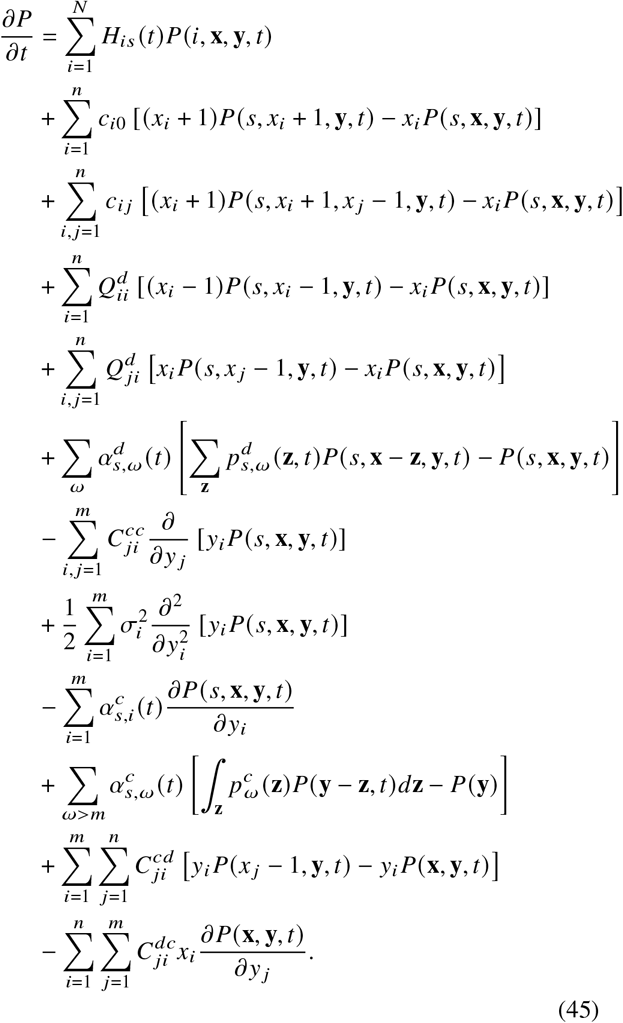

We annotate the terms in Table S1.

### 6.3 Generating function methods for biological stochasticity

The full master equation is fairly cumbersome and challenging to analyze directly. Therefore, analysis has to proceed by spectral methods. We use the generating function (GF), a length-*N* vector function **G**, such that each component is

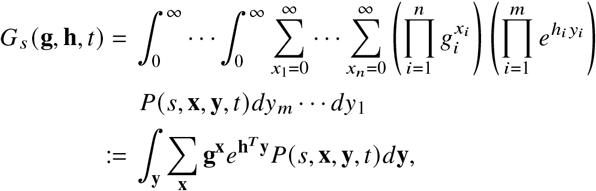

where the lowest line is the definition expressed in useful shorthand notation. Formally, the generating function is the combination of a probability-generating function (PGF) in the discrete variables and moment-generating function (MGF) in the continuous variables. The arguments **g** ∈ ℂ^*n*^ and **h** ∈ ℂ^m^ are spectral variables. By computing the generating function of both sides of Equation 45, we find (see supplemental information) that the master equation is equivalent to a much more compact system of partial differential equations:

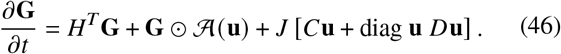

This formulation relies on defining the unified variables **u**:

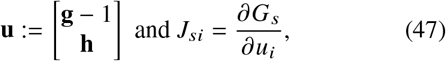

as well as unified matrices:

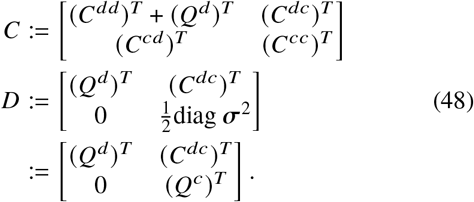

Each entry of the length-*N* matrix function 𝒜consists of the burst and drift terms:

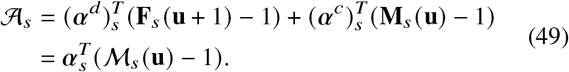

The vector 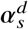 contains the frequencies of all discrete burst processes for state *s*. The first *m* entries of 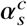 contain the continuous species’ drifts in state *s*. The remaining entries contain the corresponding rates of continuous burst processes. ***α***_*s*_ aggregates these quantities. The vector function **F**_*s*_ contains the joint PGF of the discrete burst processes, and only depends on the first *n* variables. The vector function **M**_*s*_ contains the drift terms, as well as the joint MGF of the continuous burst processes, and only depends on the last *m* variables. The parameters of the 𝒜_*s*_ operator may vary in time.

To obtain the generating function at *t*, we apply the method of characteristics. First, we calculate the characteristics parametrized by the scalar variable s:

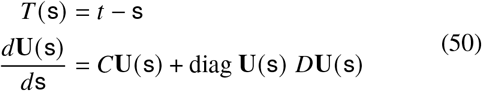

where **U** (s = 0)= **u**. This is the “downstream” ODE, which governs abundances in isolation from production and regulation.

Therefore, **G** is governed by the following system of ordinary differential equations:

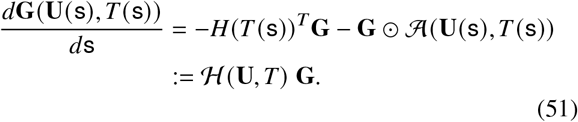

To obtain **G** at *t*, we integrate this matrix system from s = *t* to s = 0. We use **G**^0^(**U**(*t*)) as the initial condition, where **G**^0^ is the generating function of the initial distribution. This is the “upstream” ODE, which governs the full generating function.

In the general case, evaluating this system requires two applications of quadrature: first, solving the *n* + m-dimensional downstream system to obtain the values of characteristics **U** at a set of grid points over [0, *t*]; then, solving the *N*-dimensional upstream system to obtain the value of the generating function.

Some special cases afford simpler solutions. If *D* ≠ 0, the downstream ODE takes a Riccati-like form and generally resists exact analysis^17,159^. However, if *D* = 0 and *C* is diagonalizable, the system takes the tractable linear form

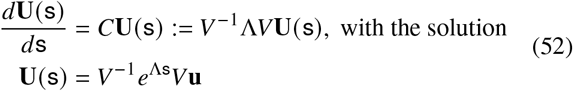

whenever all eigenvalues of *C* are distinct. When they are not, the ODE can be solved in a similar way using generalized eigenvectors. Practically, this means that only one application of quadrature is required.

If, in addition, *N* = 1, the upstream ODE reduces to a single integral:

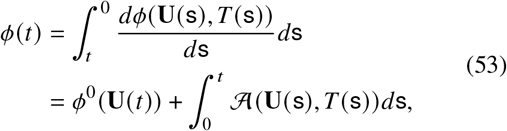

where *ϕ* := log *G, ϕ*^0^ = log *G*^0^, and the generating function *G* is no longer boldfaced because only a single gene state exists.

If 𝒜 is a linear operator *a*_1_*u*_1_ + ⋯ + *a*_*n*+m_*u*_*n*+m_, the system is in the drift-only regime; no bursting occurs. In this case, the system reduces to

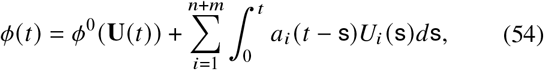

where *U*_*i*_ are the components of **U**. As each *U*_*i*_ is, in turn, a weighted sum of *u*_*i*_, the second term of the log-generating function is given by a sum of fairly simple convolutions that scale as 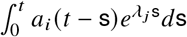.

Finally, in the simplest case, if all eigenvalues *λ*_*i*_ of *C* are negative, the transient part of Equation 54 vanishes as *t* → ∞ and the stationary log-generating function is a linear combination of *u*_*i*_. This implies that the distribution converges to a product of independent Poisson distributions^17,85^.

### 6.4 Coupling multiple genes

The results solve master equations with abstracted production and processing reactions. To connect them to systems phenomena, such as the co-regulation of multiple genes, we need to specify how upstream interactions lead to co-expression. As the simplest illustrative model system, we can consider the coregulation of two genes, indexed by *j*, with 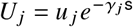. We outline several relatively simple classes of candidate models which induce expression coupling.

In the simplest case, ℋ (**u**, *t*) = Σ_*j*_ ℋ_*j*_(*u*_*j*_, *t*). In other words, the genes’ dynamics are fully separable, and produce solutions in the form *G*(**u**, *t*) = Π_*j*_ *G*_*j*_(*u*_*j*_, *t*). This formulation produces independent distributions at each *t*, but the *trajectories* may possess nontrivial statistical relationships. For example, if both genes start at *x*_1_ = *x*_2_ = 0, their trajectories will be correlated over a finite timespan [0, *T*], with the correlation decaying as *T →* ∞.

In the next simplest case, co-regulation is the consequence of parameter differences in subpopulations. For example, the full cell population may consist of cell types indexed by *κ*. If we suppose each cell type has the abundance π_*κ*_ and transcriptional parameters Θ_*κ*_, we obtain

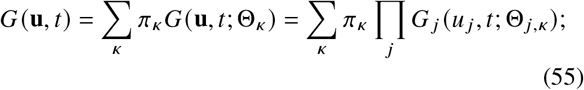

i.e., the generating function decomposes into a product of independent generating functions *conditional on* a particular cell type, but not globally. In other words, even if transcriptional processes are independent, cell type structure can produce nontrivial relationships between genes.

Alternatively, we can propose a model of co-regulation by the categorical variables. For example, two neighboring genes may prefer to have the same or opposite accessibility, depending on the polymeric properties of DNA. Assuming, for the purposes of illustration, that the system is symmetric, we obtain the following *N* = 4 form:

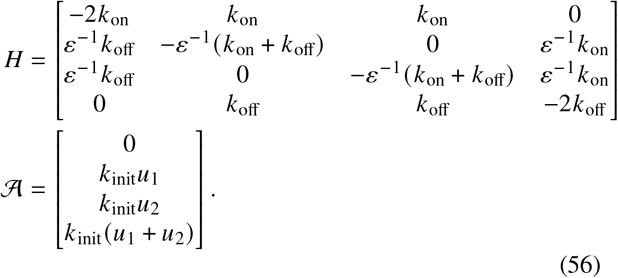

This form encodes the co-regulation of two genes, such that *s* ∈ {both off, gene 1 on, gene 2 on, both on}. If *ε* ≪ 1, the intermediate states are unstable and the genes tend to be either both on or both off. If *ε* ≫ 1, the intermediate states are particularly stable, and only one of the genes tends to be on at a time. If *ε* = 1, we recover the independent case.

We can define a similar model for co-regulation by a continuous variable *y*_1_. For example, there may be a latent regulator, such as the concentration of an activator, that controls multiple loci: if it is high, both have a high transcription rate; otherwise, both are inactive^20^. This amounts to appending the following reactions to the master equation:

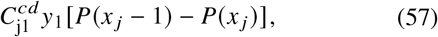

where the *C*^c]^ matrix encodes the relationship between the concentration and the transcription rate. Therefore, the genes become mutually correlated through the trajectory of *y*_1_, although the extent of correlation depends on the dynamics. If the categorical or continuous driving process is bursty, we can approximate it by a co-bursting module. For example, in the limit of *ε* → 0, the dynamics of the system in Equation 56 converge to the *N* = 2 formulation

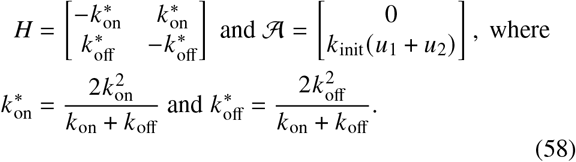

If, in addition, 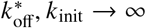, we obtain the *N* = 1 module characterized by

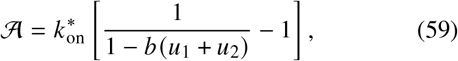

where 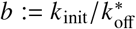^16^. This is the bursty limit of Equation 56. Interestingly, that mechanism also possesses a slow mixture limit. If *ε* → ∞ while *k*_*o*n_, *K*_off_ → 0, we obtain a special case of Equation 55, with *τ*_*K*_ = 1/2 and mutually exclusive expression in the “cell types,” or long-lived gene states.

Even when we restrict our analysis to simple feed-forward regulation, this outline of motifs is nowhere near exhaustive. Nevertheless, the “mixture” and “bursty” limits are particularly natural starting points, as their distributions are straightforward to construct. In other words, we speculate that the careful analysis of co-expression models can distinguish relationships due to “slow” variation between cell types and “fast” variation due to coupled transcriptional events.

### 6.5 Transient phenomena

This result yields a fairly simple numerical recipe for the determination of probabilities at a particular time *t*. Typically, analysis proceeds by assuming *H* and 𝒜 are time-independent and letting *t* → ∞, i.e., considering the stationary limit of the process. However, this may not be strictly justifiable: much of single-cell analysis involves the determination of trajectories from intrinsically transient data representing differentiation pathways^160^ If the transient process occurs on a timescale comparable to RNA turnover, using a stationary model may not be appropriate^16^.

To rigorously fit transient data, we need to posit just *how* a snapshot of cells may capture multiple cell states, such that some states are the progenitors of others. The solution is not yet clear, and multiple reasonable explanations exist; for example, we may suppose that the differentiation process “lags” in certain cells (in the vein of the models of variability proposed in Stumpf et al.^44^ for development, and in Sanders et al.^161^ and Perez-Carrasco et al.^125^ for the cell cycle). In other words, all cells are captured at a time *t* since the beginning of a process, but *H* and 𝒜 have different time-dependence for different cells. Although such an explanatory model can be instantiated, it may be too challenging to fit. Further, it does not appear to be compatible with processes that operate continuously; the choice of *t* becomes somewhat challenging to motivate.

We propose that the simplest model for observations relies on minimal synchronization between the biology and the experimental process. To mathematically formalize it, we take inspiration from the theory of reactor modeling in chemical engineering^105^ and extend preliminary work from our recent RNA velocity methods analysis^19^. A cell enters a medium; this entrance triggers a chemical signal that begins a transient process. The dynamics of this transient process are only dependent on time since receiving the signal, and identical between cells. After a delay, the cells exit the medium. In this framework, sequencing is the uniform random sampling of cells present within this medium. Although this formulation is admittedly simplistic—it excludes the cell cycle and stochastic driving—it allows us to take the first steps with a systematic study of using snapshot data to fit transient stochastic processes. This toy model is numerically tractable, which is useful for its simulation and characterization, and possesses a stationary state that is independent of the time at which the experiment is performed, which is useful for biological admissibility and realism.

Therefore, to marginalize over *t*, we need to augment the model with an additional property: the relationship between time along a transient process and the probability of capturing a cell. In the parlance of reactor engineering, this relationship is given by the internal-age distribution *f*. The simulations of transient processes in La Manno et al.^86^ and Bergen et al.^59^ implicitly adopt this model and assume a particular functional form of *f*. We might suppose cells enter the observation window at *t* = 0 and leave it at *t* = *T*, with a Dirac residence time distribution *δ*(*t* − *T*) and uniform sampling throughout this window. The resulting age distribution is uniform, with *f* = *T*^−1^, and formally corresponds to the ideal plug flow reactor (PFR) architecture^105^. As *T* → ∞, we obtain the *t* → ∞ ergodic limit, if such a limit exists. On the other hand, if *f→δ*(*t* −*T*), we recover the instantaneous distribution at time *T*; this limit formally corresponds to the batch reactor (BR).

To obtain the generating function for the cells inside a tissue, we represent the tissue as a reactor, specify its influx and efflux properties, and solve for the internal-age distribution *f*. This internal-age distribution yields the occupation measure of the process times, as discussed in our RNA velocity review^19^, and induces the following reactor-wide generating function:

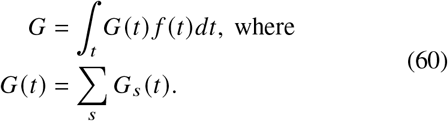

We have marginalized over the instantaneous gene state *s* because this variable is typically not observable.

### 6.6 Droplet encapsulation noise

The generating function *G* describes the biological variability due to molecular processes, transcriptional driving, and the capture of cells from a reaction medium. However, single-cell RNA sequencing data do not quantify cells—they quantify *barcodes*. Cells are randomly encapsulated into droplets with barcoded beads; to avoid the formation of “doublets,” with two cells per droplet, the microfluidic protocols typically have a fairly low encapsulation rate. If we assume that a droplet may have either zero or one cells, we obtain the following generating function for the distribution of RNA on a per-barcode level:

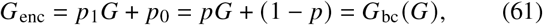

where *G*_bc_ is the PGF of the Bernoulli distribution, with *p*_1_ = *p* the probability of capturing a single cell and *p*_0_ = 1− *p* that of capturing none. Analogously, if we assume that doublets can occur, and the encapsulation of cells is independent and identically distributed (i.i.d.), we find

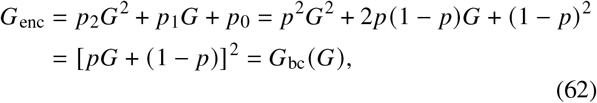

where *G*_bc_ is now the PGF of the binomial distribution. It is straightforward to extend this to the unconstrained case, with per-cell encapsulation *rate λ*, and obtain the analogous expression

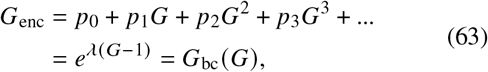

where *G*_bc_ is the PGF of the Poisson distribution.

However, even empty droplets typically contain some “background” molecules. Removing the empty droplets by filtering for cells with relatively high expression, as well as correcting for the background, is a standard part of sequencing workflows^57,109–112^. To model the joint distribution of biological and background RNA, we need to instantiate a mechanistic hypothesis about its source. The simplest hypothesis consists of two parts. First, we impose the *pseudobulk* interpretation of background: we assume that a fraction of the cells loaded in the library construction step are lysed, and produce a pool of loose molecules. Next, we assume that these molecules are free to be encapsulated into the droplets in an i.i.d. fashion. This implies the Poisson functional form for the distribution of debris entering each droplet:

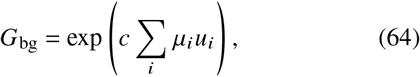

where c is some shared constant that reflects the pool size and the rate of diffusion, whereas 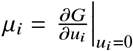 is the expectation of species *i* over the entire cell population. This simplest model assumes that all cells are equally likely to lyse and release their contents; if this assumption is violated, *μ*_*i*_ needs to be obtained by computing an expectation with respect to a measure biased toward the less stable cells. Finally, the full per-droplet distribution of molecules is

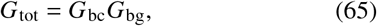

i.e., each droplet contains contributions from the encapsulated cells, as well as the background. With some abuse of notation, we occasionally use the expression *G*_bc_(*G*) *G*_bg_(*G*), where the first argument denotes composition, whereas the second denotes functional dependence.

### 6.7 Library construction and sequencing noise

We cannot observe the biological molecule content of each droplet: we are restricted to analyzing counts of complementary DNA (cDNA). In a typical dual-index 3’ microfluidic workflow (e.g., the commercialized 10x chemistry^48^), these cDNA are quantified by the following sequence of reactions. First, a synthetic primer captures a poly(A) stretch in RNA, which may be an endogenous molecule or a synthetic tag^162^. The primer contains a poly(dT) oligonucleotide, a sequencing primer, a cell barcode, and a unique molecular identifier (UMI). Next, reverse transcriptase (RTase) attaches to the RNA-primer complex and synthesizes the complementary strand. When the first strand is complete, a template-switching oligonucleotide (TSO) attaches to the end, allowing RT to synthesize the second strand of cDNA. After library construction, the droplet emulsion is broken, producing a pool of long cDNA; polymerase chain reaction (PCR) is used to amplify this pool. The long cDNA molecules are enzymatically fragmented, and another sequencing primer is attached at the end of the molecule that formerly contained the TSO. Finally, another round of PCR amplifies the pool and appends sample indices and Illumina adaptors to both sides of the molecule. The pool of cDNA is loaded onto a sequencing machine and sequenced from both sides, producing two reads. One read contains the barcode and UMI bases, whereas the other contains partial information about the 3’ end of the molecule, beginning at the fragmentation site. This sequence of reactions represents the ideal-case scenario, and the products may well include artifacts due to off-target reactions^163^.

To understand the effect of technical variability on the per-barcode distributions, we need to summarize this workflow in a mechanistic model. First, we assume that the library preparation reactions occur in an i.i.d. fashion relative to each RNA molecule in the droplet, allowing us to construct a separate description of technical noise for each discrete molecular species indexed by *i*. At this stage, we omit the modeling of continuous species. As we quantify the number of UMIs, we can considerably simplify the description by splitting the workflow into the initial cDNA synthesis and all downstream steps. For the cDNA synthesis, we may choose one of two models:

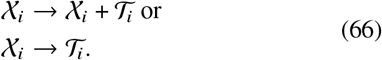

In the first model, the formation of a UMI-tagged cDNA 𝒯_*i*_ is non-sequestering, and the template RNA 𝒳_*i*_ can participate in further cDNA synthesis. In other words, a single RNA molecule can produce more than one cDNA with distinct UMIs. In the second model, the cDNA synthesis is sequestering, and each RNA can template at most one cDNA with a particular UMI. For the downstream steps, if we assume the PCR and sequencing steps produce results that are reasonably faithful to their templates, we are essentially restricted to a single model:

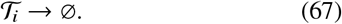

In other words, the sequence of steps after the formation of cDNA 𝒯_*i*_ may lose some UMIs, but it cannot create them. Aggregating these steps, we find the shifted per-molecule generating function for technical noise:

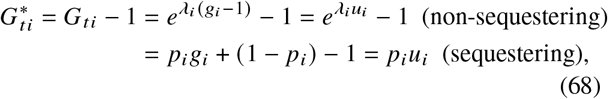

where *λ*_*i*_ = *λ*_*i*,c_ *p*_*i,p*_ and *p*_*i*_ = *p*_*i*,c_ *p*_*i,p*_. *λ*_*i*,c_ is the overall Poisson rate of the catalytic production of cDNA 𝒯_*i*_ with distinct UMIs, *p*_*i*,c_ is the probability of producing a single cDNA 𝒯_*i*_ in a non-catalytic fashion, and *p*_*i,p*_ is the probability of retaining a molecule of 𝒯_*i*_ through the PCR steps. It is straightforward to use a Taylor expansion to observe that the limit *λ*_*i*,c_ ≪ 1 yields the Bernoulli form: if non-sequestering sequencing is relatively slow or inefficient, the probability of obtaining multiple cDNA from a single RNA is low, and the mathematically simpler Bernoulli noise form approximately holds^16,133^.

Using the properties of PGFs^21^, we find that the overall generating function is given by a simple composition, substituting *G*_*ti*_ for *g*_*i*_:

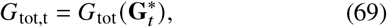

where we use the *G*_tot_ (**u**) parametrization, and each entry of 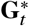 contains the shifted generating function 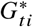 for a particular species *i*.

Finally, the reads associated with each cDNA 𝒯 are not always uniquely identifiable: for example, the sequence content is typically sufficient to identify the gene, but if a read only covers an exonic portion of the gene, it is impossible to distinguish whether or not the original molecule has been spliced^140^. To correctly represent this ambiguity, we need to transform the arguments of the generating function from a length-*n* vector to a length *𝓃*-vector, such that *𝓃* is the total number of mutually distinguishable classes of molecules. The simplest form of this transformation is a linear categorical partition:

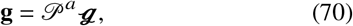

where 𝒫^*a*^ is an *n* × *𝓃* ambiguity matrix with 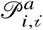 giving the probability of molecule *i* being identifiable in the equivalence class *i*. We assume that each molecule can be assigned to at least one class, implying 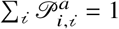. In principle, only the constraint 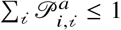 is mandatory, but the loss of molecules can be equivalently reframed as a technical noise component in 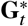.

We discuss the general case of this model component in Section S3. In summary, the entries of 𝒫^*a*^ are challenging to identify, but it may be possible to exploit genomic information, polymer physics, and orthogonal long-read sequencing data to construct it from first principles. This formulation admits several special cases. For example, if we cannot distinguish any distinct species at all and can only quantify the total RNA content, *𝓃* = 1 and 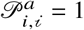 for each *i*. Then we obtain

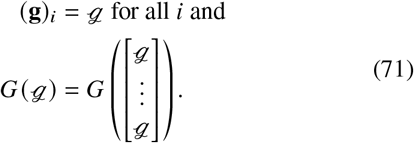

On the other hand, if all species are perfectly identifiable, we obtain *𝓃* = *n* and 𝒫^*a*^ = *I*_*n*_, the *n*-dimensional identity matrix. If, say, we have *n* = 2 but *𝓃* = 3, as in the case of nascent, mature, and ambiguous molecules described in La Manno et al.^86^ and Eldjárn Hjörleifsson et al.^140^, we obtain

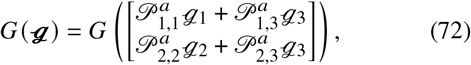

where *ℊ*_1_ and *ℊ*_2_ correspond to two unambiguously identifiable species, whereas *ℊ*_3_ corresponds to ambiguous cDNA which may have come from either. In the general case, we find

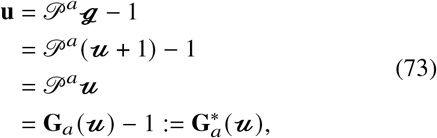

where each entry of the vector **G**_*a*_ contains the generating function of the relevant categorical distribution that governs how species *i* is parsed as one of the *𝓃* identifiable species:

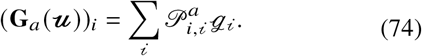

Therefore, the overall GF takes the following form:

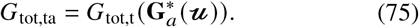

### 6.8 Example systems

The equation above provides a generic, modular framework for characterizing variability in sequencing experiments. To fit it to data, we need to specify a particular set of models for each step of the process. To do so, we should first strive to understand which modular components are realistic based on relatively simple summaries of data. Further, the process of evaluating and fitting these models is fairly involved, and often requires substantial up-front work to design scalable solvers. Therefore, it is useful to understand their qualitative behaviors relevant to statistical inquiry. In the current section, we characterize some analytically tractable systems, as well as their identifiability properties, such as our ability to distinguish between different models and parameter regimes. To illustrate these points, we apply the models to real and simulated data and speculate about their implications and physical relevance.

#### 6.8.1 Special theoretical cases

We revisit Section 6.3 to emphasize the implications and advantages of unifying the discrete and continuous degrees of freedom of the biological model in a common framework. The similarity of the discrete and continuous generating function terms is not accidental, and follows directly from the Poisson representation^93^. Occasionally, we can exploit this representation to bypass calculations for discrete processes by referring to results from the study of continuous processes, and vice versa. This approach consists of writing down the generating function PDE for a discrete process, identifying a continuous process governed by the same PDE, obtaining its solution from the stochastic process literature, and asserting that the discrete process distribution is given by compounding a Poisson distribution with the continuous law.

For instance, we may consider the case of a system with constitutive transcription at rate *α*, autocatalysis at rate *q*, and degradation at rate *γ*(*N* = 1, *n* = 1, *m* = 0):

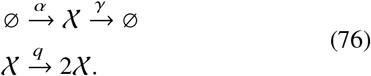

We can represent these reactions by the matrices *C* = − *γ + q* and *D* = *q*, as well as the operator *u* = *αu*. This system was introduced, but not treated, in Jahnke and Huisinga^85^, and, to our knowledge, first solved with master equation and generating function calculations by Vastola^17^. However, we can also solve it merely by matching terms, without any new calculations. We provide the full details of the parameter-matching process in Section S2.1. The derivation consists of noticing that the functional form of *C, D*, and 𝒜 can also arise from an *N* = 1, *n =* 0, *m* = 1 system with drift *α*, square-root noise 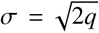, and mean-reversion at the rate *γ*−*q*. This is the Cox–Ingersoll–Ross (CIR) process, a popular mathematical finance model of interest rates^164,165^.

Its stationary distribution is gamma with shape *α*/*q* and scale 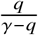. This immediately implies the distribution of the discrete process is negative binomial with the same shape and scale. This matches the result obtained by directly solving the master equation^18^. We find, then, that autocatalysis with constitutive transcription yields a stationary distribution equivalent to bursty transcription with no autocatalysis.

Obtaining this result, we may ask how the distribution changes if the molecules are produced in geometric bursts *B* with mean size *b*:

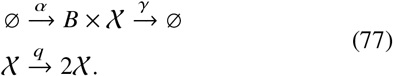

By changing the drift operator to a jump operator, we obtain a PDE with 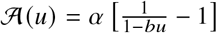. In other words, the continuous version of this process is a combination of CIR and gamma Ornstein–Uhlenbeck (Γ-OU) processes^20^, with the mean-reversion terms of both, the square-root noise of the former, and the exponentially-distributed jumps of the latter.

Define the parameter combinations

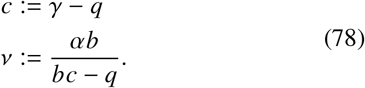

By direct integration, we find the characteristic and the stationary distribution

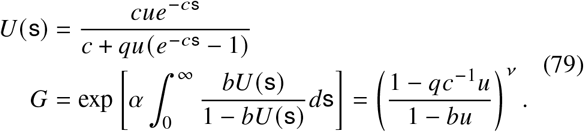

Curiously, this distribution exactly matches the *transient* MGF of the Γ-OU process, as well as the equivalent transient PGF of the bursty transcription process with no autocatalysis^16^:

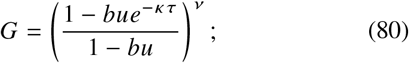

we may take advantage of the fact that *q*c^−1^ can be equivalently expressed as *be*^−*κ τ*^ < *b* for some positive *K* and *t*, because *bc* − *q* > 0 to have a steady state (i.e., positive *v*). In the continuous setting, this process is known^166^ to have a law consisting of a mixture of gamma distributions with scale *be*^−*κ τ*^ and shape *K*; in turn, *K* is drawn from a negative binomial distribution with shape *v* and scale (1+ *e*^−*κ τ*^)^−1^. This immediately implies that the distribution of the corresponding discrete process is a negative binomial-negative binomial mixture with equivalent parameters, which may be confirmed by the considerably more involved direct derivation in Section S2.2. Although this distribution cannot be expressed in closed form, its construction makes the simulation of the bursty transient and stationary autocatalytic processes trivial, and suggests that simple finite approximations (i.e., up to a modest *K*) may be developed.

The continuous formulation is a way to exploit existing quantitative results, but does not typically make problems easier. For example, we may be interested in solving an RNA/protein system with transcription, catalytic translation (at rate *q*), and the degradation of both species (at respective rates *γ*_*R*_ and *γ*_*P*_). Without specifying the transcriptional dynamics, we find that the downstream ODEs have a nontrivial *D* matrix, i.e.,

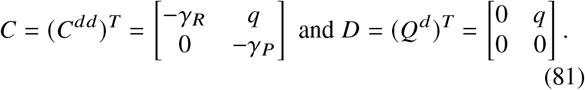

Although these matrices *can* be exploited to obtain both characteristics, the solution depends on special functions and is thus challenging to manipulate^77^. Instead, we may ask whether we can simplify the problem by eliding all stochasticity in the protein species and assuming it may be described by a continuous process. Defining the variables for this system, we find:

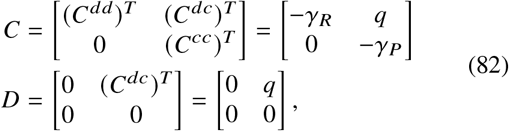

i.e., in spite of this supposed simplification, the problem is precisely as challenging as it was before. This provides an immediate and intuitive explanation for a range of results, such as the observation that the stationary distribution of proteins under constitutive transcription has a complicated solution in terms of Kummer’s hypergeometric function even if one uses a leading-order approximation (cf. Eqns. 34 and 50 of Bokes^139^).

#### 6.8.2 Empty droplets

##### Model definition

In Equation 64, we propose the simplest nontrivial model for the background distribution of RNA molecules in each droplet: the RNA content for each species *i* is described by a set of independent Poisson distributions whose mean is proportional to the mean in the entire cell population. Per Equation 65, the distribution of background is convolved with the endogenous RNA distribution of cell-containing droplets, making it challenging to distinguish technical and biological contributions. However, we *can* make predictions about the empty droplets, which have *G*_bc_ = 1, and compare these predictions to real datasets.

First, we define a baseline *n* = 2 model of biology, such that

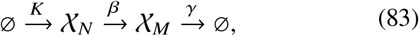

where *K* is a generic, but non-constant (bursty, multistate, or SDE-controlled) transcription process, *𝒳*_*N*_ is a *n*ascent transcript, *𝒳*_*M*_ is a *m*ature transcript, and *β* and *γ* are Markovian splicing and degradation rates, respectively. As the case of constant *K* yields a Poisson distribution of *𝒳*_*N*_ and*𝒳*_*M*_, the case of variable *K* induces an overdispersed distribution of RNA in droplets with one or more cells. Further, it implies that certain correlations are nonzero. For a given gene *j*, the correlation between counts of *𝒳*_*N*_ and *𝒳*_*M*_ should be nonzero, as the latter is, conceptually, the moving average of the former. Further, the correlation between the counts of a given species for different genes should be nonzero, as it reflects cell type heterogeneity and gene co-regulation^16^ (see Section 6.4).

This model describes the biology in living cells; to connect it to UMI measurements, we assume that 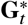 is an approximately linear map, i.e., library construction is either sequestering or non-sequestering and slow. Further, we assume 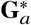 is a linear map, as in Equation 74. Therefore, for each species *i*, we have a per-cell biological distribution with mean *μ*_*i*_. In a droplet with a single cell, the mean becomes *μ*_*i*_ *p*_*i*_ (1+)≈ c *μ*_*i*_ *p*_*i*_, such that *p*_*i*_ is the overall probability of capturing, retaining, sequencing, and identifying each molecule (Section 6.7). In a droplet with no cells, the mean is c*μ*_*i*_ *p*_*i*_. We assume the number of doublets is negligible.

Under the foregoing assumptions we predict that the empty-droplet marginal per-gene UMI distribution is Poisson with mean c*μ*_*i*_ *p*_*i*_. This mean is proportional to the mean in non-empty droplets with a small coefficient of proportionality c. Further, we should observe zero correlations on an intra-gene basis, between counts of *𝒳*_*j,N*_ and *𝒳 𝒳*_*j,M*_, and on an inter-gene basis, e.g., between counts of *𝒳*_*j*1,*M*_ and *𝒳*_*j*2,*M*_. However, it is not *a priori* clear that this model should even approximately describe real data, even in the case of empty droplets. For example, these data may exhibit considerable “read depth” variability^65,83^, or, in our framework, inter-droplet variation in the probability *p*_*i*_, which would induce overdispersion or genome-wide correlations between molecule counts. By inspecting the distributional properties of empty droplet data, we can attempt to qualitatively motivate or raise doubts regarding the Poisson model.

##### Data processing

To build references and pseudoalign datasets, we used *kallisto* | *bustools* 0.26.0. We downloaded pre-built *H. sapiens* and *M. musculus* genomes from https://support.10xgenomics.com/single-cell-gene-expression/software/downloads/latest (10x Genomics, GRCh38 and mm10, 2020-A versions). Next, we used the kb ref function with the --lamanno option to build references. We obtained the raw FASTQ files for the six datasets reported in Table S2. Then, we used the kb count function with the --lamanno option, as well as the appropriate (10x v2 or v3) whitelist option -x to quantify the datasets, outputting unspliced and spliced RNA matrices. The unspliced counts correspond to molecular barcodes containing introns, whereas the spliced counts correspond to molecular barcodes not containing introns^140,167^. For the reasons outlined in Section S6 of Carilli and Gorin et al.^71^, we identify unspliced counts with “nascent” RNA species and spliced counts with “mature” RNA species, and elide any ambiguity.

##### Data analysis

We split the datasets into two categories. The “non-empty” droplets were retained after the *bustools* filter; the “empty” category contains barcodes that were discarded by the filter. Although this split is fairly coarse, as the filtering choices are heuristic, it is coherent with typical processing workflows and allows us to inspect the broad trends of distributional properties.

To investigate the overdispersion, or lack thereof, we separately computed the mean and variance of nascent and mature UMI counts for each gene in each set of cells. We plotted these quantities on a log-log scale, omitting the data points where one or both of these quantities were zero. Under the pseudobulk model, we expect the non-empty droplets to exhibit overdispersion and the empty droplets to be near identity, as the model encodes Poisson statistics for the latter. To investigate the intra-gene correlation structure, we computed the Pearson correlation coefficient *ρ* between nascent and mature UMI counts for each gene in each set of cells. We plotted the histograms of these values, as well as their relationship to the mature UMI mean, omitting the data points where *ρ* was undefined. To investigate the inter-gene correlation structure, we computed the Pearson correlation coefficient between the nascent UMI counts for each pair of genes in each set of cells, and repeated the analysis for mature count data. We plotted the histograms of these values, omitting the data points where *ρ* was undefined. As the number of gene pairs is fairly large, we first excluded all genes that were not expressed in the dataset. We expect both measures of correlation to be substantial for non-empty droplets and near zero for the empty droplets, as the model encodes statistical independence between marginal distributions for the latter.

To investigate the relationship between the empty and non-empty droplet averages, we plotted the mean mature UMI count for each gene in empty droplets against the mean mature UMI count in cell-containing droplets. As we plotted these quantities on a log-log scale, we omitted the data points where one or both of these quantities were zero. We repeated the analysis for mature RNA data. We expect these averages to be highly correlated, as the pseudobulk model proposes that the background RNA are sampled from a pool representative of the cell population.

Next, we computed and reported the Pearson correlation coefficient between the (well-defined) log-means. To characterize and explain deviations from Poisson behavior, we selected all genes with overdispersion in the mature RNA count distributions in empty droplets 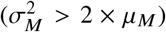 and reported their identities. Finally, to quantify the variation not included in the model, we computed the mean and variance of total mature UMI counts in empty droplets, with and without the overdispersed genes. As the sum of independent Poisson distributions is Poisson, we expect the total per-cell UMI count distributions to have a variance approximately equal to the mean.

#### 6.8.3 Noise-corrupted candidate models of transcriptional variation

##### Model definition

We would like to characterize the mutual distinguishability of superficially similar transcriptional models. In particular, we are interested in the benefits of multimodal data collection and the effects of technical noise.

As above, we begin by defining a baseline *n* = 2 model of biology, such that

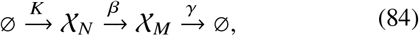

where *K* represents one of three candidate transcriptional models. The discrete dynamics are summarized by

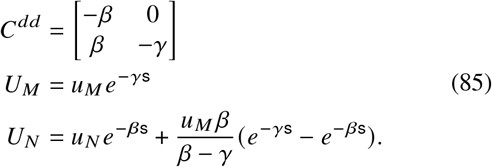

The first transcriptional model is the Γ-OU process, with *N* = 1 and *m* = 1:

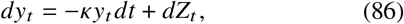

where *Z*_*t*_ is a subordinator with arrival rate *a* and exponentially distributed jumps with mean size *θ*. This system is characterized by

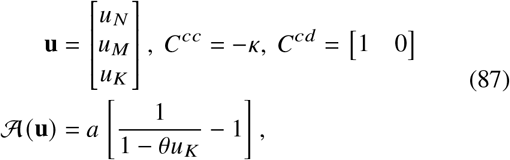

with all other matrices and operators set to zero.

The second is the CIR process, with *N* = 1 and *m* = 1:

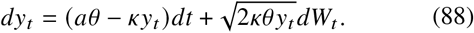

This system is characterized by

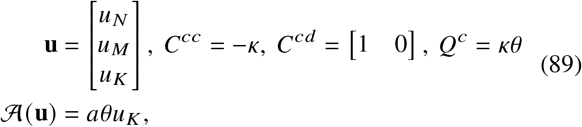

with all other matrices and operators set to zero.

We previously proposed the Γ-OU and CIR processes as potential explanatory models for gamma-distributed stochastic variability in transcription rates, solved them, and investigated the implications of their kinetics on the model properties and distinguishability^20^. The stationary distribution of the Γ-OU and CIR processes is gamma, with shape *a*/*κ* and scale *θ*, i.e., mean *aθ/k and* variance *aθ*^2^/*κ*. In addition, their (appropriately normalized) autocorrelation function is *e*^−*kt*^.

Finally, th0065 third is the telegraph process^100^, with *N* = 2 and *m* = 0. This system is characterized by

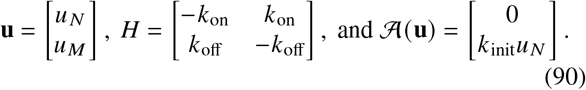

The stationary distribution of this process is Bernoulli scaled by *K*_*init*_, with mean 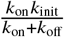 and variance 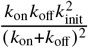. Its auto-correlation function is 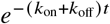^81^.

For all three models, assuming a Bernoulli observation model (i.e., that each molecule has an independent probability *p* of being observed) is equivalent to a parameter redefinition. For the Γ-OU and CIR models, this redefinition is that *θ* → *pθ*; for the telegraph model, we have analogously that *K*_init_→ *pk*_init_.

Let us see why this is true. Recall from Section 3 that the Bernoulli technical noise model amounts to a redefinition *u*_*N*_ *→ pu*_*N*_, *u*_*M*_ → *pu*_*M*_. *For* the Γ-OU model, the steady-state (log-) GF is

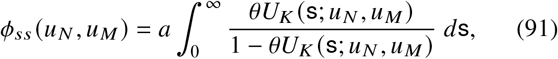

where *U*_*K*_ (s; *u*_*N*_, *u*_*M*_) is the exponential sum solution of

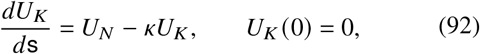

and where the characteristics *U*_*N*_ and *U*_*M*_ are as in Equation 85. Because the *U*_*K*_ ODE is linear, *U*_*K*_ depends linearly on *u*_*N*_ and *u*_*M*_ (and hence on *p*). But *ϕ*_*ss*_ only depends on *U*_*K*_ through the combination *θU*_*K*_, so the problem with technical noise is equivalent to redefining *θ* → *pθ*.

For the CIR model, the steady-state (log-) GF is

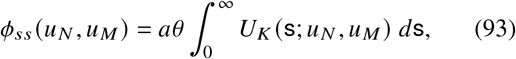

where

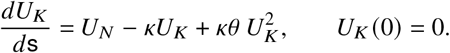

The technical noise causes *U*_*N*_ → *pU*_*N*_. Divide both sides by *p*, so that the *p* factor is moved elsewhere; we can see that

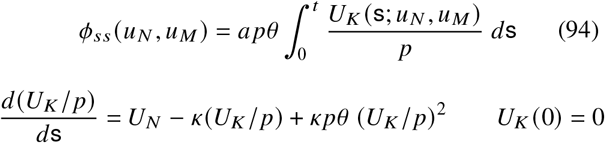

is equivalent, i.e., that again the technical noise problem is equivalent to a non-technical-noise problem with *θ*→ *pθ*.

For the telegraph model, the steady-state (log-) GF is

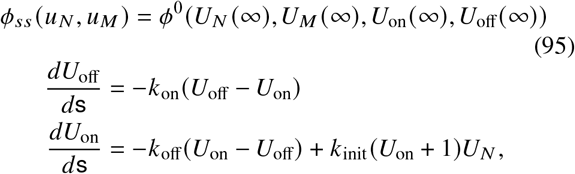

where *U*_off_ 0 = *U*_on_ 0 = 0. Since *U*_*N*_ (∞)= *U*_*M*_(∞) = 0, the values of *U*_*N*_ s only affect *ϕ*_*ss*_ through the combination *K*_init_*U*_*N*_ that appears in the *U*_on_ ODE; this means we can just redefine *K*_init_ → *pk*_init_ as promised to get a completely equivalent problem.

##### Model analysis

Formally, these models have five parameters each: three for the upstream transcriptional dynamics and two for the downstream molecular conversion. However, their qualitative behaviors at steady state can be effectively summarized by fixing *μ*_*K*_, *β*, and *γ*, and varying two key parameters, the timescale separation and the noise intensity. From a statistical point of view, *μ*_*K*_ *β* and *μ*_*K*_ *γ* are easily and robustly identifiable from the mean molecular counts; from an experimental point of view, *β* and *γ* can, in principle, be fit by orthogonal experiments^86^. At steady state, the value of *μ*_*K*_ is a somewhat arbitrary scaling factor.

For the two-species SDE driver models, the qualitative parameters take the following form:

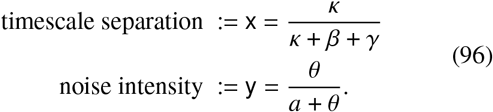

These parameters both range in 0, 1. When the timescale separation approaches zero, the transcriptional variation is much slower than the turnover, and the distribution of RNA is given by a simple Poisson mixture of the law of *K*. When the noise intensity approaches zero, the law of *K* degenerates and the distribution of RNA becomes Poisson. Most interestingly, when the timescale separation and the noise intensity are both high, the system exhibits bursty transcription^20^.

Equation 96 is defined with reference to the process parameters of the Γ-OU and CIR drivers^20^. It remains to define *κ, θ*, and *a* in terms of *K*_on_, *K*_off_, and *K*_init_ for the telegraph process. The correct identification is:

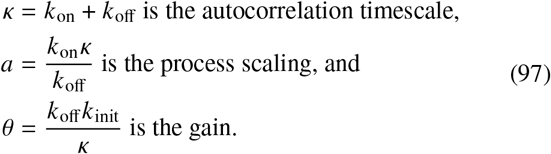

These identifications are not arbitrary, as they endow the system with lower moments that match the SDE formulation: autocorrelation function *e*^−*κt*^, mean *aθ/κ, and* variance 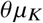. In addition, the system has the correct geometric burst limit (*K*_init_, *K*_off_ →∞) with burst size *θ*/ *k k*_init_/*K*_off_ and burst frequency *a* → *K*_on_^73^; this limit matches the Γ-OU one^20^.

Given any combination of {x, y, *μ*_*K*_, *β, γ*}, we can identify the transcriptional parameters:

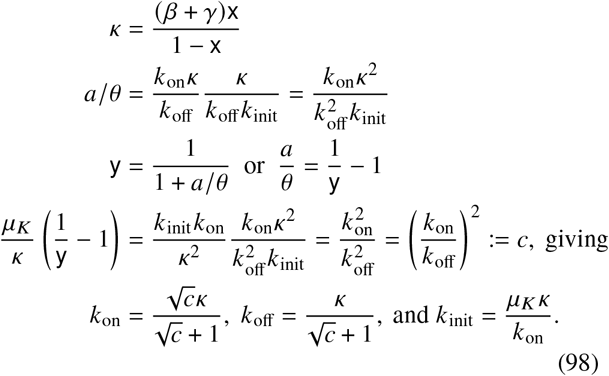

This allows us to define a particular set of {*μ*_*K*_, *β, γ*}, vary x and y over the constrained domain (0, 1) × (0, 1), and compare the model properties for each (x, y) tuple. If we are interested in a one-species model, we simply replace each instance of *β* +*γ* with *β*. Since the construction in Equation 98 is bijective, if we fairly densely sample the square, we can be confident that the results fully encompass the range of behaviors under a particular set of averages.

##### Simulated data analysis

To evaluate PMFs, we used trapezoidal quadrature for the Γ-OU generating function integral, the Runge-Kutta method for the CIR characteristic *U*_*K*_ and trapezoidal quadrature for the generating function integral, and the Runge-Kutta method for the telegraph model’s coupled differential equations^18,20^. We marginalized over the continuous and categorical dimensions. We evaluated all PMFs on *x*_*N*_, *x*_*M*_ ∈ [0, ⋯ 49] × [0, ⋯ 50]. To generate synthetic data, we sampled with replacement from the 2,550 microstates in the domain, using *P*(*x*_*N*_, *x*_*M*_) as sampling probabilities.

To investigate parameter identifiability, we generated 200 realizations from the Γ-OU model under *κ* = 0.1, *a* = 0.4, *θ* = 1, *β* = 0.8, and *γ* = 0.9. These parameters lie in the “mixture-like” regime, where the transcriptional process is slower than the RNA turnover process. Next, we constructed a uniformly spaced 14 ×15 grid of x and y, constructed at the true values of *μ*_*K*_, *β*, and *γ* and bounded by[0.01, 0.99]. In statistical terms, this model formulation is the best-case scenario where no noise exists and uncertainty in the fixed parameters is negligible.

To investigate the statistical properties of one-species data, we evaluated the log-likelihood log *L* of the nascent marginal of the data at each of the 210 x, y coordinates (with the true value being x = 1/ 9 and y = 5/7). Next, we plotted log *L* as a heatmap over x, y. The coordinates with high log *L* are not readily distinguishable, i.e., these parameters produce very similar distributions to the data. We highlighted the coordinates in the 90th percentile of log *L*—the least distinguishable region—using hatching. To illustrate a case where the one-species data are relatively uninformative, we considered a point with x = 9/10 and y = 5/7, which lies in the qualitatively different “burst-like” regime (*κ* = 7.2) but closely resembles the “mixture-like” data at steady state.

To investigate the statistical properties of two-species data, we repeated the analysis above, computing the joint likelihood rather than the marginal likelihood. In the two-species model, the true “mixture-like” parameter set has x = 1/ 18 and the illustrative “burst-like” parameter set has x ≈ 0.81; the other parameters do not change. To demonstrate the source of failure to distinguish between these parameter regimes, we plotted the PMFs in both. We used a transparent bar plot for the nascent PMFs and a heatmap for the joint PMFs, with darker colors representing a higher probability mass.

To investigate the mutual identifiability of models, we computed their Akaike weights over the x, y landscape. The Akaike weight of model 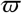 is defined as follows:

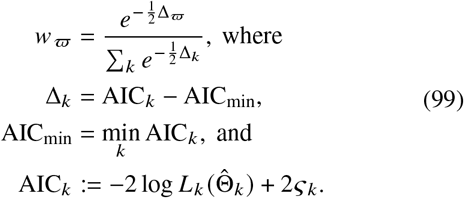

Thus, AIC_*K*_ is the Akaike information criterion (AIC) for model *K*. The AIC depends on the model log likelihood log *L*_*K*_ at the maximum likelihood estimate 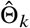, as well as number of model parameters *ς*_*K*_^120^. Therefore, the Akaike weight essentially transforms and combines the models’ relative likelihoods to provide a measure of their agreement with the data.

Although this measure has its caveats and limitations—for example, it cannot account for uncertainty in the modelspecific parameters Θ_*K*_—it is a fairly conventional criterion for model selection. Most usefully to our investigation, it admits a simple interpretation: if the Akaike weight of the true model *w*_ϖ_ ≈ 1/3, there is essentially no basis for choosing a particular model, since their distributions are not practically distinguishable. If *w*_ϖ_ >1/2, we have a basis for model discrimination: the odds for the correct model are even. In the three-model case, this may reflect both, or only one, of the competing hypotheses being substantially worse at describing the data, so more careful examination of the *w*_*K*_ values is necessary to judge the models.

To investigate model identifiability, we constructed a uniformly spaced 14 ×15 grid of x and y, bounded by 0.01, 0.99. At each grid point, we generated 200 realizations from the Γ-OU model under *μ*_*K*_ = 5, *β* = 0.8, and *γ* = 0.9. Next, we computed the log *L*_*K*_ of each model using the nascent marginal and the full data, and used the relative likelihoods to compute the Akaike weights of the Γ-OU model under these two scenarios. Finally, to reduce the impact of stochastic sampling variability, we repeated the process 50 times and computed their average. In other words, we generated 50 independent datasets at each of the 210 grid points, evaluated likelihoods of all models, computed the univariate and bivariate Γ-OU Akaike weight of each, then aggregated the 50 trials at each grid point to obtain two “average-case” performance measures. In statistical terms, this model formulation represents the best-case scenario where the parameters are perfectly known, and the problem solely consists of distinguishing between the models, as in the Γ-OU/CIR case considered in Fig. 3 of Gorin and Vastola et al.^20^

To visualize the behavior of the Akaike weights under these assumptions, we plotted its value as a heatmap over x, y. We highlighted the coordinates with *w*_*ϖ*_ < 1 2—the poorly distinguishable region—using hatching. To illustrate a case where the one-species data are relatively uninformative, we compared a point with one-species coordinates x, y = (0.4, 0.9), which lies in the “mixture-like” regime, to one with x, y = (0.9, 0.8), which lies in the “burst-like” regime. We visualized these points on the x, y axes using large, color-coded circles. From Gorin and Vastola et al.^20^ and the properties of low-x processes outlined in the definition of x, we expect the former regime to be highly distinguishable, particularly since the telegraph process converges to a bimodal Bernoulli mixture for *κ* → 0. On the other hand, we expect the latter regime to be somewhat less distinguishable; in this limit, the Γ-OU and telegraph models both converge to the bursty model discussed in Singh and Bokes^138^. We repeated this analysis for two-species Akaike weights, transforming the coordinates appropriately (i.e., x ≈ 0.24 for the mixture-like regime and x≈ 0.81 for the burst-like regime).

To demonstrate the basis of statistical distinguishability properties, we plotted the PMFs of the three models in the two parameter regimes. To simultaneously display them, we plotted marginal distributions of the nascent species as line charts, color-coded by the model identity.

To investigate the effect of drop-out technical noise, we did not perform dedicated simulations; instead, we exploited the result, derived above, that the functional form of the solutions is closed under downsampling. In other words, all distributional properties of a system with gain *θ* and the technical noise parameter *p* are identical to those of a system with gain *pθ* and no technical noise. These properties include the model distinguishability. To illustrate this result, we represented Bernoulli technical noise by arrows in the negative y direction, with small circles located on an arrow corresponding to 50%, 75%, and 85% dropout. To compute the y value under dropout, we use:

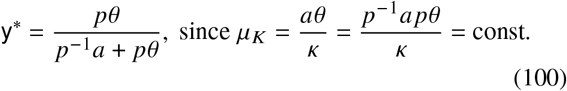

The arrows begin at 0% dropout, corresponding to the illustrative base cases (large circles) described above. This demonstrates that increasing the drop-out rate while holding the averages constant leads to the molecular distributions’ degeneration to the Poisson limit. If we do not hold the averages constant, we simply obtain the decreased 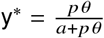on the (less identifiable) x, y. landscape with mean transcription rate *pμ*_*K*∞_

#### 6.8.4 Distributions obtained from a transient process

##### Model definition

As motivated in our RNA velocity review^19^, understanding transient developmental processes that occur on a timescale comparable to RNA lifetimes requires fitting transient probabilistic models. Even under the considerable simplifications made in Section 6.5, fully treating transient transcriptional phenomena requires identifying the *a priori* unknown (1) internal-age distribution *ft* as well as (2) process parameters for *G*(*t*). As the time since process start *t* can be conceptualized as a cell-specific latent variable, this problem can be treated by an expectation–maximization (EM) algorithm, which may proceed by probabilistically constraining the unknown (3) *c*ell-specific times *t*_c_.

Since parameter inference is mandatory for the expectation step of the EM algorithm, we begin by characterizing the upper limit on its performance. In particular, previous attempts to treat the problem have assumed simple Gaussian or Poisson error terms^59,86,121^, or applied graph methods^168^. These approaches do not recapitulate^19^ the discrete stochasticity and bursting observed in transient biophysical processes^125,169^. However, the transient distributions of bursty processes are not available in closed form, and require new algorithms. Therefore, we treat the simplest nontrivial formulation, which combines points (1) and (2), while omitting (3): if we have perfect information about the cells’ relative times, can we satisfactorily fit a bursty transcriptional model and use the results as a basis for distinguishing between internal-age distributions?

We define a baseline *N* = 1, *n* = 2, *m* = 0 model of biology with no technical noise, with the reaction schema

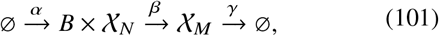

representing bursty transcription with stochastic burst sizes B drawn from a geometric distribution with time-dependent mean *b*(*t*):

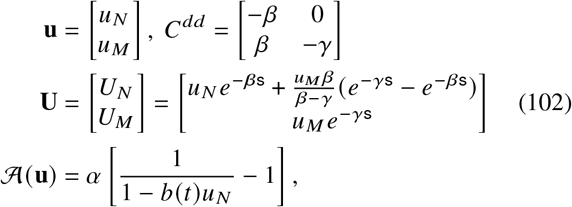

with all other operators set to zero. To specify *b*(*t*), we define a three-stage model of cell type transitions, such that

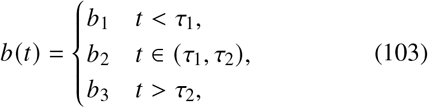

i.e., a transition is accompanied by a rapid change in burst size at a deterministic time after starting the process.

Next, we propose candidate internal-age distributions. Drawing on the chemical engineering literature^105,106^, we outline one-parameter reactor models, such that *t* = 0 corresponds to the cell entering the reactor; after some residence time *t*, which is dependent on reactor architecture and drawn from the distribution *f*_res_, the cell exits. The internal-age distribution is given by

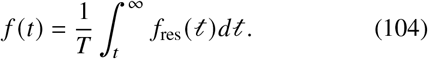

The plug flow reactor (PFR) is the model implicit in previous studies^59,86^. Formally, it represents each cell entering a reactor, then exiting after some deterministic time *T*. Its residence-time distribution is Dirac or degenerate, with *f*_res_(*t*) = *δ*(*t* − *T*), so

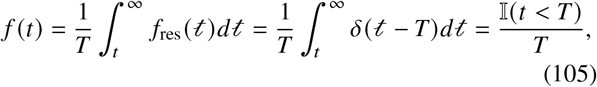

the expected uniform distribution. This distribution has the CDF and inverse CDF

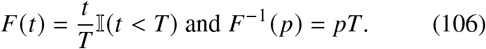

The continuously stirred tank reactor (CSTR) represents a cell entering a homogeneous reactor, then exiting after a random time, in a memoryless fashion. Therefore, the residence-time distribution 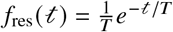 is memoryless or exponential, yielding

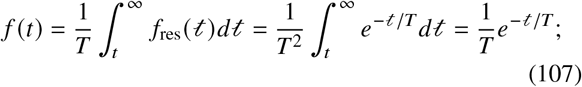

i.e., memorylessness implies that the properties inside the reactor—including the age distribution—are identical to the properties of the efflux stream. We obtain the CDF and inverse CDF

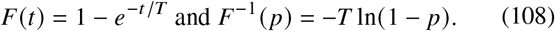

The laminar-flow reactor (LFR) is a configuration between these two extremes: it represents a cell entering a reactor, remaining in it for some time deterministic time, then being able to exit after a power-law delay. Its residence-time distribution 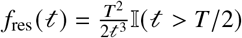 is Pareto, yielding

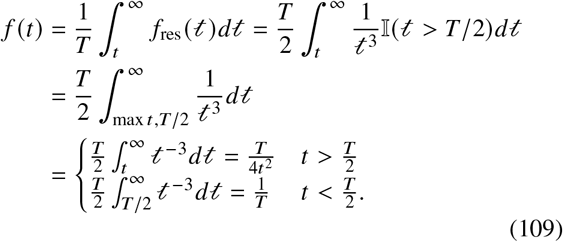

The PDF can be integrated to yield the CDF and inverse CDF

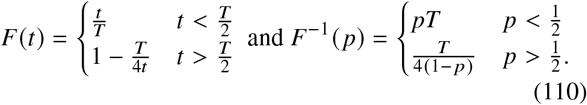

We are interested in the CDFs and inverse CDFs of the internal-age distributions because “perfect information about the cells’ relative times” properly requires specifying {*F*_*ϖ*_(*t*_*c*_)} and {*F*_*ϖ*_(*τ*_*i*_)} under the true model *ϖ* rather than the raw {*t*_*c*_} and {*τ*_*i*_)} values. Otherwise, the model selection problem becomes somewhat trivial; for example, if we know the mean residence time is *T* and we know one of *t*_c_ > *T*, we can immediately eliminate the PFR configuration without performing any calculations.

A synthetic dataset consists of observations *x*_*N*,c_, *x*_*M*,c_ for each cell c, generated from the true model *ϖ* at the true time point *t*_c_. The log-likelihood of parameters Θ_*K*_ = {*b*_1_, *b*_2_, *b*_3_, *α, β, γ*}_*K*_ for model *K* takes the form

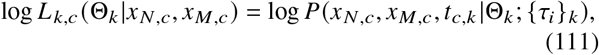

where 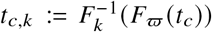 and 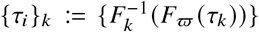 are the transformed times. This yields the full log-likelihood under the assumption of independence

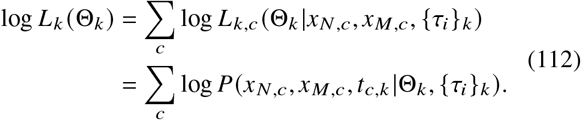

The problem of identifying the maximum likelihood parameter set consists of optimizing Equation 112 with respect to Θ_*K*_. The problem of reactor identification consists of using the resulting reactor-specific maximum likelihood value log 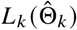 with Equation 99 to obtain the Akaike weights of each reactor configuration.

##### Simulated data analysis

To generate the illustrations in Figure 4a, we directly simulated cells entering and exiting each reactor configuration. First, we sampled arrival times from a uniform distribution on 0, 100. Next, we sampled residence times by inverse transform sampling from the inverse CDF corresponding to each f_res_, using the mean residence time *T* = 2. We arbitrarily selected the observation time 75 and selected all cells which had arrived but not exited at this time. We computed the cell age by subtracting the arrival time from the current time. We repeated this procedure 10^7^ times for each reactor to obtain the internal-age distribution. Next, we computed the histogram of the distribution on 0, 10, using 200 bins. To account for the fact that this histogram only contains part of the CSTR and LFR densities, we rescaled the bins by the internal-age distribution’s CDF value at *t* = 10. Finally, we plotted the rescaled histogram as a bar plot, and the analytical *f*as a line plot for comparison.

To understand the actionable differences between reactors, we simulated data from a single reactor model, then fit all three models to the obtained counts. First, we sampled 200 true reaction times *t*_c_ under the PFR model with *T* = 5 and sorted them. To generate synthetic data, we used Gillespie’s stochastic simulation algorithm^142,155^ with a time-dependent burst size, storing the state of the system at {*t*_c_}. We generated 200 realizations, using only one realization per time point to fit the models. To simulate, we used the parameters Θ_*ϖ*_ = {*b*_1_, *b*_2_, *b*_3_, *α, β, γ*}_*ϖ*_ = {2, 5, 1, 0.8, 1.2, 3.14}. We set {*τ*_1_, *τ*_2_} to {1, 3}. We started the system in a bivariate Poisson initial distribution with 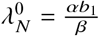 nascent and 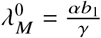 mature molecules on average. Although this initial condition is somewhat arbitrary, as it is out of equilibrium, it is readily tractable and yields a constant mean over the first stage of the process.

The instantaneous probability *P*(*x*_*N*,c_, *x*_*M*,c_, *t*_c,*K*_ |Θ_*K*_, {*τ*_*i*_} _*K*_) is not available in closed form, and needs to be obtained by inverting the generating function for each *t*_c,*K*_^16,20,138^:

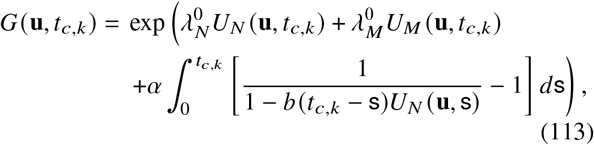

where we elide the dependence of *b* on the model-specific {*τ*_*i*_}_*K*_. For a given value of **u**, it is straightforward to propagate the initial condition. However, it is impractical to compute the integral separately for each c. We can bypass this bottleneck by reusing quadrature points. Conceptually, we define the quadrature matrices

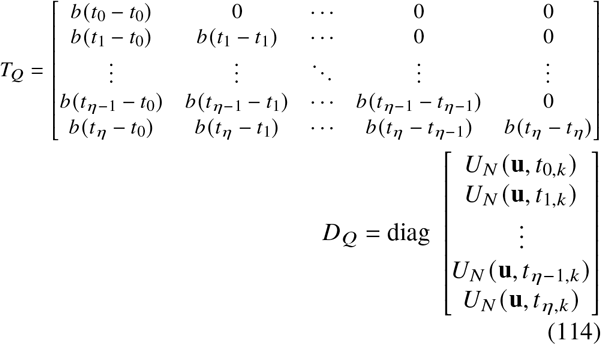

in the general case with *η* cells. We appended the starting grid point *t*_0,*K*_ := 0 to properly integrate from zero. We use the notation *T*_]_ because this matrix is Toeplitz in the narrow, but numerically relevant^19^, case of a uniformly spaced grid approximating sampling from a PFR. To lighten the notation, we drop the subscript *K* from the time points in the definition of *T*_*Q*_. *D*_*Q*_ is diagonal and does not need to be constructed explicitly; to obtain the product *T*_*Q*_ *D*_*Q*_, we broadcast *T*_*Q*_ with the vector used in the definition of *D*_*Q*_. Then, we computed *M*_*Q*_ = (1 −*T*_*Q*_ *D*_*Q*_) ^⊙(−1)^ 1, where ⊙(−1) is to be interpreted as the elementwise/Hadamard inverse of the matrix. Finally, we approximated the integral by applying the *NumPy* quadrature algorithm trapz along the rows of *M*_*Q*_, using {*t*_c,*K*_} as the integration grid^170^. The GF evaluation grid size was set to [0, ⋯, max *x*_*N*_ +4] × [0, ⋯, max *x*_*M*_ +4], where max *x*_*i*_ is the highest RNA count observed for species *i* over the entire simulation, in all cells.

Next, we used the *SciPy* algorithm optimize.minimize^171^ to minimize the negative log-likelihood of the data under all three models, and obtain a satisfactory set of parameters. Specifically, we varied the 6-dimensional vector log_10_ Θ, with each log-parameter’s bounds set to (1.5, −1.5). We optimized with the L-BFGS-B solver for a maximum of 20 steps. Since we are primarily interested in the models’ relative performance at their maximum likelihood estimates (MLEs), rather than the process of obtaining these estimates, we initialized each search at the parameters used to generate the data.

Next, we sought to illustrate the fit performance and the differences between the models’ distributions. We plotted the marginals of the simulated data at each time point *t*_c_ as bar plots, now using the counts from all 200 cells to demonstrate the full transient distribution. Next, we plotted the marginal PMFs of the three models at the corresponding time points *t*_c,*K*_ as color-coded line charts. We expect the true reactor configuration (PFR) to closely agree with the distribution shapes; however, we have no *a priori* information regarding how well other reactor architectures can recapitulate the same data. To quantify the prospects for model selection, we inserted the optimal log-likelihoods into Equation 99 and calculated the Akaike weights of the model candidates.

To characterize the identifiability properties, we reproduced the simulation and analysis process using the same parameters, but varying the dataset size, with *η* = {20, 40, 60, 80, 100, 150, 200}. For each *η*, we generated 50 synthetic datasets, fit them, and computed the Akaike weights of the models. We plotted all *w*_*ϖ*_ as a function of the number of cells, adding uniform jitter to facilitate inspection. To visualize the trends in model identifiability, we plotted the mean and standard deviations of all *w*_*ϖ*_ for a given *η*, connecting them with a line to guide the eye. We do not *a priori* know whether the reactor configurations are meaningfully distinguishable, but if they are, we expect them to become more so with more data.

Next, we sought to characterize the prospects for distinguishing reactor models for a broader range of transcriptional parameters. We used rejection sampling to draw Θ_*ϖ*_. First, we drew log_10_ *b*_*i*_ from a normal distribution with mean 0.8 and standard deviation 1, and all other log-parameters from a normal distribution with mean 0 and standard deviation 1. The parameters were clipped to stay in the domain [10^−1.4^, 10^1.4^] to avoid “trivial” regimes with excessive timescale separation relative to the reactor residence time. Next, we found the highest *b*_*i*_, computed the nascent and mature mean and standard deviation corresponding to this set of *b*_*i*_, *α, β, γ*^138^, and kept the proposed Θ_*ϖ*_ if *μ*_*N*_+4]_*N*_ and *μ*_*M*_ + 4]_*M*_ were both lower than 25. Otherwise, we regenerated Θ_*ϖ*_. This is an *ad hoc* way to limit the state space size for PMF evaluation: although we do not know what the maximum observed counts will be until we simulate the system, *μ* 4] is typically provides a reasonable estimate^97^. Rejecting parameters in this fashion approximately limited the state space size to 25 × 25. In this way, we simulated, fit, and computed the Akaike weights for 200 parameter sets. All used the PFR ground truth model, {*τ*_1_, *τ*_2_} = {1, 3}, and *T* = 5 as above.

To summarize the model identifiability over this domain of synthetic parameters, we plotted the distribution of AIC weights *w*_*ϖ*_. Finally, to characterize the relationships between the models, we plotted the distributions of log-likelihood differences log 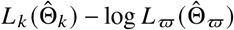, where *K* corresponds to the CSTR and LFR models, as transparent histograms colorcoded by *K*. If such a histogram is skewed toward negative values, the model *K* produces consistently worse fits than the true PFR model. In the other hand, if it is centered at zero, then model *K* is typically easily confused with the true model. We restricted this visualization to (−5, 5) to compensate for potential failure to converge, which produces inflated likelihood differences. This visualization provides a basis for explaining the distribution of *w*_*ϖ*_.

#### 6.8.5. Variability in library construction

##### Model definition

In section 6.8.3, we considered the parameter and model identifiability for a two-stage model of RNA processing, and found that several interesting distributions are closed under downsampling, so long as the downsampling is Bernoulli with equal parameters for both species. However, this assumption may be too restrictive in practice: for example, nascent RNA may be more or less likely to be captured than mature RNA, depending on the poly(A) content of their introns. In the current section, we investigate the behavior of models with differences in capture probabilities or rates.

The identifiability properties are highly model-dependent. For example, if we consider the Γ-OU or CIR models, with *N* = 1, *n* = 2, *m* = 1, such that

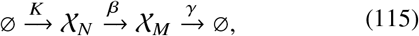

where the autocorrelation of *K* is *κ* ≪ *β, γ*, the stationary distribution of *K* is gamma with shape *ν* = *a*/*κ* and scale *θ*. We find the stationary RNA generating function is bivariate negative binomial, with

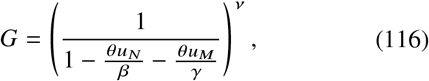

which is outlined in the supplemental section 2.5.2 of Gorin and Vastola et al.^20^ Under sampling, the distribution stays bivariate negative binomial, with GF

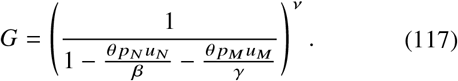

In other words, even if we have perfect information about this distribution’s three parameters *ν, θ p*_*N*_/*β*, and *θ p*_*M*_/*γ*, we cannot conclude anything about the magnitudes of *p*_*N*_ and *p*_*M*_, as they are degenerate with *θ, β*, and *γ*. If *K* is telegraph (i.e., *N* = 2, *n* = 2, *m* = 0), we obtain a finite Poisson mixture:

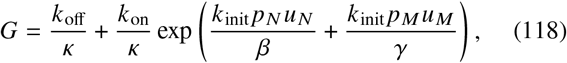

which exhibits the same degeneracy with respect to *K*_init_, *β*, and *γ*. Entirely analogously, if the system is in the Poisson limit (y 0) with average transcriptional strength *μ*_*K*_, we find that sampling yields

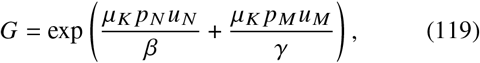

which is non-identifiable.

Interestingly, the bursty regime *is* partially identifiable. We begin by defining a baseline *N* = 1, *n* = 2, *m* = 0 model of biology with technical noise but no ambiguity, such that

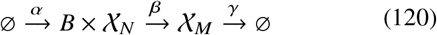

representing bursty transcription with stochastic burst sizes *B* drawn from a geometric distribution with constant mean *b*. Further, we assume that a molecule X_*i*_ is retained with probability *p*_*i*_, yielding:

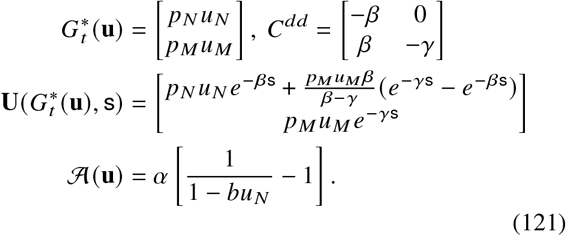

In other words, the stationary generating function is given by

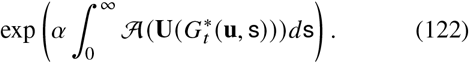

In principle, this quantity can be integrated, inverted, and optimized with respect to the parameters. However, to be thorough, we need to reformulate the optimization problem in the most compact form available, which involves identifying the distribution’s degeneracies. Although this system formally has six parameters *b, α, β, γ, p*_*N*_, *p*_*M*_, at steady state only four are identifiable. This is made clear by examining the integrand:

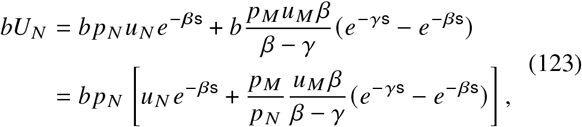

i.e., the characteristic is invariant so long as *b p*_*N*_ and *p*_*M*_/*p*_*N*_ are constant. By plugging in zero for *u*_*N*_ or *u*_*M*_, we observe that the characteristics take the functional form of the characteristics of the noise-free system, implying different values of *p*_*N*_ and *p*_*M*_ may give indistinguishable distributions. Therefore, identifying the relationship between *p*_*N*_ and *p*_*M*_ requires bivariate data. To quantitatively characterize *how* identifiable *p*_*N*_ and *p*_*M*_ are, we need to use simulations.

However, challenges particular to single-cell technologies arise when attempting to apply this model to large datasets with many genes. Although the Bernoulli model is a useful approximation, considering the sequencing process suggests that the non-sequestering technical noise model is more realistic: there is no chemical barrier to an RNA molecule being captured multiple times. In this formulation, each gene’s technical noise is parametrized by the species’ overall capture rates *λ*_*N*_ and *λ*_*M*_, which produce the Bernoulli limit when both of these parameters are small.

Furthermore, it appears implausible that *λ*_*j,N*_ and *λ*_*j,M*_, where *j* indexes over genes, vary arbitrarily. In a previous report^21^, we have found that the model *λ*_*j,N*_ = *C*_*N*_ *L*_*j*_ and *λ*_*j,M*_ = *λ*_*M*_ performs satisfactorily. In this model, the nascent species are identified with unspliced molecules, which are considerably longer than spliced molecules and contain a large number of internal poly(A) priming sites. To a first-order approximation, we may propose that nascent species are captured at a rate proportional to the gene length *L*_*j*_, where the constant of proportionality *C*_*N*_ is a dataset-wide technical noise parameter. Analogously, we identify the mature species with fully spliced, poly(A)-tailed molecules, and make the zeroth-order approximation that poly(A) tails are chemically identical. The capture rate *λ*_*M*_ is, then, also dataset-wide. Although this model is relatively simplistic, it foregrounds a key challenge. Even if we assume different genes’ transcriptional processes are independent, we cannot fit their distributions independently, as we need to account for coupling through the technical noise parameters.

##### Data analysis

To illustrate the identifiability of *p*_*M*_ *p*_*N*_ under the Bernoulli noise model, we considered the likelihood landscape for the simplest one-parameter formulation. We fixed the parameters *α* = 1, *b p*_*N*_ = 4.9, 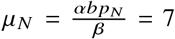, and 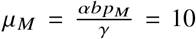; in other words, the nascent RNA distribution is negative binomial with shape 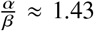 and scale *b p*_*N*_. We simulated data at *p*_*M*_/*p*_*N*_ ∈1 4, 1, 4, with *η* = {20, 50, 100, 200} simulated cells. For each of the true *p*_*M*_/*p*_*N*_ and *η* values, we generated 200 datasets by sampling from the PMF on [0, ⋯, 99] × [0, ⋯, 99]. To evaluate the PMF for *p*_*M*_ > *p*_*N*_, we set *p*_*M*_ to unity with no loss of generality. To evaluate it for *p*_*N*_ > *p*_*M*_, we set *p*_*N*_ to unity.

This yields 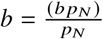 *a*nd 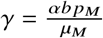. Next, we computed the likelihood of the data under log_10_ *p*_*M*_/*p*_*N*_ [2, −2], keeping *α, b p*_*N*_, *μ*_*N*_, and *μ*_*M*_ constant, using the evaluation grid size[0, ⋯, max *x*_*N*_ +3] × [0, ⋯, max *x*_*M*_ +3], where max *x*_*i*_ is the maximum observed for each species in the simulation. We used 200 log_10_ *p*_*M*/_ *p*_*N*_ grid points, evenly spaced throughout the domain. Next, we computed the posteriors over the grid by dividing each likelihood vector by its sum. Finally, we plotted the average posterior distribution using line charts, with the color indicating the true value of *p*_*M*_/*p*_*N*_ and the intensity indicating the number of cells, with more saturated colors corresponding to more simulated cells. For ease of comparison, we plotted the true values using dashed lines. From a statistical perspective, this analysis summarizes the parameter identifiability conditional on perfect information about the nascent marginal and the species averages. As we do not *a priori* know whether the differences in the PGF are actionable, the analysis illustrates the sample sizes required to fit the parameter to a particular degree of precision.

We previously motivated and fit the Poisson model of technical noise^21,133^. In Gorin et al.^21^, we inspected a variety of datasets, and observed a pronounced length bias in the nascent RNA count data, which did not appear in mature RNA counts (Section S7.3 of Gorin et al.^21^). This bias may be explained by three naïve models of biology.

The first model posits that the nascent RNA molecules are in the process of being transcribed; higher amounts of nascent RNA for longer genes simply reflect longer elongation delays. Although this explanation is superficially plausible, it is not borne out by the data. The model predicts a geometric-Poisson distribution of nascent RNA and zero correlation between nascent and mature counts^155,172^. Real data, on the other hand, have distinctly negative binomial-like marginals (as evident in, e.g., the third column of Fig. 4b of our recent work on delay CMEs^155^, which shows consistently inferior fits under the delay model), and nontrivial nascent/mature correlations (as in the red histogram in Figure 2b).

The second model posits that the differences in expression reflect real differences in the underlying biological parameters, and technical noise may be neglected. However, fitting this model produces pervasive length biases in the parameter values (Section S7.4 of Gorin et al.^21^), which are inconsistent with trends observed in orthogonal data. This is the model we explored in Gorin et al.^21^

The third model posits that technical noise *does* occur, but takes the species-independent form *p*_*N*_ = *p*_*M*_. This formulation is mathematically identical to the second model, but proposes that an *apparent* length bias in the burst size is *actually* a length bias in *b p*. This model partially bypasses the objection raised for the second model by proposing that *p* is gene length-dependent, identical for nascent and mature species, and higher for longer genes. However, this model is implausible on physical grounds, as mature transcripts do not have the intronic poly(A) content necessary to produce this length dependence. This is indirectly implied by the consistently low fraction of exonic reads in sequencing datasets, in contrast to introns and the 3’ untranslated region^137^.

These biases can be largely eliminated by proposing a length-dependent sampling rate for nascent RNA counts, suggesting that this technical noise model is more coherent with known biology. To illustrate this process, we summarize the key results from Gorin et al.^21^

We obtained the raw data for the twelve 10x v3 datasets reported in Table S4 of Gorin et al.^21^. The raw data consisted of nascent and mature count matrices for 2,500 genes per dataset. The counts were generated by running the *kallisto*|*bustools* 0.26.0 kb count command on the raw FASTQs with the --lamanno option, using an intronic/exonic index built from the GRCh38 and mm10 reference genomes, as described in Section 6.8.2. The datasets were filtered to remove low-expression droplets, first using the default *bustools* filter, then using the manually selected knee plot thresholds shown in Table S5 of Gorin et al.^21^ Next, they were filtered for the top 2,500 moderate-to high-expression genes using the procedure in Section S4.3.1 of Gorin et al.^21^ To visualize the broad trends in count averages, we obtained the gene lengths *L*_*j*_, then binned the values of log_10_ *L*_*j*_ into ten bins, with the edges given by the deciles *d*_0_, *d*_1_, ⋯, *d*_10_. Next, we computed the average log_10_ mean of nascent and mature expression levels for genes falling into each bin. Finally, we plotted these mean levels at each bin center 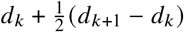, connecting the values with a line to guide the eye. We repeated this analysis for all twelve datasets, distinguishing the nascent and mature statistics by color.

Next, we obtained the fit results for these datasets. The fits were performed using *Monod* 0.2.5.0 Python package^133^ as described in Gorin et al.^21^ Fitting the model with no technical noise entailed gradient optimization over the per-gene joint distributions to fit *b*_*j*_, *β*_*j*_, and *γ*_*j*_. Although the model did not explicitly include technical noise, the theoretical discussion above implies that the results can be interpreted as those from a *p* = *p*_*N*_ = *p*_*M*_ model, with the inferred “burst size” corresponding to *b*_*j*_ *p*_*j*_ for gene *j*. Fitting the model with technical noise entailed scanning over a grid of *C*_*N*_ and *λ*_*M*_, obtaining per-gene maximum likelihood estimates of *b*_*j*_, *β*_*j*_, and *γ*_*j*_ *conditional on* the technical parameter values at the grid point, then identifying the grid point which produced the lowest sum of Kullback-Leibler divergences over all genes. In both cases, the genes underwent a round of goodness-of-fit filtering to remove fits that did not accurately recapitulate the data, as in Section S4.3.5 of Gorin et al.^21^. Next, we computed the average inferred log_10_ burst size for the genes falling into each length bin. As with the means, we plotted the average burst sizes at each bin center. connecting the values with a line to guide the eye. We repeated this analysis for all twelve datasets, distinguishing the results fit with and without a technical noise component by color.

## Supplementary Tables

**Table S1.** Components of the full master equation.

| Term | Interpretation |
| --- | --- |
| $H_{is}(t)P(i, \mathbf{x}, \mathbf{y}, t)$ | Transition from categorical state $i$ to $s$ |
| $c_{i0} [(x_i + 1)P(s, x_i + 1, \mathbf{y}, t) - x_i P(s, \mathbf{x}, \mathbf{y}, t)]$ | Degradation of discrete species $i$ |
| $c_{ij} [(x_i + 1)P(s, x_i + 1, x_j - 1, \mathbf{y}, t) - x_i P(s, \mathbf{x}, \mathbf{y}, t)]$ | Conversion of discrete species $i$ to $j$ |
| $Q_{ii}^d [(x_i - 1)P(s, x_i - 1, \mathbf{y}, t) - x_i P(s, \mathbf{x}, \mathbf{y}, t)]$ | Autocatalysis of discrete species $i$ |
| $Q_{ji}^d [x_i P(s, x_j - 1, \mathbf{y}, t) - x_i P(s, \mathbf{x}, \mathbf{y}, t)]$ | Catalysis of discrete species $j$ by $i$ |
| $\alpha_{s,\omega}^d(t) [\sum_{\mathbf{z}} p_{s,\omega}^d(\mathbf{z}, t) P(s, \mathbf{x} - \mathbf{z}, \mathbf{y}, t) - P(s, \mathbf{x}, \mathbf{y}, t)]$ | Bursty production of discrete species |
| $-C_{ji}^{cc} \frac{\partial}{\partial y_j} [y_i P(s, \mathbf{x}, \mathbf{y}, t)]$ | Increase in continuous species $j$<br>proportional to level of continuous species $i$ |
| $\sigma_i^2 \frac{\partial^2}{\partial y_i^2} [y_i P(s, \mathbf{x}, \mathbf{y}, t)]$ | Square-root noise in continuous species $i$ |
| $-\alpha_{s,i}^c(t) \frac{\partial P(s, \mathbf{x}, \mathbf{y}, t)}{\partial y_i}$ | Drift in continuous species $i$ |
| $\alpha_{s,\omega}^c(t) [\int_{\mathbf{z}} p_{\omega}^c(\mathbf{z}) P(\mathbf{y} - \mathbf{z}, t) d\mathbf{z} - P(\mathbf{y})]$ | Bursting in continuous species |
| $C_{ji}^{cd} [y_i P(x_j - 1, \mathbf{y}, t) - y_i P(\mathbf{x}, \mathbf{y}, t)]$ | Production of discrete species $j$<br>proportional to level of continuous species $i$ |
| $-C_{ji}^{dc} x_i \frac{\partial P(\mathbf{x}, \mathbf{y}, t)}{\partial y_j}$ | Drift in continuous species $j$<br>proportional to number of discrete species $i$ |

**Table S2.**
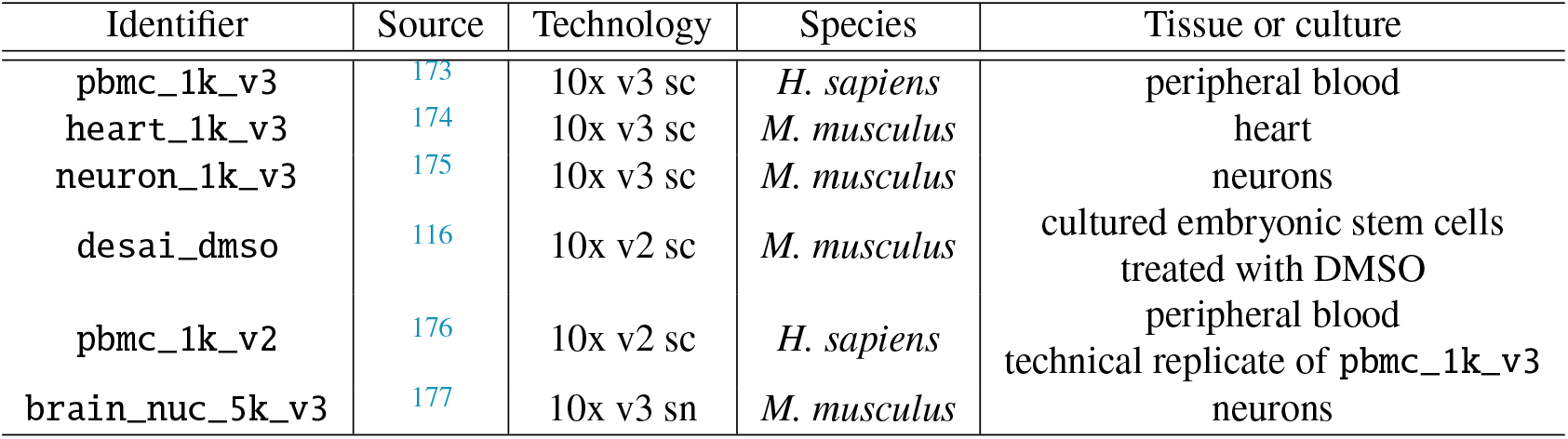
Datasets used for empty droplet analysis (sc: single-cell; sn: single-nucleus).

**Table S3** Genes discovered to be overdispersed (*σ*^2^ > 2*μ*) in empty droplets for each dataset in Table S2. [Table provided in external spreadsheet]

**Table S4** Genes discovered to be overdispersed (*σ*^2^ > 2*μ*) in empty droplets for the `neuron_1k_v3` and `desai_dmso` datasets, with function annotations. [Table provided in external spreadsheet]

