## Supplemental Information for "Studying stochastic systems biology of the cell with single-cell genomics data"

#### S1 MASTER EQUATION DERIVATIONS

One of the first steps in our generating function approach is to convert the master equation (which, in general, is given by Equation 45) satisfied by $P(s, \mathbf{x}, \mathbf{y}, t)$ to an equivalent partial differential equation satisfied by the generating function $\mathbf{G}$, whose components are

$$G_s = \int_{\mathbf{y}} \sum_{\mathbf{x}} \mathbf{g}^{\mathbf{x}} e^{\mathbf{h}^T \mathbf{y}} P(s, \mathbf{x}, \mathbf{y}, t) d\mathbf{y}. \tag{S1}$$

Because the master equation is linear, each of its terms will correspond to a particular term in the generating function PDE, and vice versa. One can develop a "dictionary" that allows one to go back and forth between these two different descriptions of the same system.

To see a simple and concrete example of this, let us consider the construction of the generating function PDE for the constitutive model (see Section 3). Recall that the CME and GF are

$$\begin{aligned} \frac{\partial P(x,t)}{\partial t} &= K\left[P(x-1,t) - P(x,t)\right] + \gamma\left[(x+1)P(x+1,t) - xP(x,t)\right], \\ G(g,t) &:= \sum_{x=0}^{\infty} g^x \, P(x,t). \end{aligned} \tag{S2}$$

Consider applying the generating function transformation (multiplying by $g^x$ and summing over $x \in \mathbb{N}_0$) on both sides of the above CME. We obtain

$$\sum_{x=0}^{\infty} g^x \, \frac{\partial P(x,t)}{\partial t} = K \sum_{x=0}^{\infty} g^x \, \left[P(x-1,t) - P(x,t)\right] + \gamma \sum_{x=0}^{\infty} g^x \, \left[(x+1)P(x+1,t) - xP(x,t)\right]. \tag{S3}$$

Completing the conversion requires noticing how to convert each individual piece. In particular:

$$\begin{aligned} \sum_{x=0}^{\infty} g^x \, \frac{\partial P(x,t)}{\partial t} &= \frac{\partial G(g,t)}{\partial t}, \\ \sum_{x=0}^{\infty} g^x \, \left[P(x-1,t) - P(x,t)\right] &= \sum_{x=0}^{\infty} [g-1] \, g^x \, P(x,t) = (g-1)G(g,t), \\ \sum_{x=0}^{\infty} g^x \, \left[(x+1)P(x+1,t) - xP(x,t)\right] &= \sum_{x=0}^{\infty} [1-g] \, \frac{\partial}{\partial g} g^x \, P(x,t) = -(g-1)\frac{\partial G(g,t)}{\partial g}. \end{aligned} \tag{S4}$$

This yields the PDE

$$\frac{\partial G}{\partial t} = (g-1)\left[KG - \frac{\partial G}{\partial g}\right]. \tag{S5}$$

This procedure can also be applied to much more complicated systems. In what follows, we will show how to convert each of the terms in Equation 45 into the corresponding PDE term, and hence construct the desired generating function PDE for our most general stochastic model.

A note on notation: whenever the argument of $P$ is not explicitly specified, it consists of $s$, $\mathbf{x}$, $\mathbf{y}$, and $t$. Similarly, whenever the argument of $G$ is not explicitly specified, it consists of $s$, $\mathbf{g}$, $\mathbf{h}$, and $t$.

##### S1.1 Fully discrete master equation terms

Multiplying $G$ through by $g_i$, we obtain:

$$g_i G = \int_{\mathbf{y}} \sum_{\mathbf{x}} \mathbf{g}^{\mathbf{x}} e^{\mathbf{h}^T \mathbf{y}} g_i P d\mathbf{y} = \int_{\mathbf{y}} \sum_{\mathbf{x}} \mathbf{g}^{\mathbf{x}} e^{\mathbf{h}^T \mathbf{y}} P(x_i - 1) d\mathbf{y}. \tag{S6}$$

This follows from rewriting $\sum_{x_i=0}^{\infty} g_i^{x_i+1} P(x_i)$ as $\sum_{x_i=-1}^{\infty} g_i^{x_i+1} P(x_i)$, noting that $P(x_i) = 0$ whenever $x_i < 0$, and reindexing to obtain the equivalent sum $\sum_{x_i=0}^{\infty} g_i^{x_i} P(x_i - 1)$. Therefore, the generating function of all terms that scale as $P(x_i - 1)$ is $g_i G$.

Differentiating with respect to $g_i$:

$$\begin{aligned}\frac{\partial G}{\partial g_i} &= \int_{\mathbf{y}} \sum_{\mathbf{x}} \mathbf{g}^{\mathbf{x}} e^{\mathbf{h}^T\mathbf{y}} x_i g_i^{-1} P d\mathbf{y}, \text{ i.e.,} \\ g_i \frac{\partial G}{\partial g_i} &= \int_{\mathbf{y}} \sum_{\mathbf{x}} \mathbf{g}^{\mathbf{x}} e^{\mathbf{h}^T\mathbf{y}} x_i P d\mathbf{y}.\end{aligned} \tag{S7}$$

Therefore, the generating function of all terms that scale as $x_i P$ is $g_i \frac{\partial G_s}{\partial g_i}$.

Alternatively, we can use the fact that the $x_i = 0$ term of $\sum_{x_i=0}^{\infty} g_i^{x_i-1} x_i P(x_i)$ is zero to shift the summand by one: $\sum_{x_i=0}^{\infty} g_i^{x_i}(x_i+1)P(x_i+1)$. Therefore, the generating function of all terms that scale as $(x_i+1)P(x_i+1)$ is $\frac{\partial G_s}{\partial g_i}$.

Multiplying this equation through by $g_j$, we obtain:

$$\begin{aligned}g_j \frac{\partial G}{\partial g_i} &= \int_{\mathbf{y}} \sum_{\mathbf{x}} \mathbf{g}^{\mathbf{x}} e^{\mathbf{h}^T\mathbf{y}} g_j (x_i+1) P(x_i+1) d\mathbf{y} \\ &= \int_{\mathbf{y}} \sum_{\mathbf{x}} \mathbf{g}^{\mathbf{x}} e^{\mathbf{h}^T\mathbf{y}} (x_i+1) P(x_i+1, x_j-1) d\mathbf{y}.\end{aligned} \tag{S8}$$

This follows from rewriting $\sum_{x_j=0}^{\infty} g_j^{x_j+1} P(x_i+1, j)$ as $\sum_{x_j=-1}^{\infty} g_j^{x_j+1} P(x_i+1, x_j)$, noting that $P(x_j) = 0$ whenever $x_j < 0$, and reindexing to obtain the equivalent sum $\sum_{x_j=0}^{\infty} g_j^{x_j} P(x_i+1, x_j-1)$. Therefore, the generating function of all terms that scale as $(x_i+1)P(x_i+1, x_j-1)$ is $g_j \frac{\partial G}{\partial g_i}$.

Multiplying the derivative by $g_i$ twice, we find:

$$\begin{aligned}g_i^2 \frac{\partial G}{\partial g_i} &= \int_{\mathbf{y}} \sum_{\mathbf{x}} \mathbf{g}^{\mathbf{x}} e^{\mathbf{h}^T\mathbf{y}} x_i g_i P d\mathbf{y} \\ &= \int_{\mathbf{y}} \sum_{\mathbf{x}} \mathbf{g}^{\mathbf{x}} e^{\mathbf{h}^T\mathbf{y}} (x_i-1) P(x_i-1) d\mathbf{y}.\end{aligned} \tag{S9}$$

This follows from rewriting $\sum_{x_i=0}^{\infty} g_i^{x_i+1} x_i P(x_i)$ as $\sum_{x_i=-1}^{\infty} g_i^{x_i+1} x_i P(x_i)$, noting that $P(x_i) = 0$ whenever $x_i < 0$, and reindexing to obtain the equivalent sum $\sum_{x_i=0}^{\infty} g_i^{x_i}(x_i-1)P(x_i-1)$. Therefore, the generating function of all terms that scale as $(x_i-1)P(x_i-1)$ is $g_i^2 \frac{\partial G}{\partial g_i}$.

Multiplying the derivative by $g_i g_j$ we obtain:

$$\begin{aligned}g_i g_j \frac{\partial G}{\partial g_i} &= g_j \int_{\mathbf{y}} \sum_{\mathbf{x}} \mathbf{g}^{\mathbf{x}} e^{\mathbf{h}^T\mathbf{y}} x_i P d\mathbf{y} \\ &= \int_{\mathbf{y}} \sum_{\mathbf{x}} \mathbf{g}^{\mathbf{x}} e^{\mathbf{h}^T\mathbf{y}} x_i g_j P d\mathbf{y} \\ &= \int_{\mathbf{y}} \sum_{\mathbf{x}} \mathbf{g}^{\mathbf{x}} e^{\mathbf{h}^T\mathbf{y}} x_i P(x_j-1) d\mathbf{y}.\end{aligned} \tag{S10}$$

This follows from rewriting $\sum_{x_i=0}^{\infty} g_j^{x_j+1} P(x_j)$ as $\sum_{x_j=-1}^{\infty} g_j^{x_j+1} P(x_j)$, noting that $P(x_j) = 0$ whenever $x_j < 0$, and reindexing to obtain the equivalent sum $\sum_{x_j=0}^{\infty} g_j^{x_j} P(x_j)$. Therefore, the generating function of all terms that scale as $x_i P(x_j-1)$ is $g_i g_j \frac{\partial G}{\partial g_i}$.

Multiplying the generating function by a probability-generating function $F$ of a discrete burst distribution $p$ yields:

$$\begin{aligned}FG &= \int_{\mathbf{y}} \sum_{\mathbf{x}} \mathbf{g}^{\mathbf{x}} e^{\mathbf{h}^T\mathbf{y}} F P d\mathbf{y} \\ &= \int_{\mathbf{y}} \sum_{\mathbf{x}} \mathbf{g}^{\mathbf{x}} e^{\mathbf{h}^T\mathbf{y}} \sum_{\mathbf{z}} p(\mathbf{z}) P(\mathbf{x}-\mathbf{z}) d\mathbf{y}.\end{aligned} \tag{S11}$$

This identity may be derived from three directions. The simplest approach is to note that it follows from the convolution theorem[178]. Alternatively, it may be proven directly using the repeated ($n$-fold) application of Cauchy products[16]. Finally, the master equation term essentially aggregates two independent random variables—the process values and the burst sizes—whose sum is equal to $\mathbf{x}$. The generating function of the sum of independent variates is the product of their generating functions. Therefore, the generating function of all the terms that scale as $\sum_{\mathbf{z}} p(\mathbf{z}) P(\mathbf{x}-\mathbf{z})$ is $FG$.

#### S1.2 Fully continuous master equation terms

Multiplying $G$ through by $h_i$, we obtain:

$$\begin{aligned} h_i G &= \int_{\mathbf{y}} \sum_{\mathbf{x}} \mathbf{g}^{\mathbf{x}} e^{\mathbf{h}^T \mathbf{y}} h_i P d\mathbf{y} = \int_{\mathbf{y}} \sum_{\mathbf{x}} \mathbf{g}^{\mathbf{x}} \frac{\partial}{\partial y_i} \left[ e^{\mathbf{h}^T \mathbf{y}} \right] [P] \, d\mathbf{y} \\ &= - \int_{\mathbf{y}} \sum_{\mathbf{x}} \mathbf{g}^{\mathbf{x}} e^{\mathbf{h}^T \mathbf{y}} \frac{\partial P}{\partial y_i} d\mathbf{y}. \end{aligned} \tag{S12}$$

This follows from integration by parts. The product term $e^{\mathbf{h}^T \mathbf{y}} P$ does not contribute to this expression because $P$ is a density with zero mass at any particular value of $\mathbf{y}$. Therefore, the generating function of all terms that scale as $\frac{\partial P}{\partial y_i}$ is $-h_i G$.

Differentiating with respect to $h_i$, we obtain:

$$\frac{\partial G}{\partial h_i} = \int_{\mathbf{y}} \sum_{\mathbf{x}} \mathbf{g}^{\mathbf{x}} e^{\mathbf{h}^T \mathbf{y}} y_i P d\mathbf{y}. \tag{S13}$$

Therefore, the generating function of all terms that scale as $y_i P$ is $\frac{\partial G_s}{\partial h_i}$.

Multiplying through by $h_j$ yields:

$$\begin{aligned} h_j \frac{\partial G}{\partial h_i} &= \int_{\mathbf{y}} \sum_{\mathbf{x}} \mathbf{g}^{\mathbf{x}} e^{\mathbf{h}^T \mathbf{y}} h_j y_i P d\mathbf{y} = \int_{\mathbf{y}} \sum_{\mathbf{x}} \mathbf{g}^{\mathbf{x}} \frac{\partial}{\partial y_j} \left[ e^{\mathbf{h}^T \mathbf{y}} \right] [y_i P] \, d\mathbf{y} \\ &= - \int_{\mathbf{y}} \sum_{\mathbf{x}} \mathbf{g}^{\mathbf{x}} e^{\mathbf{h}^T \mathbf{y}} \frac{\partial \left[ y_i P \right]}{\partial y_j} d\mathbf{y}. \end{aligned} \tag{S14}$$

This also follows from integration by parts. The product term $e^{\mathbf{h}^T \mathbf{y}} y_i P$ does not contribute to this expression because it is identically zero at $y_i = 0$ and $P$ vanishes as $y_i \to \infty$. Therefore, the generating function of all terms that scale as $\frac{\partial [y_i P]}{\partial y_j}$ is $-h_j \frac{\partial G}{\partial h_i}$.

Multiplying the derivative by $h_i$ twice results in:

$$\begin{aligned} h_i^2 \frac{\partial G}{\partial h_i} &= -h_i \int_{\mathbf{y}} \sum_{\mathbf{x}} \mathbf{g}^{\mathbf{x}} e^{\mathbf{h}^T \mathbf{y}} \frac{\partial \left[ y_i P \right]}{\partial y_i} d\mathbf{y} \\ &= - \int_{\mathbf{y}} \sum_{\mathbf{x}} \mathbf{g}^{\mathbf{x}} \frac{\partial}{\partial y_i} \left[ e^{\mathbf{h}^T \mathbf{y}} \right] \frac{\partial \left[ y_i P \right]}{\partial y_i} d\mathbf{y} \\ &= \int_{\mathbf{y}} \sum_{\mathbf{x}} \mathbf{g}^{\mathbf{x}} e^{\mathbf{h}^T \mathbf{y}} \frac{\partial^2 \left[ y_i P \right]}{\partial y_i^2} d\mathbf{y}. \end{aligned} \tag{S15}$$

Again, this follows from integration by parts. Moreover, the product term $e^{\mathbf{h}^T \mathbf{y}} \frac{\partial [y_i P]}{\partial y_i} = e^{\mathbf{h}^T \mathbf{y}} y_i \frac{\partial P}{\partial y_i} + e^{\mathbf{h}^T \mathbf{y}} P$ again does not contribute to this expression because $P$ is a density that vanishes as $y_i \to \infty$. Therefore, the generating function of all terms that scale as $\frac{\partial^2 [y_i P]}{\partial y_i^2}$ is $h_i^2 \frac{\partial G}{\partial h_i}$.

Multiplying the generating function by a moment-generating function $M$ of continuous burst distribution $p$ leads to:

$$MG = \int_{\mathbf{y}} \sum_{\mathbf{x}} \mathbf{g}^{\mathbf{x}} e^{\mathbf{h}^T \mathbf{y}} M P d\mathbf{y} = \int_{\mathbf{y}} \sum_{\mathbf{x}} \mathbf{g}^{\mathbf{x}} e^{\mathbf{h}^T \mathbf{y}} \int_{\mathbf{z}} p(\mathbf{z}) P(\mathbf{y} - \mathbf{z}) d\mathbf{z} d\mathbf{y}, \tag{S16}$$

which may be derived from the convolution theorem, or MGF identities, identically to Equation S11. Therefore, the generating function of all terms that scale as $p(\mathbf{z}) P(\mathbf{y} - \mathbf{z}) d\mathbf{z}$ is $MG$.

#### S1.3 Mixed master equation terms

Considering the case where a continuous process drives a discrete one, and multiplying $\frac{\partial G}{\partial h_i}$ by $g_j$ we obtain:

$$\begin{aligned} g_j \frac{\partial G}{\partial h_i} &= \int_{\mathbf{y}} \sum_{\mathbf{x}} \mathbf{g}^{\mathbf{x}} e^{\mathbf{h}^T \mathbf{y}} y_i g_j P d\mathbf{y} \\ &= \int_{\mathbf{y}} \sum_{\mathbf{x}} \mathbf{g}^{\mathbf{x}} e^{\mathbf{h}^T \mathbf{y}} y_i P(x_j - 1) d\mathbf{y}. \end{aligned} \tag{S17}$$

The derivation is identical to Equation S6. Therefore, the generating function of all terms that scale as $y_i P(x_j - 1)$ is $g_j \frac{\partial G}{\partial h_i}$.

Considering the case where a discrete process drives a continuous one, and multiplying $g_i \frac{\partial G}{\partial g_i}$ by $h_j$ yields:

$$\begin{aligned} h_j g_i \frac{\partial G}{\partial g_i} &= \int_{\mathbf{y}} \sum_{\mathbf{x}} \mathbf{g}^{\mathbf{x}} e^{\mathbf{h}^T \mathbf{y}} x_i h_j P d\mathbf{y} \\ &= \int_{\mathbf{y}} \sum_{\mathbf{x}} \mathbf{g}^{\mathbf{x}} \frac{\partial}{\partial y_j} \left[ e^{\mathbf{h}^T \mathbf{y}} \right] [x_i P] \, d\mathbf{y} \\ &= - \int_{\mathbf{y}} \sum_{\mathbf{x}} \mathbf{g}^{\mathbf{x}} e^{\mathbf{h}^T \mathbf{y}} \frac{\partial [x_i P]}{\partial y_j} d\mathbf{y}. \end{aligned} \tag{S18}$$

This follows from integration by parts; as before, the product term does not appear because $P$ is a density. Therefore, the generating function of all terms that scale as $\frac{\partial [x_i P]}{\partial y_j} = x_i \frac{\partial P}{\partial y_j}$ is $-h_j g_i \frac{\partial G}{\partial g_i}$. This concludes the enumeration of generating function identities.

#### S1.4 Converting the master equation to a partial differential equation

By exploiting the identities derived above and the linearity of the generating function, we can represent Equation 45 by an equivalent deterministic partial differential equation. We begin by considering the expressions for each entry of $\mathbf{G}$ separately, eliding the gene state $s$.

Each entry of the second term on the right-hand side, which represents degradation of the discrete species, takes the form

$$\begin{aligned} &\int_{\mathbf{y}} \sum_{\mathbf{x}} \mathbf{g}^{\mathbf{x}} e^{\mathbf{h}^T \mathbf{y}} \left[ (x_i + 1) P(x_i + 1) - x_i P \right] d\mathbf{y} \\ &= \frac{\partial G}{\partial g_i} - g_i \frac{\partial G}{\partial g_i} = (1 - g_i) \frac{\partial G}{\partial g_i}, \text{ yielding} \\ &\sum_{i=1}^{n} c_{i0} (1 - g_i) \frac{\partial G}{\partial g_i}. \end{aligned} \tag{S19}$$

Each entry of the third term, which represents interconversion of the discrete species, takes the form

$$\begin{aligned} &\int_{\mathbf{y}} \sum_{\mathbf{x}} \mathbf{g}^{\mathbf{x}} e^{\mathbf{h}^T \mathbf{y}} \left[ (x_i + 1) P(x_i + 1, x_j - 1) - x_i P \right] d\mathbf{y} \\ &= g_j \frac{\partial G}{\partial g_i} - g_i \frac{\partial G}{\partial g_i} = (g_j - g_i) \frac{\partial G}{\partial g_i}, \text{ yielding} \\ &\sum_{i,j=1}^{n} c_{ij} (g_j - g_i) \frac{\partial G}{\partial g_i}. \end{aligned} \tag{S20}$$

Each entry of the fourth term, which represents autocatalysis of the discrete species, takes the form

$$\begin{aligned} &\int_{\mathbf{y}} \sum_{\mathbf{x}} \mathbf{g}^{\mathbf{x}} e^{\mathbf{h}^T \mathbf{y}} \left[ (x_i - 1) P(x_i - 1) - x_i P \right] d\mathbf{y} \\ &= g_i^2 \frac{\partial G}{\partial g_i} - g_i \frac{\partial G}{\partial g_i} = (g_i - 1) g_i \frac{\partial G}{\partial g_i}, \text{ yielding} \\ &\sum_{i=1}^{n} Q_{ii}^d (g_i - 1) g_i \frac{\partial G}{\partial g_i}. \end{aligned} \tag{S21}$$

Each entry of the fifth term, which represents catalysis of the discrete species, takes the form

$$\begin{aligned} &\int_{\mathbf{y}} \sum_{\mathbf{x}} \mathbf{g}^{\mathbf{x}} e^{\mathbf{h}^T \mathbf{y}} \left[ x_i P(x_j - 1) - x_i P \right] d\mathbf{y} \\ &= g_i g_j \frac{\partial G}{\partial g_i} - g_i \frac{\partial G}{\partial g_i} = (g_j - 1) g_i \frac{\partial G}{\partial g_i}, \text{ yielding} \\ &\sum_{i,j=1}^{n} Q_{ji}^d (g_j - 1) g_i \frac{\partial G}{\partial g_i}. \end{aligned} \tag{S22}$$

Each entry of the sixth term, which represents bursty production of the discrete species, takes the form

$$\begin{aligned} &\int_{\mathbf{y}} \sum_{\mathbf{x}} \mathbf{g}^{\mathbf{x}} e^{\mathbf{h}^T \mathbf{y}} \left[ \sum_{\mathbf{z}} p^d_{s,\omega}(\mathbf{z}, t) P(\mathbf{x} - \mathbf{z}) - P \right] d\mathbf{y} \\ &= FG - G = (F-1)G, \text{ yielding} \\ &\sum_{\omega} \alpha^d_{\omega} (F_\omega - 1) G. \end{aligned} \tag{S23}$$

Each entry of the seventh term, which represents the deterministic dynamics of the continuous species, takes the form

$$\begin{aligned} &\int_{\mathbf{y}} \sum_{\mathbf{x}} \mathbf{g}^{\mathbf{x}} e^{\mathbf{h}^T \mathbf{y}} \frac{\partial}{\partial y_j} [y_i P] \, d\mathbf{y} \\ &= -h_j \frac{\partial G}{\partial h_i}, \text{ yielding} \\ &\sum_{i,j=1}^{m} C^{cc}_{ji} h_j \frac{\partial G}{\partial h_i}. \end{aligned} \tag{S24}$$

Each entry of the eighth term, which represents the diffusion dynamics of the continuous species, takes the form

$$\begin{aligned} &\int_{\mathbf{y}} \sum_{\mathbf{x}} \mathbf{g}^{\mathbf{x}} e^{\mathbf{h}^T \mathbf{y}} \frac{\partial^2}{\partial y_i^2} [y_i P] \, d\mathbf{y} \\ &= h_i^2 \frac{\partial G}{\partial h_i}, \text{ yielding} \\ &\frac{1}{2} \sum_{i=1}^{m} \sigma_i^2 h_i^2 \frac{\partial G}{\partial h_i}. \end{aligned} \tag{S25}$$

Each entry of the ninth term, which represents the drift of the continuous species, takes the form

$$\begin{aligned} &\int_{\mathbf{y}} \sum_{\mathbf{x}} \mathbf{g}^{\mathbf{x}} e^{\mathbf{h}^T \mathbf{y}} \frac{\partial P}{\partial y_i} d\mathbf{y} \\ &= -h_i G, \text{ yielding} \\ &\sum_{i=1}^{m} \alpha^c_i h_i G. \end{aligned} \tag{S26}$$

Each entry of the tenth term, which represents the bursty production of the continuous species, takes the form

$$\begin{aligned} &\int_{\mathbf{y}} \sum_{\mathbf{x}} \mathbf{g}^{\mathbf{x}} e^{\mathbf{h}^T \mathbf{y}} \left[ \int_{\mathbf{z}} p^c_{\omega}(\mathbf{z}) P(\mathbf{y} - \mathbf{z}, t) d\mathbf{z} - P \right] d\mathbf{y} \\ &= MG - G = (M-1)G, \text{ yielding} \\ &\sum_{\omega > m} \alpha^c_{\omega} (M_\omega - 1) G. \end{aligned} \tag{S27}$$

Each entry of the eleventh term, which represents a continuous species driving a discrete one, takes the form

$$\begin{aligned} &\int_{\mathbf{y}} \sum_{\mathbf{x}} \mathbf{g}^{\mathbf{x}} e^{\mathbf{h}^T \mathbf{y}} \left[ y_i P(x_j - 1) - y_i P \right] d\mathbf{y} \\ &= g_j \frac{\partial G}{\partial h_i} - \frac{\partial G}{\partial h_i} = (g_j - 1) \frac{\partial G}{\partial h_i}, \text{ yielding} \\ &\sum_{i=1}^{m} \sum_{j=1}^{n} C^{cd}_{ji} (g_j - 1) \frac{\partial G}{\partial h_i}. \end{aligned} \tag{S28}$$

Each entry of the twelfth term, which represents a discrete species driving a continuous one, takes the form

$$\begin{aligned}&\int_{\mathbf{y}}\sum_{\mathbf{x}}\mathbf{g}^{\mathbf{x}}e^{\mathbf{h}^T\mathbf{y}}x_i\frac{\partial P}{\partial y_j}d\mathbf{y}\\&=h_jg_i\frac{\partial G}{\partial g_i},\text{ yielding}\\&\sum_{i=1}^{n}\sum_{j=1}^{m}C_{ji}^{dc}h_jg_i\frac{\partial G}{\partial g_i}.\end{aligned}\tag{S29}$$

Therefore, the PDE form of the master equation is

$$\begin{aligned}\frac{\partial G}{\partial t}=&\sum_{i=1}^{N}H_{is}G_i+\sum_{i=1}^{n}c_{i0}(1-g_i)\frac{\partial G}{\partial g_i}+\sum_{i,j=1}^{n}c_{ij}(g_j-g_i)\frac{\partial G}{\partial g_i}\\&+\sum_{i=1}^{n}Q_{ii}^{d}(g_i-1)g_i\frac{\partial G}{\partial g_i}+\sum_{i,j=1}^{n}Q_{ji}^{d}(g_j-1)g_i\frac{\partial G}{\partial g_i}+\sum_{\omega}\alpha_{\omega}^{d}(F_\omega-1)G\\&+\sum_{i,j=1}^{m}C_{ji}^{cc}h_j\frac{\partial G}{\partial h_i}+\frac{1}{2}\sum_{i=1}^{m}\sigma_i^2h_i^2\frac{\partial G}{\partial h_i}+\sum_{i=1}^{m}\alpha_i^{c}h_iG+\sum_{\omega>m}\alpha_{\omega}^{c}(M_\omega-1)G\\&+\sum_{i=1}^{m}\sum_{j=1}^{n}C_{ji}^{cd}(g_j-1)\frac{\partial G}{\partial h_i}+\sum_{i=1}^{n}\sum_{j=1}^{m}C_{ji}^{dc}h_jg_i\frac{\partial G}{\partial g_i}.\end{aligned}\tag{S30}$$

This equation governs the dynamics of $G_s$, one of $N$ coupled PDEs. We elide this subscript when it does not directly factor into the calculation. As in Equation 45, the terms that scale with $G$, rather than one of its derivatives, may be time- and $s$-dependent.

#### S1.5 Representing the PDE in matrix form

This formulation in Equation S30 is somewhat more compact, but may be simplified further. First, we note that the second and third terms on the right-hand side can be represented by the matrix equation

$$(\nabla^d G)^T(C^{dd})^T(\mathbf{g}-1),\tag{S31}$$

where $\nabla^d$ is the gradient with respect to entries of $\mathbf{g}$.

Next, the fourth and fifth terms can be represented by

$$(\nabla^d G)^T(\text{diag }\mathbf{g})(Q^d)^T(\mathbf{g}-1).\tag{S32}$$

The sixth term can be represented by constructing vectors indexed by $\omega$:

$$G(\boldsymbol{\alpha}^d)^T(\mathbf{F}(\mathbf{g})-1).\tag{S33}$$

The seventh term can be represented similarly to second and third:

$$(\nabla^c G)^T(C^{cc})^T\mathbf{h},\tag{S34}$$

where $\nabla^c$ is the gradient with respect to entries of $\mathbf{h}$.

The eighth term can be represented analogously to the fourth and fifth:

$$(\nabla^c G)^T(\text{diag }\mathbf{h})\left(\frac{1}{2}\text{diag }\boldsymbol{\sigma}^2\right)\mathbf{h}.\tag{S35}$$

The ninth and tenth term can be represented analogously to the sixth:

$$G(\boldsymbol{\alpha}^c)^T(\mathbf{M}(\mathbf{h})-1),\tag{S36}$$

where $\mathbf{M}$ is a vector-valued function indexed by $\omega$. The first $m$ entries of $\boldsymbol{\alpha}^c$ contain the $m$ scalar drift rates, whereas the other entries contain jump rates, i.e., $(\mathbf{M})_i := h_i + 1$ for $i \leq m$.

The eleventh term takes a form analogous to those for second, third, and seventh:

$$(\nabla^c G)^T (C^{cd})^T (\mathbf{g} - 1). \tag{S37}$$

Finally, the twelfth is analogous to fourth and fifth:

$$(\nabla^d G)^T (\operatorname{diag} \mathbf{g})(C^{dc})^T \mathbf{h}. \tag{S38}$$

Therefore, Equation S30 can be condensed further for a particular $s$:

$$\begin{aligned}\frac{\partial G}{\partial t} = \sum_{i=1}^{N} H_{is} G_i &+ (\nabla^d G)^T (C^{dd})^T (\mathbf{g} - 1) + (\nabla^d G)^T (\operatorname{diag} \mathbf{g})(Q^d)^T (\mathbf{g} - 1) + G(\boldsymbol{\alpha}^d)^T (\mathbf{F}(\mathbf{g}) - 1) \\ &+ (\nabla^c G)^T (C^{cc})^T \mathbf{h} + (\nabla^c G)^T (\operatorname{diag} \mathbf{h}) \left(\frac{1}{2} \operatorname{diag} \boldsymbol{\sigma}^2\right) (\mathbf{h}) + G(\boldsymbol{\alpha}^c)^T (\mathbf{M}(\mathbf{h}) - 1) \\ &+ (\nabla^c G)^T (C^{cd})^T (\mathbf{g} - 1) + (\nabla^d G)^T (\operatorname{diag} \mathbf{g})(C^{dc})^T \mathbf{h}.\end{aligned} \tag{S39}$$

Collecting terms:

$$\begin{aligned}\frac{\partial G}{\partial t} &= (\nabla^d G)^T \left[(C^{dd})^T (\mathbf{g} - 1) + (\operatorname{diag} \mathbf{g})(Q^d)^T (\mathbf{g} - 1) + (\operatorname{diag} \mathbf{g})(C^{dc})^T \mathbf{h}\right] \\ &+ (\nabla^c G)^T \left[(C^{cc})^T \mathbf{h} + (\operatorname{diag} \mathbf{h}) \left(\frac{1}{2} \operatorname{diag} \boldsymbol{\sigma}^2\right) \mathbf{h} + (C^{cd})^T (\mathbf{g} - 1)\right] \\ &+ G\left[(\boldsymbol{\alpha}^d)^T (\mathbf{F}(\mathbf{g}) - 1) + (\boldsymbol{\alpha}^c)^T (\mathbf{M}(\mathbf{h}) - 1)\right] + \sum_{i=1}^{N} H_{is} G_i.\end{aligned} \tag{S40}$$

In other words, the PDE separates into the expected first-order linear form, with terms corresponding to $G$, $\nabla^d G$, and $\nabla^c G$.

#### S1.6 Unifying the discrete and continuous species

Analyzing Equation S40 as is obfuscates the mathematical similarities of the discrete and continuous species.

To exploit them, we first introduce the variable $\mathbf{u}$, which is a shifted version of $\mathbf{g}$ concatenated to $\mathbf{h}$:

$$\mathbf{u} := \begin{bmatrix} \mathbf{u}_d \\ \mathbf{u}_c \end{bmatrix} = \begin{bmatrix} \mathbf{g} - 1 \\ \mathbf{h} \end{bmatrix}. \tag{S41}$$

This yields the following form for the gradient-dependent terms:

$$\begin{aligned}&(\nabla^d G)^T \left[(C^{dd})^T \mathbf{u}_d + (\operatorname{diag} \mathbf{u}_d + I)(Q^d)^T \mathbf{u}_d + (\operatorname{diag} \mathbf{u}_d + I)(C^{dc})^T \mathbf{u}_c\right] \\ &+ (\nabla^c G)^T \left[(C^{cc})^T \mathbf{u}_c + (\operatorname{diag} \mathbf{u}_c) \left(\frac{1}{2} \operatorname{diag} \boldsymbol{\sigma}^2\right) \mathbf{u}_c + (C^{cd})^T \mathbf{u}_d\right] \\ &= (\nabla^d G)^T \left[\left((C^{dd})^T + (Q^d)^T\right) \mathbf{u}_d + (C^{dc})^T \mathbf{u}_c + (\operatorname{diag} \mathbf{u}_d)(Q^d)^T \mathbf{u}_d + (\operatorname{diag} \mathbf{u}_d)(C^{dc})^T \mathbf{u}_c\right] \\ &+ (\nabla^c G)^T \left[(C^{cd})^T \mathbf{u}_d + (C^{cc})^T \mathbf{u}_c + (\operatorname{diag} \mathbf{u}_c) \left(\frac{1}{2} \operatorname{diag} \boldsymbol{\sigma}^2\right) \mathbf{u}_c\right].\end{aligned} \tag{S42}$$

Next, we define the full gradient of $G$, such that $\nabla G$ contains the derivatives with respect to all entries of $\mathbf{u}$. We define common jump rates and generating functions:

$$\begin{aligned}\boldsymbol{\alpha} &:= \begin{bmatrix} \boldsymbol{\alpha}^d \\ \boldsymbol{\alpha}^c \end{bmatrix}, \\ \mathcal{M}(\mathbf{u}) &:= \begin{bmatrix} F(1 + \mathbf{u}_{1,\cdots,n}) \\ M(\mathbf{u}_{n+1,\cdots,n+m}) \end{bmatrix}.\end{aligned} \tag{S43}$$

We define the common interconversion matrix:

$$C := \begin{bmatrix} (C^{dd})^T + (Q^d)^T & (C^{dc})^T \\ (C^{cd})^T & (C^{cc})^T \end{bmatrix}, \tag{S44}$$

as well as the common diffusion matrix:

$$D := \begin{bmatrix} (Q^d)^T & (C^{dc})^T \\ 0 & \frac{1}{2}\text{diag}\,\boldsymbol{\sigma}^2 \end{bmatrix} := \begin{bmatrix} (Q^d)^T & (C^{dc})^T \\ 0 & (Q^c)^T \end{bmatrix}. \tag{S45}$$

This yields the following unified expression for a single state:

$$\frac{\partial G}{\partial t} = (\nabla G)^T \left[C\mathbf{u} + \text{diag}\,\mathbf{u}\,D\mathbf{u}\right] + G\left[\boldsymbol{\alpha}^T(\mathcal{M}(\mathbf{y}) - 1)\right] + \sum_{i=1}^{N} H_{is} G_i. \tag{S46}$$

Only the final two terms differ between states. To characterize the bursting dynamics, we need to define the full bursting operator:

$$\mathcal{A}(\mathbf{u}) = \begin{bmatrix} \boldsymbol{\alpha}_1^T(\mathcal{M}_1(\mathbf{u}) - 1) \\ \cdots \\ \boldsymbol{\alpha}_N^T(\mathcal{M}_N(\mathbf{u}) - 1) \end{bmatrix}, \tag{S47}$$

where the subscripts of $\boldsymbol{\alpha}$ and $\mathcal{M}$ now indicate the gene state. Finally, recalling that the full Jacobian has entries $J_{si} = \frac{\partial G_s}{\partial u_i}$, the full PDE system takes the following form:

$$\frac{\partial \mathbf{G}}{\partial t} = H^T\mathbf{G} + \mathbf{G} \odot \mathcal{A}(\mathbf{u}) + J\left[C\mathbf{u} + \text{diag}\,\mathbf{u}\,D\mathbf{u}\right]. \tag{S48}$$

#### S1.7 Solving the partial differential equation

We seek to integrate this PDE to obtain the generating function at an arbitrary time $t$. The form is conducive to applying the method of characteristics. First, we define the characteristic variable $\mathsf{s}$. By taking a total derivative with respect to $\mathsf{s}$, we obtain

$$\frac{dG_s}{d\mathsf{s}} = \frac{\partial G_s}{\partial T}\frac{dT}{d\mathsf{s}} + \sum_{i=1}^{n+m} \frac{\partial G_s}{\partial U_i}\frac{dU_i}{d\mathsf{s}}. \tag{S49}$$

Next, we rewrite the PDE to match the form of the total derivative:

$$-H^T\mathbf{G} - \mathbf{G} \odot \mathcal{A}(\mathbf{u}) = -\frac{\partial \mathbf{G}}{\partial t} + J\left[C\mathbf{u} + \text{diag}\,\mathbf{u}\,D\mathbf{u}\right]. \tag{S50}$$

The characteristic curves emanating from $(t, \mathbf{u})$ that satisfy the PDE are given by:

$$\begin{aligned} \frac{dT(\mathsf{s})}{d\mathsf{s}} &= -1 \text{ such that } T(\mathsf{s}=0) = t, \text{ i.e., } T(\mathsf{s}) = t - \mathsf{s} \text{ and} \\ \frac{dU_i(\mathsf{s})}{d\mathsf{s}} &= \left(C\mathbf{U}(\mathsf{s}) + \text{diag}\,\mathbf{U}(\mathsf{s})\,D\mathbf{U}(\mathsf{s})\right)_i \text{ such that } \mathbf{U}(\mathsf{s}=0) = \mathbf{u}. \end{aligned} \tag{S51}$$

We denote the latter system as the "downstream" ODE, as it only involves the downstream system components. To solve the "upstream" ODE, we follow Equation 51.

### S2 POISSON REPRESENTATION ISOMORPHISMS

#### S2.1 Autocatalysis with constitutive production

Suppose we are interested in a 1-species system with $N = 1$, $n = 1$, $m = 0$, involving birth at $\alpha$, death at $\gamma$, and autocatalysis at $q$:

$$\begin{aligned} \varnothing &\xrightarrow{\alpha} \mathcal{X}, \\ \mathcal{X} &\xrightarrow{q} \mathcal{X} + \mathcal{X}, \\ \mathcal{X} &\xrightarrow{\gamma} \varnothing. \end{aligned} \tag{S52}$$

These reactions yield the following PDE terms:

$$\begin{aligned} \mathcal{M}(u) &= u, \\ \mathcal{A}(u) &= \alpha u, \\ C &= -\gamma + q, \\ D &= q. \end{aligned} \tag{S53}$$

The same form can be obtained by defining a system with $N = 1$, $n = 0$, and $m = 1$, with drift $\alpha$, mean-reversion at rate $\gamma - q$, and diffusion $q$. This system matches the functional form of a Cox–Ingersoll–Ross (CIR) process with drift $ab$, mean-reversion rate $a$ and square-root noise with intensity $\sigma$[164]:

$$\begin{aligned} dy_t &= a(b - y_t)dt + \sigma\sqrt{y_t}dW_t, \\ D = \frac{1}{2}\sigma^2 &= q \implies \sigma = \sqrt{2q}, \\ ab &= \alpha, \\ C = -a &= -\gamma + q. \end{aligned} \tag{S54}$$

As $t \longrightarrow \infty$, the CIR process approaches the gamma distribution with shape $\nu$ and scale $\theta$[164]:

$$\begin{aligned} P(y; \nu, \theta) &= \frac{1}{\Gamma(\nu)\theta^\nu} y^\nu e^{-y/\theta}, \\ \nu &= \frac{2ab}{\sigma^2} = \frac{\alpha}{q}, \\ \theta &= \frac{\sigma^2}{2a} = \frac{q}{\gamma - q}, \\ \text{with the MGF } M &= \left(\frac{1}{1 - \theta u}\right)^\nu \\ \text{resulting in the Poisson mixture PGF } G &= \left(\frac{1}{1 - \theta(g - 1)}\right)^\nu \\ \text{for the discrete law } P_{\text{NB}}(x; \nu, \theta) &= \frac{\Gamma(\nu + x)}{x!\Gamma(\nu)} \left(\frac{\theta}{1 + \theta}\right)^\nu \left(\frac{1}{1 + \theta}\right)^x. \end{aligned} \tag{S55}$$

This is the probability mass function of a negative binomial distribution with the shape $\nu = \alpha/q$ and mean $\nu\theta = \frac{\alpha}{\gamma - q}$. With some algebra, we can rewrite the PGF as

$$\left(\frac{\gamma - c}{\gamma}\right)^{\alpha/q} \left(\frac{1}{1 - \frac{qg}{\gamma}}\right)^{\alpha/q}, \tag{S56}$$

which is the analytical solution reported in the second line of Eq. 4.47 of Vastola[18] under the assumption $\gamma > q$.

Equivalently, we can write out the PMF:

$$\begin{aligned}\frac{\Gamma(\nu+x)}{x!\Gamma(\nu)}\left(\frac{\nu}{\nu+\nu\theta}\right)^{\nu}\left(\frac{\nu\theta}{\nu+\nu\theta}\right)^{x} &= \frac{(\nu)_x}{x!}\left(\frac{1}{1+\theta}\right)^{\nu}\left(\frac{\theta}{1+\theta}\right)^{x}\\ &= \frac{(\alpha/q)_x}{x!}\left(\frac{1}{1+\frac{q}{\gamma-q}}\right)^{\alpha/q}\left(\frac{\frac{q}{\gamma-q}}{1+\frac{q}{\gamma-q}}\right)^{x}\\ &= \frac{(\alpha/q)_x}{x!}\left(\frac{\gamma-q}{\gamma}\right)^{\alpha/q}\left(\frac{q}{\gamma}\right)^{x},\end{aligned} \tag{S57}$$

which matches the analytical solution reported in the first line of Eq. 4.47 of Vastola[18].

#### S2.2 Autocatalysis with bursty production

Suppose now the 1-species system has bursty production with rate $\alpha$ and mean geometric burst size $b$:

$$\begin{aligned}&\varnothing \xrightarrow{\alpha} B\times\mathcal{X},\\ &\mathcal{X}\xrightarrow{q}\mathcal{X}+\mathcal{X},\\ &\mathcal{X}\xrightarrow{\gamma}\varnothing.\end{aligned} \tag{S58}$$

These reactions yield the following PDE terms:

$$\begin{aligned}\mathcal{M}(u) &= \frac{1}{1-bu}-1,\\ \mathcal{A}(u) &= \alpha\left[\frac{1}{1-bu}-1\right],\\ C &= -\gamma+q,\\ D &= q.\end{aligned} \tag{S59}$$

To solve the system, we first find the solution to the Bernoulli-type differential equation[179], defining $c=-C=\gamma-q$ for convenience:

$$\begin{aligned}\frac{dU}{d\mathsf{s}} &= -cU+qU^2 \text{ such that } U(\mathsf{s}=0)=u \text{ yields}\\ U(\mathsf{s}) &= \frac{cue^{-c\mathsf{s}}}{c+qu(e^{-c\mathsf{s}}-1)}.\end{aligned} \tag{S60}$$

Then the log-generating function of the stationary distribution is given by the integral of $\mathcal{A}(U(\mathsf{s}))$:

$$\begin{aligned}\frac{1}{1-bU}-1 &= \frac{b\frac{cue^{-c\mathsf{s}}}{c+qu(e^{-c\mathsf{s}}-1)}}{1-b\frac{cue^{-c\mathsf{s}}}{c+qu(e^{-c\mathsf{s}}-1)}} = \frac{bcue^{-c\mathsf{s}}}{c+qu(e^{-c\mathsf{s}}-1)-bcue^{-c\mathsf{s}}}\\ &= \frac{bcue^{-c\mathsf{s}}}{(c-qu)-(bc-q)ue^{-c\mathsf{s}}} = \frac{bcu}{c-qu}\frac{e^{-c\mathsf{s}}}{1-\frac{(bc-q)u}{c-qu}e^{-c\mathsf{s}}}\\ \log G &= \int_0^\infty \mathcal{A}(U(\mathsf{s}))d\mathsf{s} = \frac{\alpha bcu}{c-qu}\int_0^\infty \frac{e^{-c\mathsf{s}}}{1-\frac{(bc-q)u}{c-qu}e^{-c\mathsf{s}}}d\mathsf{s}\\ &= -\frac{\alpha b}{bc-q}\log\left(\frac{c-qu-(bc-q)u}{c-qu}\right) = \frac{\alpha b}{bc-q}\log\left(\frac{c-qu}{c-bcu}\right)\\ &= \frac{\alpha b}{bc-q}\log\left(\frac{1-qc^{-1}u}{1-bu}\right) = \frac{\alpha b}{bc-q}\log\left(\frac{b^{-1}-(bc)^{-1}qu}{b^{-1}-u}\right)\\ G &= \left(\frac{b^{-1}-(bc)^{-1}qu}{b^{-1}-u}\right)^{\nu} = \left(\frac{1-qc^{-1}u}{1-bu}\right)^{\nu}, \text{ such that}\\ \nu &= \frac{\alpha b}{bc-q}.\end{aligned} \tag{S61}$$

To achieve a positive $\nu$, we must have

$$\begin{aligned} bc - q &> 0, \\ b(\gamma - q) &> q, \\ b\gamma &> q(1+b), \\ \gamma &> q\frac{1+b}{b}, \end{aligned} \tag{S62}$$

i.e., whereas in the non-bursty case, a steady state was guaranteed by having $\gamma > q$, in the bursty case we must impose a more restrictive condition. The second inequality implies that the coefficient of $u$ in the numerator

$$\frac{q}{bc} = \frac{q}{b(\gamma - q)} < 1; \tag{S63}$$

in other words, $(bc)^{-1}q \in (0,1)$ can be represented by $e^{-\kappa\tau}$ for some positive $\kappa$ and $\tau$. Therefore, the GF

$$G = \left(\frac{b^{-1} - ue^{-\kappa\tau}}{b^{-1} - u}\right)^{\nu} \tag{S64}$$

matches the functional form of the time-dependent MGF of the gamma Ornstein–Uhlenbeck process started at $y = 0$ [166], and the PGF of the bursty transcription/degradation process started at $x = 0$ [16].

Finally, we define $a = e^{-\kappa\tau}$ and rewrite $G$ again:

$$\begin{aligned} G &= \left(\frac{1-abu}{1-bu}\right)^{\nu} = \left(\frac{a(1-abu)}{a(1-bu)}\right)^{\nu} = \left(\frac{a(1-abu)}{1-abu-(1-a))}\right)^{\nu} \\ &= \left(\frac{a}{1-(1-a)\frac{1}{1-abu}}\right)^{\nu} \\ &= G_{\text{NB}}\left(\frac{1}{1-abu}\right) \\ &= \sum_{k=0}^{\infty} P_{\text{NB}}(k)\left(\frac{1}{1-abu}\right)^{k}. \end{aligned} \tag{S65}$$

This is a negative binomial-negative binomial mixture. In other words, the distribution is equivalent to that of a negative binomial distribution with scale parameter $ab$ and stochastic shape parameter $k$, with $k$ in turn drawn from a negative binomial distribution with the shape $\nu$ and success probability $a$. We can confirm this result through a direct calculation:

$$\begin{aligned} P(x) &= \sum_{k=0}^{\infty} P(k)P(x|k), \text{ with PGF} \\ G(g) &= \sum_{x=0}^{\infty} g^x \sum_{k=0}^{\infty} P(k)P(x|k) = \sum_{k=0}^{\infty}\sum_{x=0}^{\infty} g^x P(k)P(x|k), \text{ and assuming NB } P, \\ &= \sum_{k=0}^{\infty}\sum_{x=0}^{\infty} g^x \frac{\Gamma(\nu+k)}{k!\Gamma(\nu)} (a)^{\nu} (1-a)^k \times \frac{\Gamma(k+x)}{x!\Gamma(k)} \left(\frac{k}{k+kab}\right)^k \left(\frac{kab}{k+kab}\right)^x \\ &= \sum_{k=0}^{\infty} \left[\sum_{x=0}^{\infty} \frac{\Gamma(k+x)}{x!\Gamma(k)} \left(\frac{gab}{1+ab}\right)^x\right] \times \frac{\Gamma(\nu+k)}{k!\Gamma(\nu)} (a)^{\nu} (1-a)^k \left(\frac{1}{1+ab}\right)^k \\ &= \sum_{k=0}^{\infty} \left(\frac{1}{1-\frac{gab}{1+ab}}\right)^k \frac{\Gamma(\nu+k)}{k!\Gamma(\nu)} (a)^{\nu} (1-a)^k \left(\frac{1}{1+ab}\right)^k \\ &= \sum_{k=0}^{\infty} \left(\frac{1}{1+ab-gab}\right)^k \frac{\Gamma(\nu+k)}{k!\Gamma(\nu)} (a)^{\nu} (1-a)^k \\ &= \sum_{k=0}^{\infty} \left(\frac{1}{1-abu}\right)^k \frac{\Gamma(\nu+k)}{k!\Gamma(\nu)} (a)^{\nu} (1-a)^k = \sum_{k=0}^{\infty} P_{\text{NB}}(k)\left(\frac{1}{1-abu}\right)^k. \end{aligned} \tag{S66}$$

This expression uses the shape-mean parametrization for the conditional probability $P(x|k)$ and the shape-probability parametrization for the mixing probability $P(k)$ [151].

However, is is considerably easier to notice that the analysis of the Γ-OU model by Sabino and Petroni [166] states that the transient process law is equivalent to that of an Erlang distribution with scale parameter $ab$ and stochastic shape parameter $k$, with $k$ in turn drawn from a negative binomial distribution with the shape $\nu$ and success probability $a$, or scale $\frac{1}{1+a}$. Under Poisson mixing, the Erlang distribution becomes negative binomial.

#### S2.3 RNA and protein catalysis

Suppose we are interested in a 2-species system with $N = 1$, $n = 2$, and $m = 0$, characterized by the following reactions:

$$\begin{aligned} &\mathcal{R} \xrightarrow{\gamma_R} \varnothing, \\ &\mathcal{R} \xrightarrow{q} \mathcal{R} + \mathcal{P}, \\ &\mathcal{P} \xrightarrow{\gamma_P} \varnothing, \end{aligned} \tag{S67}$$

i.e., RNA catalyzes the protein production at rate $q$, and RNA and proteins degrade at rates $\gamma_R$ and $\gamma_P$ respectively. We do not specify the transcriptional dynamics. By plugging in the rates, we obtain the following matrices:

$$\begin{aligned} &C = \begin{bmatrix} -\gamma_R & q \\ 0 & -\gamma_P \end{bmatrix} \text{ and } D = \begin{bmatrix} 0 & q \\ 0 & 0 \end{bmatrix}, \text{ i.e.,} \\ &\frac{\partial G}{\partial t} = \text{transcription} - \gamma_R u_R \frac{\partial G}{\partial u_R} + q u_P (u_R + 1) \frac{\partial G}{\partial u_R} - \gamma_P u_P \frac{\partial G}{\partial u_P}. \end{aligned} \tag{S68}$$

The protein characteristic $U_P = u_p e^{-\gamma_P \mathrm{s}}$ is trivial. The RNA characteristic $U_R$ is described by the following equation:

$$\frac{dU_R}{d\mathrm{s}} = -\gamma_R U_R + q u_P e^{-\gamma_P \mathrm{s}} (U_R + 1), \tag{S69}$$

which has a fairly complicated solution in terms of the incomplete gamma function [77]. Faced with this problem, and drawing on the knowledge that protein abundance is typically much higher than RNA abundance, we may ask whether we can obtain a simpler, approximate form by describing the former by a continuous process (i.e., an $n = 1$ and $m = 1$ system):

$$dy_p = (q x_R - \gamma_P y_P) dt. \tag{S70}$$

Since the $D$ matrix depends on $q = C^{dc}$, $D$ is nonzero and, we immediately find that the discrete-continuous formulation is identical to the original problem.

#### S3 NOTES ON AMBIGUITY

In Section 6.7, we formalized potential ambiguities in the quantification of different transcripts, mentioning two special cases of perfect identifiability and perfect ambiguity. We did not elaborate on this model component, as it is, at this time, less immediately actionable than other components, and requires considerable bioinformatics infrastructure to integrate with analysis. In the current section, we explore this model in more detail.

Even a simple system, as shown in Figure S7a, can contain ambiguity that limits or prevents the identification of transcriptional dynamics. In this illustrative example, we consider a gene with only one intron and two flanking exonic sequences. We suppose that quantification and assignment only consider whether the read overlaps an intron or splice junction. An intron-containing read uniquely identifies the transcript as nascent, whereas a junction-spanning read uniquely identifies the transcript as mature. On the other hand, a fully exonic read does not provide any information about the source molecule. For the sake of completeness, we use “read” as a shorthand for the union of all reads corresponding to a particular UMI: since fragmentation is random, a given UMI will be associated with reads that cover slightly different regions.

The abundance of fully exonic reads depends on the structure and poly(A) content of the source transcript, as well as the sequencing technology. For example, in Figure S7b, we consider reads that can be obtained from a gene that has little to no genomic poly(A) content (red). The unspliced molecules, as well as spliced molecules that have not yet been capped (top), cannot be observed at all. Therefore, their kinetics are not identifiable. On the other hand, the fully mature, poly(A)-tailed transcript (bottom) can be captured at the tail. This capture pattern can give rise to junction reads or formally non-identifiable exonic reads. If the 3’ exon is particularly long relative to fragment length, sequenced fragments will be enriched for purely exonic reads in the 3’ exon. Analogously, if fragment length and the 5’ exon are long relative to read length, sequenced fragments will be enriched for exonic reads in the 5’ exon. In Figure S7c, we illustrate the analogous patterns that can emerge if the unspliced molecule has a single intronic poly(A) region. If the intron and read length are short relative to fragment length, sequencing will produce reads in the 5’ exon.

To characterize transcriptional kinetics, we seek to quantify transient transcripts, which may or may not be mutually identifiable. This is infeasible to optimize on an experimental level. Even in the simple example we provided for illustration, to characterize the source transcript, the transcript region, fragment, and read lengths need to reside in a regime that produces unambiguous reads. To identify the splice junction in Figure S7b, we require fragments that are slightly longer than the 3’ exon and reads that can cover the distance to the junction. However, read length cannot be changed without switching technologies; longer reads typically mean sacrificing the number of sampled cells [180–182]. In the same vein, fragmentation protocols cannot be easily interchanged, as they are optimized for a particular sequencing chemistry. Finally, even if these technical constraints were no object, it would *still* be impossible to optimize for unambiguous capture genome-wide: intron and exon length vary over many orders of magnitude [183,184], and require different read and fragment length regimes for different genes.

Hypothetically, it may be possible to parametrize the $\mathscr{P}^a$ matrix as in Section 6.7, and fit it alongside the biological noise parameters. This approach may not be entirely futile. For example, Equation 73 demonstrates the relevant generating function for two biological species and three identifiable equivalence classes (1: unambiguous nascent, 2: unambiguous mature, 3: ambiguous). The marginal of the nascent species has a functional form distinct from the marginal of the mature species [138], which immediately implies that $\mathscr{P}^a_{1,3} = 0$, $\mathscr{P}^a_{2,3} > 0$ produces distributions functionally distinct from $\mathscr{P}^a_{2,3} = 0$, $\mathscr{P}^a_{1,3} > 0$. In other words, we ought to be able to distinguish the case where all ambiguous counts originate from nascent RNA from the case where they originate from mature RNA, at least in the limit of immaculate and infinite data. We do not, however, expect this approach to be practical for real datasets.

We speculate that it may be more productive to use genomic information to constrain the ambiguity properties, in a similar spirit to Gorin et al. [21] and He et al. [185] For example, if a read aligns to the 3’ untranslated region, *and* we know there is little endogenous poly(A) content 5’ of the read, then we should conclude the read is generated by priming at the poly(A) tail of a capped molecule. In other words, it may be possible to exploit the base information from the genome annotation, the fragment size distributions from orthogonal experiments, and the read size characteristic of the technology to directly construct the $\mathscr{P}^a$ matrix for each transcript.

We illustrate this point in a more quantitative way. Consider the simplest case shown in Figure S7, and further assume that poly(A) capping is rapid. Using the notation in Equation 73, we find that $\mathscr{P}^a_{1,3} = p(\mathscr{f}|1)$, i.e., the probability of sequencing a nascent molecule to obtain a read with an insert from the 5’ exon. Analogously, $\mathscr{P}^a_{2,3} = p(\mathscr{a}|2) + p(\mathscr{e}|2)$, i.e., the probability of sequencing a mature molecule to obtain a read with an insert from the 5’ or 3’ exon. It appears legitimate to propose that these probabilities are only dependent on the sequence and experimental conditions, and may be effectively approximated by exploiting polymer physics or long-read data.

On one hand, this example is somewhat trivial by design. On the other, even this simplified picture of splicing omits important features. First, we presuppose that annotations exist for all downstream transcripts. As we discussed in prior work [16], this is not typically the case, and the identities of and causal relationships between intermediate transcripts are obscure without dedicated study. Second, we presuppose that transcripts can be described as some combination of introns and exons, which transform by the excision of introns. However, even this seemingly reasonable latent assumption ignores elongation, which has been the subject of considerable study elsewhere. For example, a "nascent" transcript may not exist as a physical object, because splicing may complete before the 3' exon is fully transcribed. A more biophysically realistic picture should take into account the fact that splicing occurs during and after elongation. In addition to these biological challenges, various technical effects may need consideration. For example, introns that have already been spliced out may, in principle, be captured and sequenced. If splicing is Markovian, this would be represented as a splitting reaction $\mathcal{X} \to \mathcal{Y} + \mathcal{Z}$, which is not immediately tractable using our framework. Finally, due to a variety of artifacts, the reads themselves may have a more complex relationship to the source transcripts. These effects include strand invasion and aberrant priming, and may lead to reads containing antisense and template switch oligo sequences[163]. Formally, all of these effects can be integrated into a sufficiently complicated stochastic model. In practice, we recommend introducing complexity only when simpler models fail.

#### S4 NOTES ON REGULATION

In the current section, we outline one notable limitation of the generating function approach we have presented. For essentially any system one could define, we could proceed as in the first few steps of our approach: one could write down a master equation, define a generating function, and convert the master equation into a PDE (or system of PDEs) satisfied by the generating function. But our approach relies on the fact that, for the systems we consider, the PDEs one obtains can be solved using the method of characteristics; this breaks down for systems where this is not the case.

Notable examples of systems where the method of characteristics is insufficient include ones involving feedback, like systems with proteins that can bind to their gene's promoter to influence that gene's transcription rate. In cases like this, the method breaks down for a straightforward reason. In this section, we will attempt to illustrate this breakdown in the context of a simple and concrete example.

Consider a system with one discrete species and two possible production states, and assume that it features feedback analogous to protein-promoter binding:

$$\mathcal{S}_{\text{off}} + \mathcal{X} \underset{k_{\text{off}}}{\overset{k_{\text{on}}}{\rightleftarrows}} \mathcal{S}_{\text{on}}, \ \mathcal{S}_{\text{on}} \xrightarrow{k_{\text{init}}} \mathcal{S}_{\text{on}} + \mathcal{X}, \ \mathcal{X} \xrightarrow{\gamma} \varnothing. \tag{S71}$$

One could interpret the discrete species as representing either some sort of microRNA that binds to its own gene's promoter, or a protein that does (in which case, we assume RNA dynamics can be ignored). But the plausibility or interpretation of this model is not particularly important for what follows, given that the main point is how feedback makes even simple models like this not amenable to our primary technique.

The corresponding master equation is

$$\begin{aligned}
\frac{\partial P_{\text{on}}}{\partial t} &= -k_{\text{off}} P_{\text{on}}(x-1,t) + k_{\text{on}}(x+1) P_{\text{off}}(x+1,t) \\
&\quad + k_{\text{init}} \left[ P_{\text{on}}(x-1,t) - P_{\text{on}}(x,t) \right] \\
&\quad + \gamma \left[ (x+1) P_{\text{on}}(x+1,t) - x P_{\text{on}}(x,t) \right] \\
\frac{\partial P_{\text{off}}}{\partial t} &= k_{\text{off}} P_{\text{on}}(x-1,t) - k_{\text{on}}(x+1) P_{\text{off}}(x+1,t) \\
&\quad + \gamma \left[ (x+1) P_{\text{off}}(x+1,t) - x P_{\text{off}}(x,t) \right],
\end{aligned} \tag{S72}$$

and we have a generating function $\mathbf{G} = (G_{\text{on}}, G_{\text{off}})^T$ with

$$\begin{aligned}
G_{\text{on}}(g,t) &:= \sum_{x=0}^{\infty} g^x \, P_{\text{on}}(x,t) \\
G_{\text{off}}(g,t) &:= \sum_{x=0}^{\infty} g^x \, P_{\text{off}}(x,t).
\end{aligned} \tag{S73}$$

From the master equation, we obtain PDEs

$$\begin{aligned}
\frac{\partial G_{\text{on}}}{\partial t} &= -k_{\text{off}} g \, G_{\text{on}} + k_{\text{on}} \frac{\partial G_{\text{off}}}{\partial g} + k_{\text{init}}(g-1) G_{\text{on}} - \gamma(g-1) \frac{\partial G_{\text{on}}}{\partial g} \\
\frac{\partial G_{\text{off}}}{\partial t} &= k_{\text{off}} g \, G_{\text{on}} - k_{\text{on}} \frac{\partial G_{\text{off}}}{\partial g} - \gamma(g-1) \frac{\partial G_{\text{off}}}{\partial g}.
\end{aligned} \tag{S74}$$

Normally, to apply the method of characteristics, we move all terms that depend on generating function derivatives to the left-hand side, and keep all other terms on the right-hand side. We would rearrange the terms of this particular set of equations so that

$$\begin{aligned}
\frac{\partial G_{\text{on}}}{\partial t} + \gamma(g-1) \frac{\partial G_{\text{on}}}{\partial g} - k_{\text{on}} \frac{\partial G_{\text{off}}}{\partial g} &= -k_{\text{off}} g \, G_{\text{on}} + k_{\text{init}}(g-1) G_{\text{on}} \\
\frac{\partial G_{\text{off}}}{\partial t} + \gamma(g-1) \frac{\partial G_{\text{off}}}{\partial g} + k_{\text{on}} \frac{\partial G_{\text{off}}}{\partial g} &= k_{\text{off}} g \, G_{\text{on}}.
\end{aligned} \tag{S75}$$

Next, we would try to argue that there exists a solution to the above equation along some one-dimensional curve parameterized by a variable $\mathsf{s}$. But we have the following problem: the left-hand side mixes $G_{\text{on}}$ and $G_{\text{off}}$ is a nontrivial way. This is something that never happens in the class of systems we consider in the rest of the paper, since state switching is "upstream" of every other possible reaction (i.e., terms which mix $G_{\text{on}}$ and $G_{\text{off}}$ only appear on the right-hand side, where they cannot affect the applicability of the method of characteristics).

Is there a way to separate $G_{\text{on}}$ and $G_{\text{off}}$, so that we can rescue our approach? This is somewhat unclear, but the answer is probably *no*. Independent variables are often determined through some kind of diagonalization procedure, and this is a strategy which does not work here. Define the matrix

$$M := \begin{bmatrix} \gamma(g-1) & -k_{\text{on}} \\ 0 & \gamma(g-1)+k_{\text{on}} \end{bmatrix}, \tag{S76}$$

so that we can write the left-hand side in vector form as

$$\frac{\partial \mathbf{G}}{\partial t} + M \frac{\partial \mathbf{G}}{\partial g}. \tag{S77}$$

In principle, we could separate the variables by diagonalizing $M$ and working in terms of its eigenvectors. Here, because $M$ is a $2 \times 2$ matrix, we can write down explicitly that

$$\mathbf{v}_0 = (1,0)^T \qquad M\mathbf{v}_0 = \gamma(g-1)\mathbf{v}_0, \tag{S78}$$

$$\mathbf{v}_1 = (1,-1)^T \qquad M\mathbf{v}_1 = [\gamma(g-1)+k_{\text{on}}]\,\mathbf{v}_1. \tag{S79}$$

Most importantly, the eigenvalues of $M$ are distinct as long as $k_{\text{on}} > 0$—which must be true for our problem to have nontrivial switching dynamics. We can write down a decomposition $M = Q^{-1}\Lambda Q$, define $\tilde{\mathbf{G}} := Q\mathbf{G}$, and convert the left-hand side of our PDE to

$$\frac{\partial \tilde{\mathbf{G}}}{\partial t} + \Lambda \frac{\partial \tilde{\mathbf{G}}}{\partial g}. \tag{S80}$$

Here is where our attempt to use the method of characteristics fails. When we unpack this expression, we have components

$$\begin{aligned} &\frac{\partial \tilde{G}_0}{\partial t} + \lambda_0 \frac{\partial \tilde{G}_0}{\partial u} \\ &\frac{\partial \tilde{G}_1}{\partial t} + \lambda_1 \frac{\partial \tilde{G}_1}{\partial u}, \end{aligned} \tag{S81}$$

where we have converted from $g$ to $u := g-1$ to match our arguments elsewhere. We would like there to be a solution to our PDE where $G$ depends on $u$ through a parameterized curve $U(\mathsf{s})$ which has $U(\mathsf{s}=0) = u$, and not in a more arbitrary way. In particular, it should be true that

$$\begin{aligned} \frac{d\tilde{G}_0}{d\mathsf{s}} &= \frac{\partial \tilde{G}_0}{\partial t}\frac{dt}{d\mathsf{s}} + \frac{\partial \tilde{G}_0}{\partial U}\frac{dU}{d\mathsf{s}} = \frac{\partial \tilde{G}_0}{\partial t} + \lambda_0 \frac{\partial \tilde{G}_0}{\partial u}, \\ \frac{d\tilde{G}_1}{d\mathsf{s}} &= \frac{\partial \tilde{G}_1}{\partial t}\frac{dt}{d\mathsf{s}} + \frac{\partial \tilde{G}_1}{\partial U}\frac{dU}{d\mathsf{s}} = \frac{\partial \tilde{G}_1}{\partial t} + \lambda_1 \frac{\partial \tilde{G}_1}{\partial u}. \end{aligned} \tag{S82}$$

But this is impossible, since we obtain two incompatible constraints on the characteristic $U$. It must be simultaneously true that

$$\begin{aligned} &\frac{dU}{d\mathsf{s}} = \gamma U \qquad \text{and} \\ &\frac{dU}{d\mathsf{s}} = \gamma U + k_{\text{on}}, \end{aligned} \tag{S83}$$

which is only possible if $k_{\text{on}} = 0$.

This problem is generic, and not particular to this simple example, since a diagonalization strategy on a problem with more states would again yield incompatible constraints unless the analogue to $M$ were proportional to the identity matrix (i.e., there is no feedback). This functional form is certainly precedented in the study of partial differential equations (see, e.g., Section 22.4 of the Ebert and Reissig textbook[98] and Section 2.5 of the John textbook[186]), but does not appear to produce analytically tractable solutions.

One might expect there to be some appropriate generalization of the method of characteristics that works for problems like this. While there exist other strategies to solve very specific problems that involve feedback[101–104], we are not aware of a generalization of the method of characteristics that permits a strategy similar to the one we rely on throughout the main text. There may be deep reasons for this; systems with feedback are capable of substantially more complex behavior, and are likely to resist the simple descriptions we can provide for the essentially "feedforward" systems we focus on.

### S5 SUPPLEMENTARY FIGURES

#### S5.1 Empty droplets

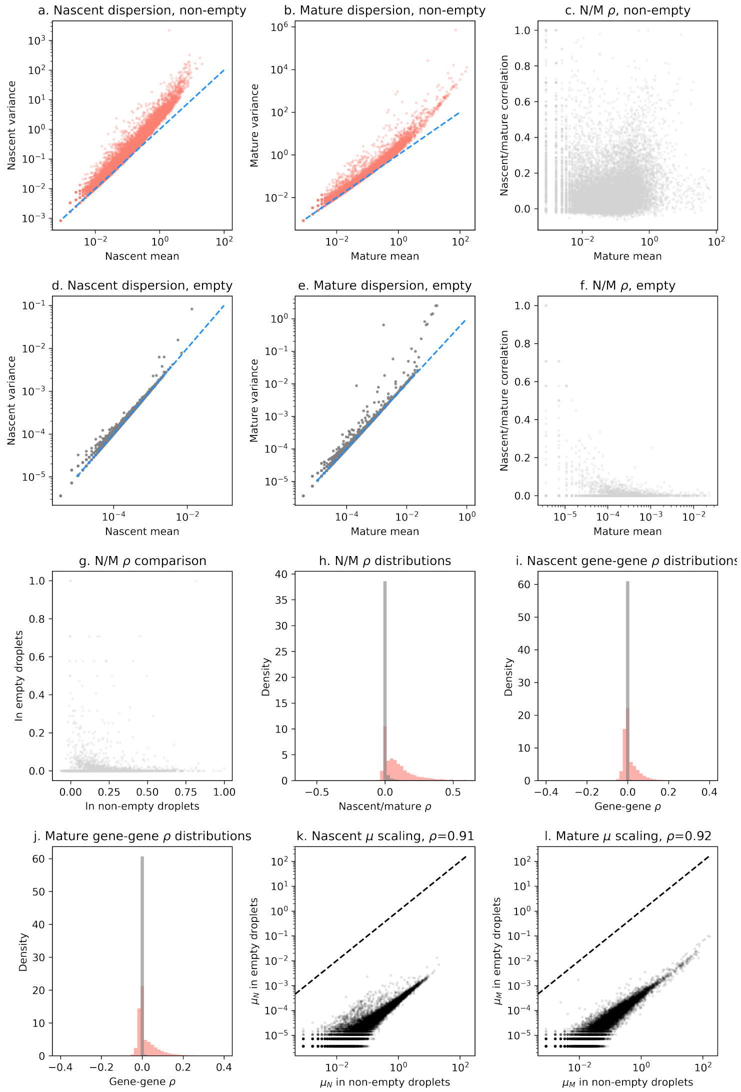


**Figure S1** The statistical properties of empty and non-empty droplets, computed using the `pbmc_1k_v3` dataset.
**a.** Nascent RNA counts in non-empty drops are conspicuously overdispersed (points: genes; dashed line: Poisson identity line; genes with zero mean or variance omitted).
**b.** As in **a**, considering mature RNA counts.
**c.** Nascent and mature RNA counts in non-empty droplets show a considerable range of correlations (points: genes; genes with zero mean or undefined correlation omitted).
**d.** Nascent RNA counts in empty drops exhibit dispersions close to the Poisson limit (conventions as in **a**).
**e.** As in **f**, considering mature RNA counts.
**f.** Nascent and mature RNA counts in empty droplets typically have low correlations (conventions as in **c**).
**g.** Nascent/mature count correlations in empty and non-empty droplets do not appear to show a statistical relationship (points: genes; genes with undefined correlation omitted).
**h.** Marginal histograms of panels **c** and **f**: empty droplets overwhelmingly display nascent/mature correlations near zero, whereas non-empty droplets show nontrivial correlations (red histogram: values in non-empty droplets; gray histogram: values in empty droplets; undefined correlation values omitted). **i.** Empty droplets overwhelmingly demonstrate gene-gene correlations for nascent counts near zero, whereas non-empty droplets show nontrivial correlations (conventions as in **h**).
**j.** As in **i**, considering mature RNA counts.
**k.** Average nascent RNA content in empty droplets is highly correlated with the convent of non-empty droplets, but lowered by several orders of magnitude (points: genes; genes with zero mean omitted).
**l.** As in **k**, considering mature RNA counts.

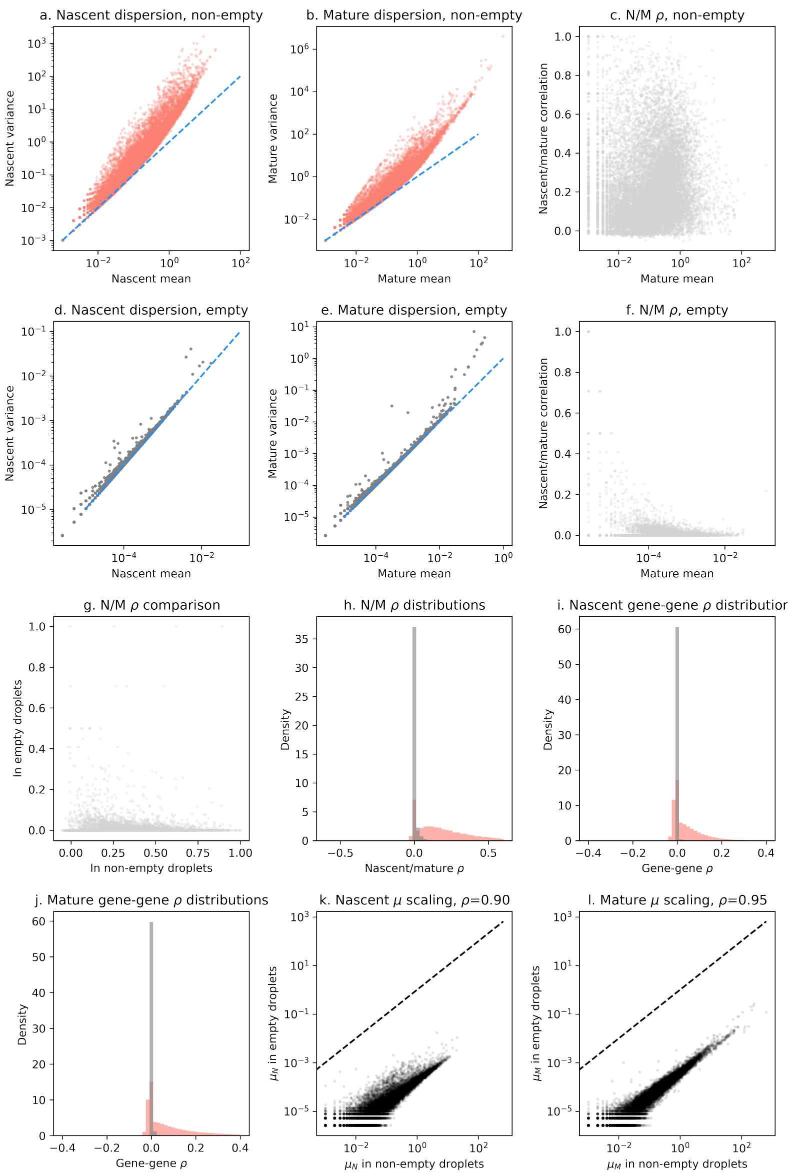


**Figure S2** The statistical properties of empty and non-empty droplets, computed using the `heart_1k_v3` dataset. All conventions as in Figure S1.

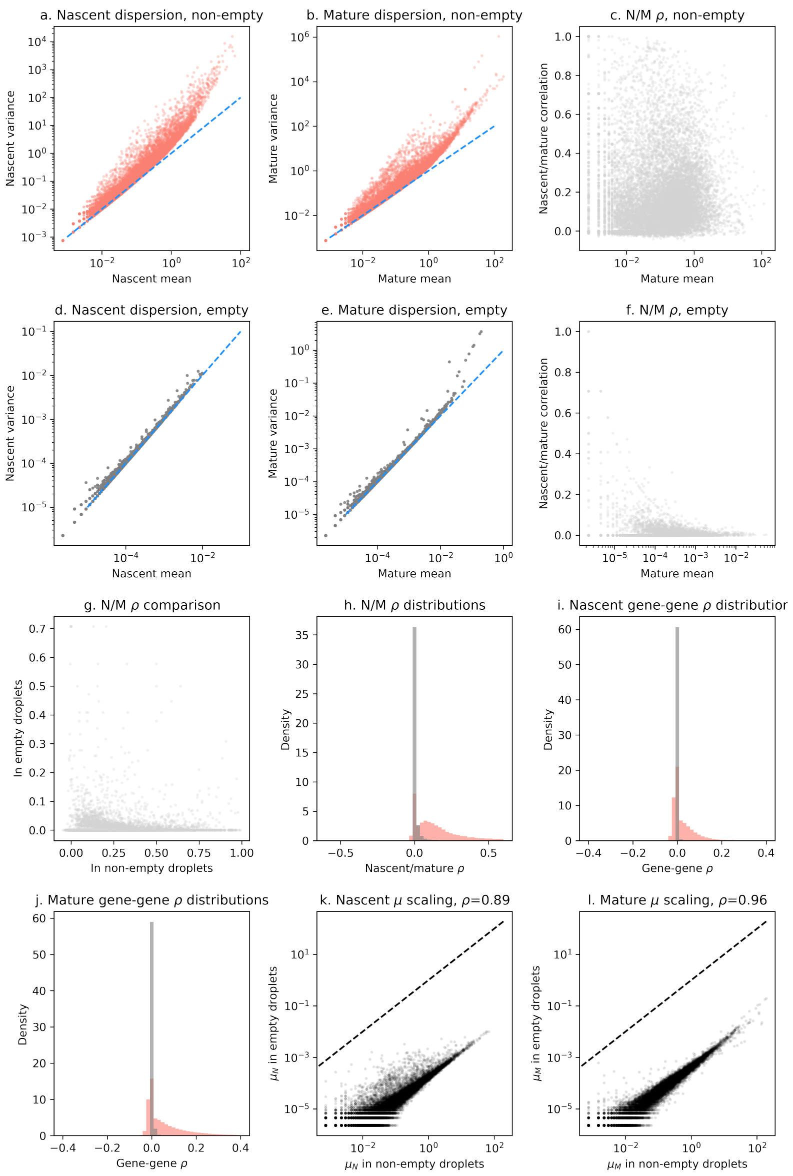


**Figure S3** The statistical properties of empty and non-empty droplets, computed using the `neuron_1k_v3` dataset. All conventions as in Figure S1.

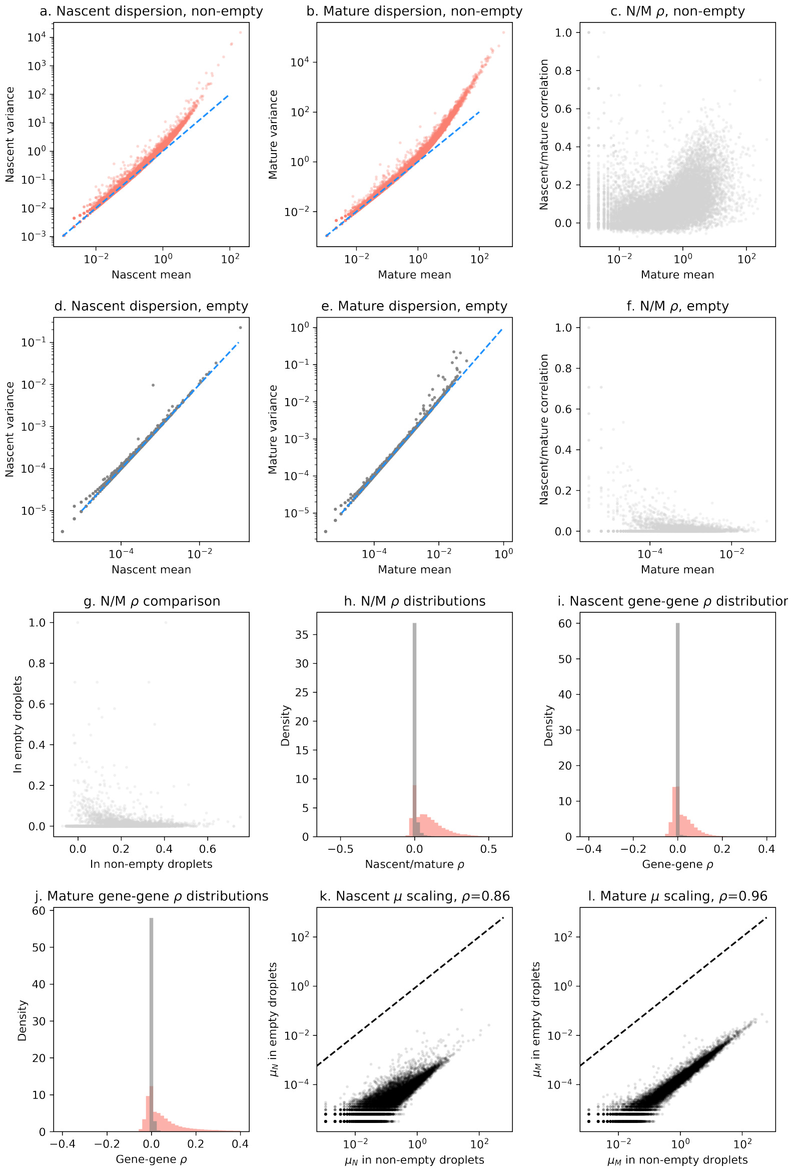


**Figure S4** The statistical properties of empty and non-empty droplets, computed using the `desai_dmso` dataset. All conventions as in Figure S1.

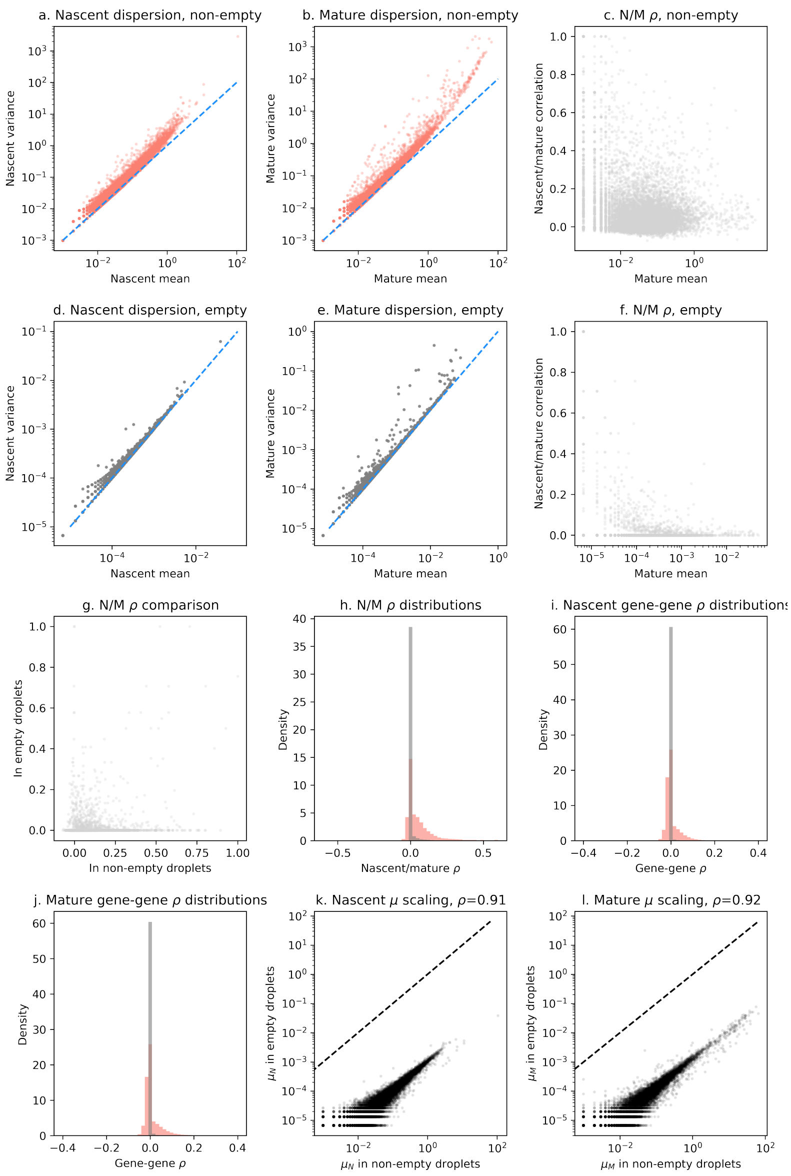


**Figure S5** The statistical properties of empty and non-empty droplets, computed using the `pbmc_1k_v2` dataset. All conventions as in Figure S1.

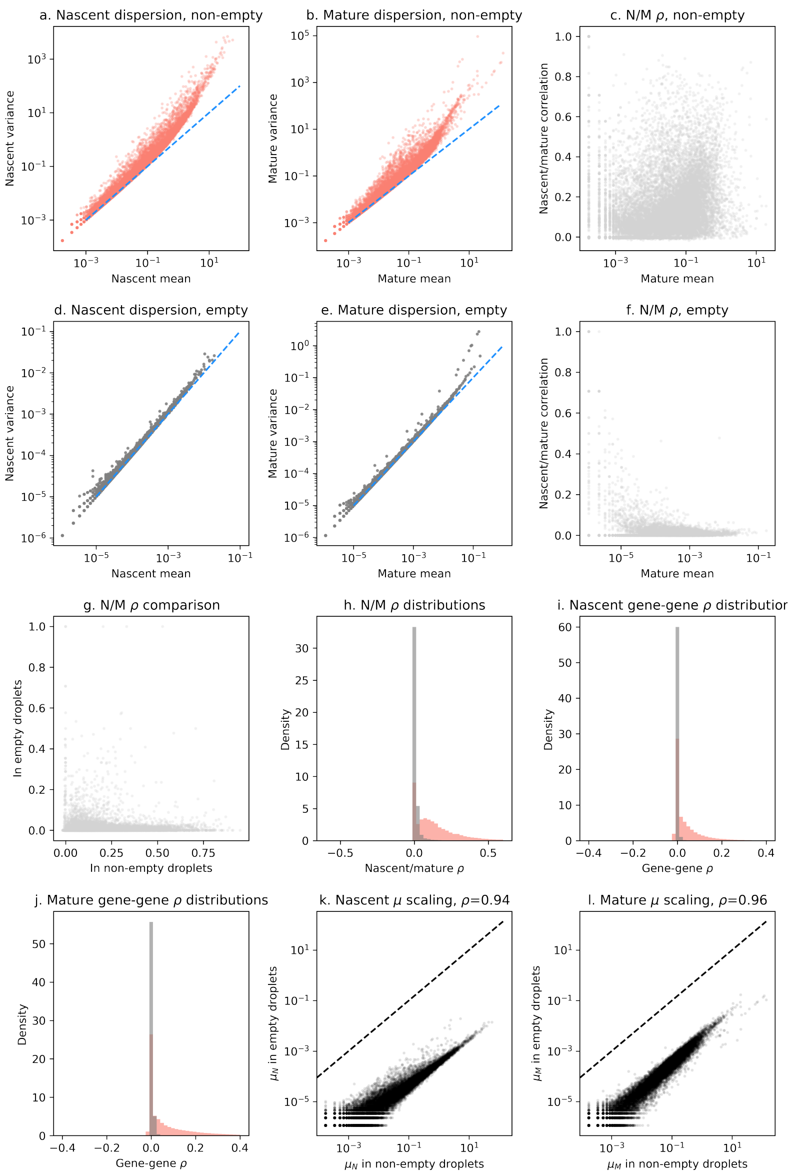


**Figure S6** The statistical properties of empty and non-empty droplets, computed using the `brain_nuc_5k_v3` dataset. All conventions as in Figure S6.

#### S5.2 Read ambiguity

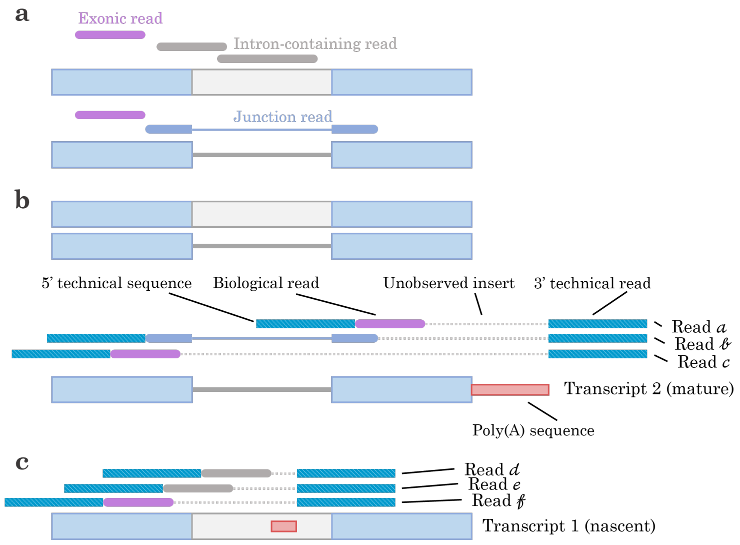


**Figure S7** Potential sources of short-read sequencing ambiguity in a hypothetical one-intron, two-exon transcript.
**a.** Possible splicing information conveyed by reads in the hypothetical transcript (magenta: reads that only contain exonic information; dark gray: reads that contain intronic information; dark blue: reads that overlap a splice junction. Blue block: exon; gray block: present intron; line: excised intron. 5' end is toward the left).
**b.** Categories of reads that can be obtained by sequencing the transcript, assuming no endogenous poly(A) content (cyan block: technical reads and indices; dotted lines: residual inserts not observed by sequencing; red block: poly(A) sequence).
**c.** Categories of reads that can be obtained by capturing a transcript at an endogenous, intronic poly(A) sequence (conventions as in **a** and **b**).
